# Model metamers illuminate divergences between biological and artificial neural networks

**DOI:** 10.1101/2022.05.19.492678

**Authors:** Jenelle Feather, Guillaume Leclerc, Aleksander Mądry, Josh H. McDermott

**Author notes:** Corresponding author jenellefeather.com.

## Abstract

Deep neural network models of sensory systems are often proposed to learn representational transformations with invariances like those in the brain. To reveal these invariances we generated “model metamers” – stimuli whose activations within a model stage are matched to those of a natural stimulus. Metamers for state-of-the-art supervised and unsupervised neural network models of vision and audition were often completely unrecognizable to humans when generated from deep model stages, suggesting differences between model and human invariances. Targeted model changes improved human-recognizability of model metamers, but did not eliminate the overall human-model discrepancy. The human-recognizability of a model’s metamers was well predicted by their recognizability by other models, suggesting that models learn idiosyncratic invariances in addition to those required by the task. Metamer recognition dissociated from both traditional brain-based benchmarks and adversarial vulnerability, revealing a distinct failure mode of existing sensory models and providing a complementary benchmark for model assessment.

## Introduction

A central goal of neuroscience is to build models that reproduce brain responses and behavior. The hierarchical nature of biological sensory systems (1, 2) has motivated the use of hierarchical neural network models that transform sensory inputs into task-relevant representations (3, 4). As such models have become the top-performing machine perception systems over the last decade, they have also emerged as the leading models of both the visual and auditory systems (5–11).

One hypothesis for why artificial neural network models might replicate computations found in biological sensory systems is that they instantiate invariances that mirror those in such systems (12–14). For instance, visual object recognition must often be invariant to pose, and to the direction of illumination. Similarly, speech recognition must be invariant to speaker identity and to details of the prosodic contour. Sensory systems are hypothesized to build up invariances (15–17) that enable robust recognition. Such invariances plausibly arise in neural network models as a consequence of optimization for recognition tasks or other training objectives.

Although biological and artificial neural networks might be supposed to have similar internal invariances, there are some known human-model discrepancies that suggest the invariances of the two systems do not perfectly match. For instance, model judgments are often impaired by stimulus manipulations to which human judgments are invariant, such as additive noise (18–20) or small translations of the input (21, 22). Another such discrepancy is the vulnerability to adversarial perturbations – small changes to stimuli that alter model decisions despite being imperceptible to humans (23–26). These findings illustrate that current task-optimized models lack some of the invariances of human perception, but leave many questions unresolved. For instance, because the established discrepancies rely on only the model’s output decisions, they do not reveal where in the model the discrepancies arise. It also remains unclear whether observed discrepancies are specific to supervised learning procedures that are known to deviate from biological learning. And because the known discrepancies do not point to a general method to assess model invariances in the absence of a specific hypothesis, it remains possible that current models possess many other invariances that humans lack.

In this paper we present a general test of whether the invariances present in computational models of the auditory and visual systems are also present in human perception, and apply this test to a set of contemporary and classical models. Rather than target particular known human invariances, we visualize or sonify model invariances by synthesizing stimuli that produce approximately the same activations in a model. We draw inspiration from human perceptual metamers, which have previously been characterized in the domains of color perception (27, 28), texture (29–31), cue combination (32), Bayesian decision making (33), and visual crowding (34, 35). We call the stimuli we generate “model metamers” because they are metameric for a computational model^1^.

We generated model metamers from a variety of state-of-the-art deep neural network models of vision and audition by synthesizing stimuli that yield the same activations in a model stage as particular natural images or speech signals. We then evaluated human recognition of the model metamers. If the model invariances match those of humans, then humans should be able to recognize the model metamer as belonging to the same class as the natural signal to which it is matched.

Across both visual and auditory task-optimized neural networks, metamers from late model stages were nearly always misclassified by humans, suggesting that many of their invariances are not present in human sensory systems. The same phenomenon also occurred for networks trained with unsupervised learning, demonstrating that the model failure is not specific to supervised classifiers. Model metamers could be made more recognizable to humans with selective changes to the training procedure or architecture. However, late-stage model metamers remained much less recognizable than natural stimuli in every model we tested regardless of architecture or training. Some model changes that produced more recognizable metamers did not improve conventional neural prediction metrics or evaluations of robustness, demonstrating that the metamer test provides a complementary tool to guide model improvements. The human-recognizability of a model’s metamers was well predicted by other models’ recognition of the same metamers, suggesting that the discrepancy with humans lies in idiosyncratic model-specific invariances. Model metamers demonstrate a qualitative gap that remains between current models of sensory systems and their biological counterparts, and provide a metric for future model evaluation.

## Results

### General procedure

The goal of our metamer generation procedure (Figure 1a) was to generate stimuli that produce nearly identical activations at some stage within a model, but that were otherwise unconstrained, and thus could physically differ in ways to which the model was invariant. We first measured the activations evoked by a natural image or speech signal at a particular model stage. The metamer for the natural image or speech signal was then initialized as a white noise signal (either an image or a sound waveform; white noise was chosen to sample the metamers as broadly as possible subject to the model constraints, without biasing the initialization towards a specific object class). The noise signal was then modified to minimize the difference between its activations at the model stage of interest and those for the natural signal to which it was matched. The optimization procedure performed gradient descent on the input, iteratively updating the input while holding the model parameters fixed. Model metamers can be generated in this way for any model stage constructed from differentiable operations. Because the models we considered are hierarchical, if the image or sound was matched with high fidelity at a particular stage, all subsequent stages were also matched (including the final classification stage in the case of supervised models, yielding the same decision).

**Figure 1.**
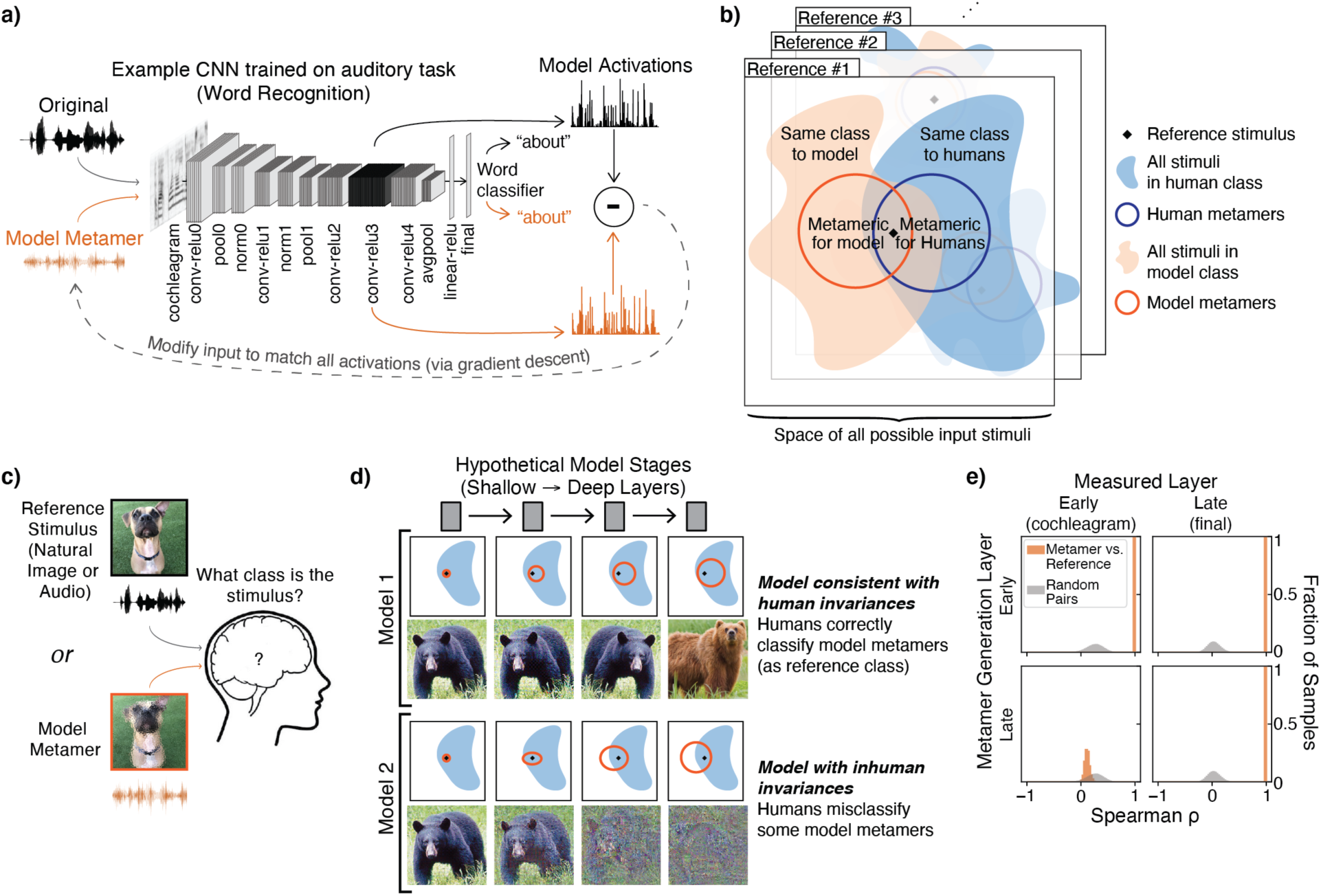
a) Model metamer generation. Metamers are synthesized by performing gradient descent on a noise signal to minimize the difference between its activations at a model stage and those of a natural signal. The model architecture shown here is that of the CochCNN9 auditory model class used throughout the paper. b) Each reference stimulus has an associated set of stimuli in the space of all possible stimuli that are categorized as the same class by humans (blue) or by models (orange, if models have a classification decision). Metamers for humans and metamers for models are also sets of stimuli in the space of all possible stimuli (subsets of the set of same-class stimuli). In this paper, model metamers are derived for a specific model stage, taking advantage of access to the internal representations of the model at each stage. Because we do not similarly have access to the internal representations of humans, we evaluate the human-consistency of a model’s metamers with a behavioral classification judgment. c) General experimental setup. Humans make classification judgements on natural stimuli or model metamers. d) Possible scenarios for how model metamers could relate to human classification decisions. Each square panel depicts sets of stimuli in the input space. The top row depicts a model that passes our proposed behavioral test. The set of metamers for a reference stimulus grows over the course of the model, but even at the deepest stage, all model metamers are classified as the reference category by humans. The second row depicts a model whose invariances diverge from those of humans. By the late stages of the model, many model metamers are no longer recognizable by humans as the reference stimulus class. The metamer test results also constrain the model stage at which model invariances diverge from those of humans. e) Example distributions of activation similarity for pairs of metamers (a natural signal and its corresponding metamer) along with random pairs of natural signals from the training set. The latter provides a null distribution that we used to verify the success of the model metamer generation. The correlation between metamer activations in the model stage used for metamer generation (top left and bottom right) falls outside the null distribution, as intended. When the metamer is generated from an early stage, the activations in a deeper stage are also well matched (top right), as expected given that the model is feedforward and deterministic. By contrast, when metamers are generated from a deep stage, activations in the deep stage are well matched, but those in the early stage are not (bottom left). This is because the model builds up invariances, such that metamers for the deep stage produce very different activations in an early stage.

### Experimental logic

The logic of our approach can be related to four sets of stimuli. For a given “reference” stimulus, there is a set of stimuli for which humans produce the same classification judgment as the reference (Figure 1b; blue shaded). A subset of these are stimuli that are indistinguishable from the reference stimulus (i.e., metameric) to human observers (blue circle). If a model performs a classification task, it will also have a set of stimuli judged to be the same category as the reference stimulus (orange shaded). However, even if the model does not itself perform classification, it could instantiate invariances that define model metamers for the reference stimulus at each model stage (orange circle).

In our experiments, we generate stimuli (sounds or images) that are metameric to a model and present these stimuli to humans performing a classification task (Figure 1c). Because we have access to the internal representations of the model, we can generate metamers for each stage of a model (i.e. sampling from the preimage of the reference stimulus activations at each model stage; Figure 1d). In many models there is limited invariance in the early stages (as is believed to be true of early stages of biological sensory systems (17)), with model metamers closely approximating the stimulus from which they are generated (Figure 1d, small orange sets in leftmost column). But successive stages of a model may build up invariance, producing successively larger sets of model metamers. In a feedforward model, if two distinct inputs map onto the same representation at a given model stage, then any differences in the inputs cannot be recovered in subsequent stages, such that invariance cannot decrease from one stage to the next. If a model replicates a human sensory system, every model metamer from each stage should also be classified as the reference class by human observers (first row of Figure 1d). Such a result does not imply that all human invariances will be shared by the model, but it is a necessary condition for a model to replicate human invariances.

Discrepancies in human and model invariances could result in model metamers that are not recognizable by human observers (second row of Figure 1d). Assessing model metamers from different model stages may provide insight into how the invariances are built up over the model, and where any discrepancies with humans arise within the model.

Our approach differs from classical work on metamers (28) in that we do not directly assess whether model metamers are also metamers for human observers (i.e., indistinguishable). The reason for this is that a human judgment of whether two stimuli are the same or different could rely on any representations within their sensory system that distinguish the stimuli, rather than just those that are relevant to a particular behavior. By contrast, most neural network models of sensory systems are trained to perform a single behavioral task. As a result, we do not expect metamers of such models to be fully indistinguishable to a human observer. But if the model replicates the representations that support a particular behavioral task in humans, its metamers should nonetheless produce the same human behavioral judgment on that task, because they should be indistinguishable to the human representations that mediate the judgment. We thus use a recognition judgment as the behavioral assay of whether model metamers reflect the same invariances that are instantiated in an associated human sensory system. Moreover, if humans cannot recognize a model metamer, they would also be able to discriminate it from the reference stimulus, and the model would also not pass a traditional metamerism test.

We sought to answer several questions. First, we asked whether the learned invariances of commonly used neural network models are shared by human sensory systems. Second, we asked where any discrepancies with human perception arise within models. Third, we asked whether any discrepancies between model and human invariances would also be present in models obtained without supervised learning. Fourth, we explored whether model modifications intended to improve robustness would also make model metamers more recognizable to humans. Fifth, we asked whether metamer recognition identifies model discrepancies that are not evident using other methods of model assessment, such as brain predictions or adversarial vulnerability. Sixth, we asked whether human-discrepant metamers are shared across models.

### Metamer optimization

Because the metamer generation relies on an iterative optimization procedure, it was important to measure optimization success for each metamer. We considered the metamer generation to have succeeded only if it satisfied two conditions. First, measures of the match between the activations for the natural reference stimulus and its model metamer at the matched stage had to be much higher than would be expected by chance, quantified with a null distribution (Figure 1e, grey distributions) measured between randomly chosen pairs of examples from the training dataset. This criterion was adopted in part because it is equally applicable to models that do not perform a task. Metamers had to pass this criterion for each of three different measures of the match (Pearson and Spearman correlations, and dB SNR; see Methods). Second, for models that performed a classification task, the metamer had to result in the same classification decision by the model as the paired natural signal. In practice we trained linear classifiers on top of all unsupervised models, such that we were able to apply this second criterion for them as well (to be conservative).

Figure 1e shows example distributions of the match fidelity (using Spearman’s rho in this example). Activations in the matched model stage have a correlation close to 1, as intended, and are well outside the null distribution for random pairs of training examples. And as expected given the feedforward nature of the model, matching at an early stage produces matched activations in a late stage (Figure 1e, orange distributions, top row). But because the models we consider build up invariances over a series of feedforward stages, stages earlier than the matched stage need not have the same activations, and in general differ from those for the original stimulus to which the metamer was matched (Figure 1e, orange distributions, bottom row). This is particularly true for model metamers generated from late stages of the model, where many different signals can yield the same model activations. The match fidelity of this example was typical, and optimization summaries for each analyzed model are included at https://github.com/jenellefeather/model_metamers_pytorch.

### Metamers for standard visual deep neural networks are unrecognizable to humans

We generated model metamers for multiple stages of five standard visual neural networks trained to recognize objects (using the ImageNet1K dataset (37)) (Figure 2a). The five network models spanned a wide range of architectural building blocks and depths: CORnet-S (38), VGG-19 (39), ResNet50 and ResNet101 (40), and AlexNet (41). These models have been posited to capture similar features as primate visual representations, and at the time the experiments were run, they respectively placed 1^st^, 2^nd^, 11^th^, 4^th^, and 59^th^ on a neural prediction benchmark (38, 42). We subsequently ran a second experiment on an additional five models pretrained on larger scale datasets that became available at later stages of the project (ResNet50:CLIP and ViT-B_32:CLIP (43), ResNet50:SWSL and ResNeXt101-32×8d:SWSL (44), ViT_large_patch-16_224 (45)). To evaluate human recognition of the model metamers, humans performed a 16-way categorization task on the natural stimuli and model metamers (Figure 2b) (18). In models with residual connections, we only generated metamers at stages where all branches converge, which ensured that all subsequent model stages, and the model decision, remained matched.

**Figure 2.**
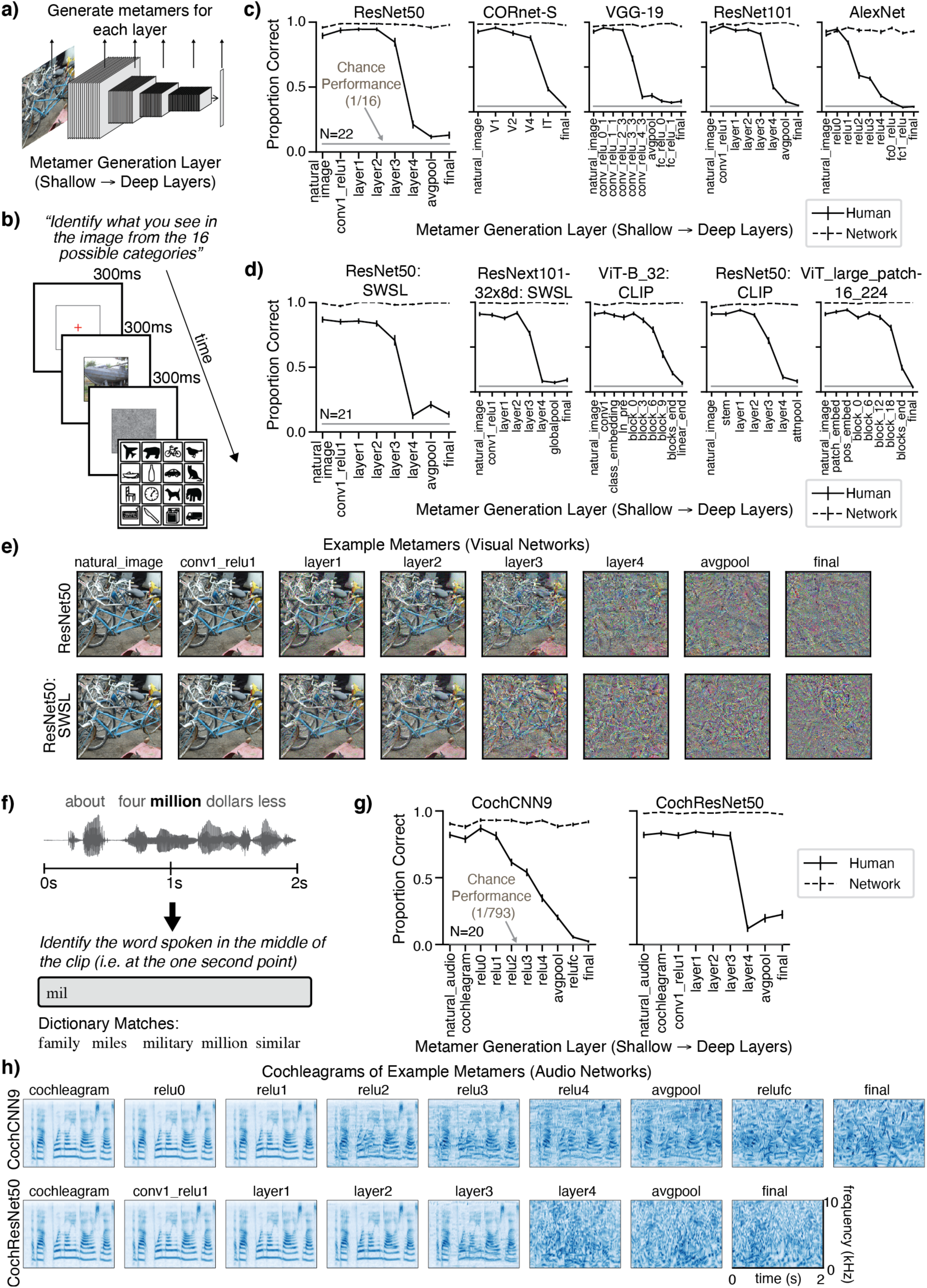
*(previous page)*. Metamers of standard-trained visual and auditory deep neural networks are often unrecognizable to human observers. a) Model metamers are generated from different stages of the model. b) Experimental task used to assess human recognition of visual model metamers. Humans were presented with an image (either a natural image or a model metamer of a natural image) followed by a noise mask. They were then presented with 16 icons representing 16 object categories, and classified each image as belonging to one of these 16 categories by clicking on the icon. c) Human recognition of image model metamers (N=22). For all tested models, human recognition of model metamers declined for late model stages, while model metamers remained recognizable to the model (as expected). Error bars plot SEM across participants (or participant matched stimulus subsets for model curves) d) Human recognition of image model metamers (N=21) trained on larger scale datasets. e) Example metamers from ResNet50 and ResNet50: CLIP trained visual models. f) Experimental task used to assess human recognition of auditory model metamers. Humans classified the word that was present at the midpoint of a two-second sound clip. Participants selected from 793 possible words by typing any part of the word into a response box and seeing matching dictionary entries from which to complete their response. A response could only be submitted if it matched an entry in the dictionary. g) Human recognition of auditory model metamers (N=20). For both tested models, human recognition of model metamers decreased at late model stages. By contrast, model metamers remain recognizable to the auditory models, as expected. When plotted, chance performance (1/793) is indistinguishable from the x-axis. Error bars plot SEM across participants (or participant matched stimulus subsets for model curves). h) Cochleagram visualizations of example auditory model metamers from CochCNN9 and CochResNet50 architectures. Color intensity denotes instantaneous sound amplitude in a frequency channel (arbitrary units)

Contrary to the idea that the trained neural networks have learned human-like invariances, we found that human recognition of the model metamers decreased across model stages, reaching near-chance performance at the deepest stages even though the model metamers remained as recognizable to the models as the corresponding natural stimuli, as intended (Figure 2c&d). This reduction in human recognition of model metamers was evident as a main effect of observer and an interaction between the effect of metamer generation stage and the observer, both of which were statistically significant for each of the ten models (main effects, CORnet-S: F(1, 42)=424.4, p<0.0001, η^2^*p* =0.91; VGG-19: F(1, 42)=1612.1, p<0.0001, η^2^*p* =0.97; ResNet50: F(1, 42)=554.9, p<0.0001, η^2^*p* =0.93; ResNet101: F(1, 42)=1001.7, p<0.0001, η^2^*p* =0.96; AlexNet: F(1, 42)=935.4, p<0.0001, η^2^*p* =0.96; ResNet50-CLIP: F(1, 40)= 517.3, p<0.0001, η^2^*p=*0.93, ViT-B_32-CLIP: F(1, 40)= 604.5, p<0.0001, η^2^*p*=0.94, ResNet50-SWSL: F(1, 40)= 1159.3, p<0.0001, η^2^*p*=0.97, ResNeXt101-32×3d-SWSL: F(1, 40)=2363.1, p<0.0001, η^2^*p*=0.98, ViT_large_patch-16_224: F(1, 40)=852.01 p<0.0001, η^2^*p*=0.96; interactions, CORnet-S: F(5, 210)=293.6, p<0.0001, η^2^*p* =0.87; VGG-19: F(9, 378)=268.4, p<0.0001, η^2^*p* =0.86; ResNet50: F(7, 294)=290.2, p<0.0001, η^2^*p* =0.87; ResNet101: F(7, 294)=345.8, p<0.0001, η^2^*p* =0.89; AlexNet: F(8, 336)=195.0, p<0.0001, η^2^*p* =0.82; ResNet50-CLIP: F(6, 240)=181.17, p<0.0001, η^2^*p*=0.82, ViT-B_32-CLIP: F(9, 360)=110.53, p<0.0001, η^2^*p*=0.73, ResNet50-SWSL: F(7, 280)=164.7, p<0.0001, η^2^*p*=0.80, ResNeXt101-32×3d-SWSL: F(7, 280)= 225.93, p<0.0001, η^2^*p*=0.85, ViT_large_patch-16_224: F(8, 320)=158.94, p<0.0001, η^2^*p*=0.80).

From visual inspection, many of the metamers from late stages resemble noise rather than natural images (Figure 2e; see Supplementary Figure 1a for examples of metamers generated from different white noise initializations). Moreover, analysis of confusion matrices revealed that for the late model stages there was no detectably reliable structure in participant responses (Supplementary Figure 2). Although the specific optimization strategies we used had some effect on the subjective appearance of the model metamers, human recognition of the generated stimuli remained poor regardless of the optimization procedure (Supplementary Figure 3). The poor recognizability of late-stage metamers was also not explained by optimization difficulties. The activation matches achieved by the optimization were generally good (e.g. with correlations close to 1), and what variation we did observe was not predictive of metamer recognizability (Supplementary Figure 4).

### Metamers for standard auditory deep neural networks are unrecognizable to humans

To investigate whether this phenomenon generalized across modalities, we performed an analogous experiment with two auditory neural networks trained to recognize speech (the word recognition task in the Word-Speaker-Noise dataset (36)). Each model consisted of a biologically-inspired “cochleagram” representation with parameters matched to estimates of the frequency tuning of the human ear (46, 47) followed by a convolutional neural network whose parameters were optimized during training. We tested two model architectures: a ResNet50 architecture (referred to here as CochResNet50) and a model with nine stages (including five convolutional stages) similar to that used in a previously published auditory neural network model (5), referred to here as CochCNN9. Model metamers were generated for clean speech examples from the validation set (Figure 1a). Humans performed a 793-way classification task (5) to identify the word in the middle of either a natural speech example or a model metamer of a speech example generated from a particular model stage (Figure 2e; humans typed responses but were only allowed to enter one of the 793 words from the models’ word-recognition task).

As with the image recognition models, human recognition of the auditory model metamers decreased markedly at late model stages for both architectures (Figure 2f). As with the vision models, there was a significant main effect of human vs. model observer (CochCNN9: F(1, 38)=551.9, p<0.0001, η^2^*p* =0.94; CochResNet50: F(1, 38)=467.8, p<0.0001, η^2^*p* =0.92) and a significant interaction between effect of metamer generation stage and the observer (CochCNN9: F(9, 342)=189.2, p<0.0001, η^2^*p* =0.83; CochResNet50: F(8, 304)=227.4, p<0.0001, η^2^*p* =0.86). Subjectively, the model metamers from deeper stages sound like noise (and appear noise-like when visualized as cochleagrams; Figure 2g). This result suggests that many of the invariances present in these models are not invariances for the human auditory system.

Overall, these results demonstrate that the invariances of many common visual and auditory neural networks are substantially misaligned with those of human perception, even though these models are currently the best predictors of brain responses in each modality.

### Metamers of unsupervised deep network models are also unrecognizable to humans

It is widely appreciated that the learning procedures used to train most contemporary deep neural networks deviate markedly from the learning that occurs in biological systems (48). Perhaps the most fundamental difference lies in supervised learning, as biological systems are not provided with explicit category labels for millions of stimuli during development. Do the divergent invariances evident in neural network models result in some way from supervised training? Metamers are well-suited to address this question given that their generation is not dependent on a classifier, and thus can be generated for any sensory model.

Recent advances in certain types of unsupervised learning have produced neural networks whose representations support classification tasks without requiring millions of labeled examples during training (49). The leading such models are self-supervised, being trained with a loss function favoring representations in which variants of a single training example (different crops of an image, for instance) are similar while those from different training examples are not (Figure 3a). To assess whether this unsupervised training produces more human-like invariances compared to traditional supervised training, we generated model metamers for three such models with a ResNet50 architecture: SimCLR (49), MoCo_V2 (50) and BYOL (51) (examples shown in Figure 3b), and for one model with an AlexNet architecture with group-normalization: IPCL (52). After training, we verified that the learned representations were sufficient to support object classification performance (on ImageNet1K) by training a linear classifier on the final average pooling stage (without changing any of the model weights; Figure 3c&d show classifier performance for each stage of each model). We measured human recognition of metamers from these models as well as those of the same ResNet50 or AlexNet architecture fully trained with supervision (on the ImageNet1K task).

**Figure 3.**
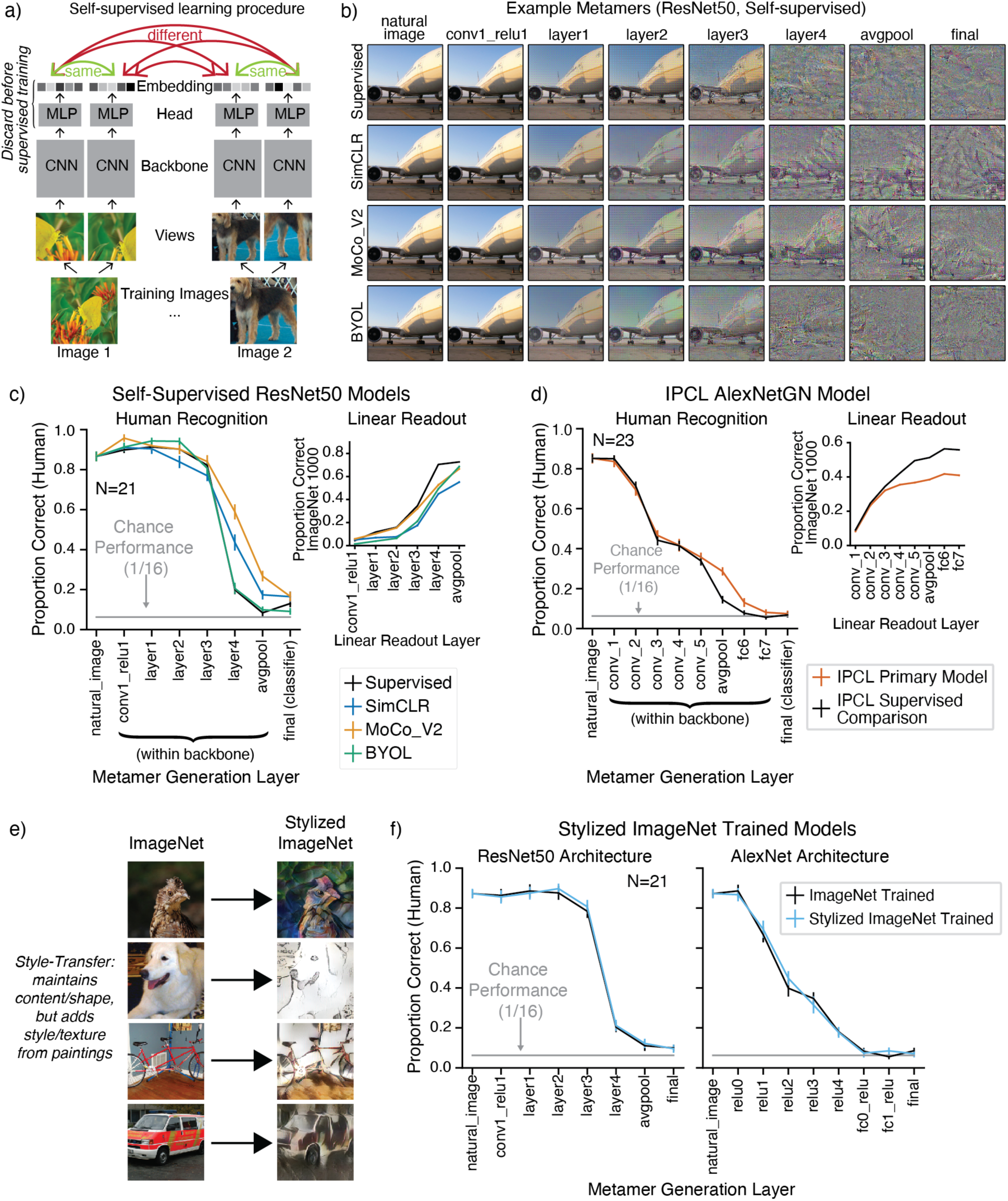
Model metamers are unrecognizable to humans even with alternative training procedures. a) Overview of self-supervised learning, adapted from (49). Models were trained to map multiple views of the same image to nearby points in the embedding space. The SimCLR,MoCo_V2, and IPCL models also had an additional training objective that explicitly pushed apart embeddings from different images. b) Example metamers from select stages of ResNet50 supervised and self-supervised models. In all models, late-stage metamers are mostly unrecognizable. c) Human recognition of metamers from supervised and self-supervised models (left, N=21) along with linear evaluation on the ImageNet1K task at each stage of the models (right). Self-supervised models (SimCLR, MoCo_V2, and BYOL) used a ResNet50 backbone; the comparison supervised model was also a ResNet50. For self-supervised models, model metamers from the “final” stage were generated from a linear classifier trained on the ImageNet1K task on the avgpool representation (intended to validate the representations learned via self-supervision, but included here for completeness). Model recognition curves of model metamers were close to ceiling as in Figure 2, and are omitted here and in later figures for brevity. Here and in d, error bars plot SEM across participants (left) or across three random seeds of model evaluations (right). d) Same as c, but for IPCL self-supervised model and supervised comparison with the same dataset augmentations (N=23). e) Examples of natural and stylized images using the Stylized ImageNet augmentation. Training models on Stylized ImageNet was previously shown to reduce a model’s dependence on texture cues for classification. f) Human recognition of model metamers for ResNet50 and AlexNet architectures trained with Stylized ImageNet (N=21). Removing the texture-bias of models by training on Stylized ImageNet does not result in more recognizable model metamers compared to the standard model.

As shown in Figure 3b-d, the unsupervised models (SimCLR, MoCo_V2, BYOL, and IPCL) produced similar results to those for supervised models: human recognition of model metamers declined at deeper model stages, approaching chance levels for the final stages. Two of the self-supervised ResNet50 models had somewhat more recognizable metamers at intermediate stages, with MoCo_V2 producing the largest boost (significant interaction between model type and model stage F(21, 420)=16.0, p<0.0001, η^2^*p* =0.44). The IPCL model also showed a small but significant improvement in metamer recognizability at intermediate stages (F(9, 198)=3.13, p=0.0018, η^2^*p* =0.12). However, for both architectures, recognition was low in absolute terms, with the metamers bearing little resemblance to the original image they were matched to. Overall, the results suggest that the failure of standard neural network models to pass our metamer test is not specific to the supervised training procedure. This result also demonstrates the generality of the metamers method, as it can be applied to models that do not have a behavioral read-out.

#### Making model recognition more shape-dependent does not resolve discrepant metamers

Other observations of recognition discrepancies between deep neural networks and humans have led to attempts to make model recognition more human-like via changes to the training data. One commonly highlighted discrepancy is the tendency for models to base their judgments on texture rather than shape (53–55). Training datasets of “stylized” images have been designed to reduce this “texture bias” (Figure 3e), and have the effect of increasing a model’s reliance on shape cues, making them more human-like in this respect (53). To assess whether these changes also serve to make model metamers less discrepant, we generated metamers from two models trained on Stylized ImageNet. As shown in Fig. 3f, these models had metamers that were just as unrecognizable to humans as models trained on the standard ImageNet1K training set (no interaction between model type and model stage, ResNet50: F(7, 140)= 0.225, p= 0.979, η^2^*p*= 0.011, AlexNet: F(8, 160)=0.949, p=0.487, η^2^*p*=0.045). This result suggests that metamer discrepancies are not simply due to texture bias in the models.

#### Model metamers reveal invariances of classical hierarchical models of sensory systems

As a further demonstration of the generality of the model metamer method, we generated metamers for classical visual and auditory models that were designed by hand based on neuroscience and engineering principles. Although these models do not perform classification tasks as well as contemporary neural network models, their comparative simplicity might be hypothesized to yield more human-like invariances.

The HMAX vision model is a biologically-motivated architecture with cascaded filtering and pooling operations inspired by simple and complex cells in the primate visual system and was intended to capture aspects of biological object recognition (4, 16). We generated model metamers by matching all units at the S1, C1, S2, or C2 stage of the model (Supplementary Figure 5a). Although HMAX is significantly shallower than the neural network models investigated in previous sections, it is evident that by the C2 model stage its model metamers are comparably unrecognizable to humans (Supplementary Figure 5b&c; significant main effect of model stage, F(4, 76)=351.9, p<0.0001, η^2^*p* =0.95). This classical model thus also has invariances that differ from those of the human object recognition system.

We performed an analogous experiment on a classical model of auditory cortex, consisting of a set of spectrotemporal filters applied to a cochleagram representation (Spectemp model, (56). We used a version of the model in which the convolutional responses are summarized with the mean power in each filter (5, 57, 58) (Supplementary Figure 5d). Metamers from the first two stages were fully recognizable, and very similar to the original audio, indicating that these stages instantiate few invariances, as expected for overcomplete filter bank decompositions (Supplementary Figure 5e&f). By contrast, metamers from the mean power representation were unrecognizable (significant main effect of model stage F(4, 76)=515.3, p<0.0001, η^2^*p* =0.96), indicating that this model stage produces invariances that humans do not share (plausibly because the human speech recognition system retains information that is lost in the averaging stage).

Overall, these results show how metamers can reveal the invariances present in classical models as well as state-of-the-art deep neural networks, and demonstrate that both types of models fail to fully capture the invariances of biological sensory systems.

### Adversarial training of visual models increases human recognition of model metamers

A known peculiarity of contemporary artificial neural networks is their vulnerability to small adversarial perturbations designed to change the class label assigned by a model (23–26, 59). Such perturbations are typically imperceptible to humans due to their small magnitude, but can drastically alter model decisions, and have been the subject of intense interest in part due to the security risk they pose for machine systems. One way to reduce this vulnerability is to generate perturbation-based adversarial examples during training and add them to the training set, forcing the model to learn to recognize the perturbed images as the “correct” human-interpretable class (Figure 4a) (60). This procedure has been found to yield models that are less susceptible to adversarial examples for reasons that remain debated (61).

**Figure 4.**
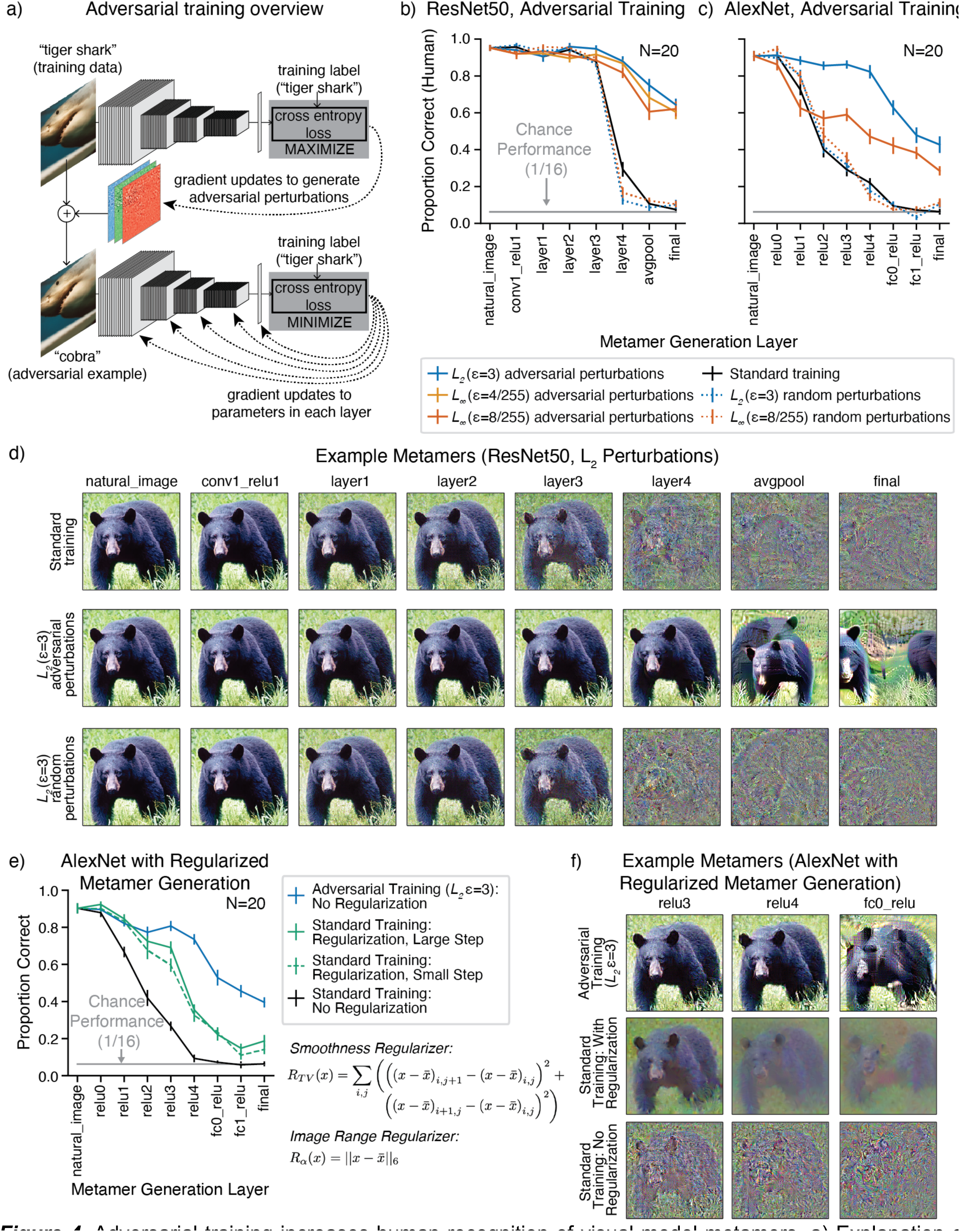
Adversarial training increases human recognition of visual model metamers. a) Explanation of adversarial training. Adversarial examples are derived at each training step by finding an additive perturbation to the input that moves the classification label away from the training label class (top). These derived adversarial examples are then provided to the model as training examples and used to update the model parameters (bottom). The resulting model learns to classify the adversarial examples as the training data label, and is subsequently more robust to adversarial perturbations than if standard training were used. As a control experiment, we also trained models with random perturbations to the input rather than adversarial perturbations. b) Human recognition of visual model metamers on a 16-way classification task with metamers generated from ResNet-50 models (N=20) trained with and without adversarial or random perturbations. Here and elsewhere, error bars plot SEM across participants. c) Same as b, but for AlexNet models (N=20). In both ResNet50 and AlexNet models, adversarial training leads to more recognizable metamers at the deep stages of the models, though in both cases the metamers remain less than fully recognizable. d) Example visual model metamers for ResNet50 models trained with and without adversarial or random *L_2_* perturbations. e) Recognizability of model metamers from standard trained models with and without regularization, compared to that for an adversarially trained model (N=20). Two regularization terms were included in the optimization: a total variation regularizer to promote smoothness, and a constraint to stay within the image range. Two different optimization step sizes were evaluated. f) Example metamers for an adversarially trained AlexNet (top) and a standard AlexNet with (middle) and without (bottom) regularization.

We asked whether reducing adversarial vulnerability in this way would improve human recognition of model metamers. A priori it was not clear what to expect. Perturbation-based adversarial examples can be viewed as the converse of a model metamer, in that they are generated by perturbations to which humans are invariant but which cause changes in model decisions. Making models robust to adversarial perturbations causes them to exhibit more of the invariances of humans (the shaded orange covers more of the blue outline in Figure 1b), but it is not obvious that this will reduce the model invariances that are not shared by humans (i.e., to decrease the orange outlined regions that don’t overlap with blue shaded region in Figure 1b). Previous work visualizing latent representations of visual neural networks suggested that robust training might make model representations more human-like (62), but human recognition of model metamers had not been behaviorally evaluated.

We first generated model metamers for vision models trained to be adversarially robust (ImageNet1K-trained ResNet50 and AlexNet architectures) (62). As a control, we also trained models with equal magnitude perturbations in random, rather than adversarial, directions. Such random perturbations are typically ineffective at preventing adversarial attacks (59). As intended, models with adversarial perturbations during training were more robust to adversarial examples than the standard model or models with random perturbations (Supplementary Figure 6a&b).

For both architectures, metamers for the robust models were significantly more recognizable than those from the standard model (Figure 4b-d; Supplementary Figure 1), evident as a significant main effect of training type (repeated measures ANOVAs comparing human recognition of standard and each adversarial model, significant main effect in each of the five models, F(1, 19)>104.61, p<0.0001, η^2^*p*>0.85). Training with random rather than adversarial perturbations did not yield the same benefit (significant main effect of random vs. adversarial for each perturbation of the same ε type and size, F(1, 19)>121.38, p<0.0001, η^2^*p*>0.86). Model metamers were more recognizable to humans for some adversarial training variants than others, but all variants that we tried produced a human recognition benefit. It was nonetheless the case that metamers from all variants remained less than fully recognizable to humans when generated from the deep stages. We note that performance is inflated by the use of a 16-way alternative-force-choice (AFC) task, for which above-chance performance is possible even with severely distorted images. See Supplementary Figure 7 for an analysis of the consistency of metamer recognition across human observer, and examples of metamers that are most and least recognizable.

Given that metamers from adversarially trained models look less noise-like than those from standard models, and that standard models may over-represent high spatial frequencies compared to adversarially robust models (63), we wondered whether their recognition advantage could be replicated in a standard-trained model by including a smoothness regularizer in the metamer optimization. Such regularizers are common in neural network visualizations, and although we view them as side-stepping the goal of human-model comparison, it was nonetheless of interest to assess their effect. We implemented the regularizer used in a well-known visualization paper (64) which biases the solution to the metamer optimization to have low total pixel variation (encouraging smoothness). As shown in Figure 4e, adding smoothness regularization to the metamer generation procedure for the standard-trained AlexNet model improved the recognizability of its metamers, but not as much as did adversarial training (and did not come close to generating metamers as recognizable as natural images; see Supplementary Figure 8 for examples generated with different regularization coefficients). This result suggests that a) the benefit of adversarial training is not simply replicated by imposing smoothness constraints, and b) discrepant metamers more generally cannot be resolved simply with the addition of a smoothness prior.

#### Adversarial training of auditory models increases human recognition of model metamers

To investigate whether adversarial training also leads to more human-like auditory representations, we trained CochResNet50 and CochCNN9 architectures on the word recognition task described above, using both standard and adversarial training. Because the auditory models contain a fixed cochlear stage at their front end, there are two natural places to generate adversarial examples (they can be added to the waveform or the cochleagram), and we explored both for completeness.

We first investigated the effects of adversarial perturbations to the waveform (Figure 5a, Supplementary Figure 6c&d). As a control experiment, we again trained models with random, rather than adversarial, perturbations. As with the visual models, human recognition was generally better for metamers from adversarially trained models (Figure 5b&c; ANOVAs comparing standard and each adversarial model, significant main effect in 4/5 cases: F(1, 19)>9.26, p<0.0075, η^2^*p*>0.33; no significant main effect for CochResNet50 with *L2* (ε=1) perturbations: F(1, 19)=0.29, p=0.59, η^2^*p*=0.015). This benefit again did not occur for models trained with random perturbations (ANOVAs comparing each random and adversarial perturbation model with the same ε type and size; significant main effect in each case: F(1, 19)>4.76, p<0.0444, η^2^*p*>0.20). The model metamers from the robust models are visibly less noise-like when viewed in the cochleagram representation (Figure 5d).

**Figure 5.**
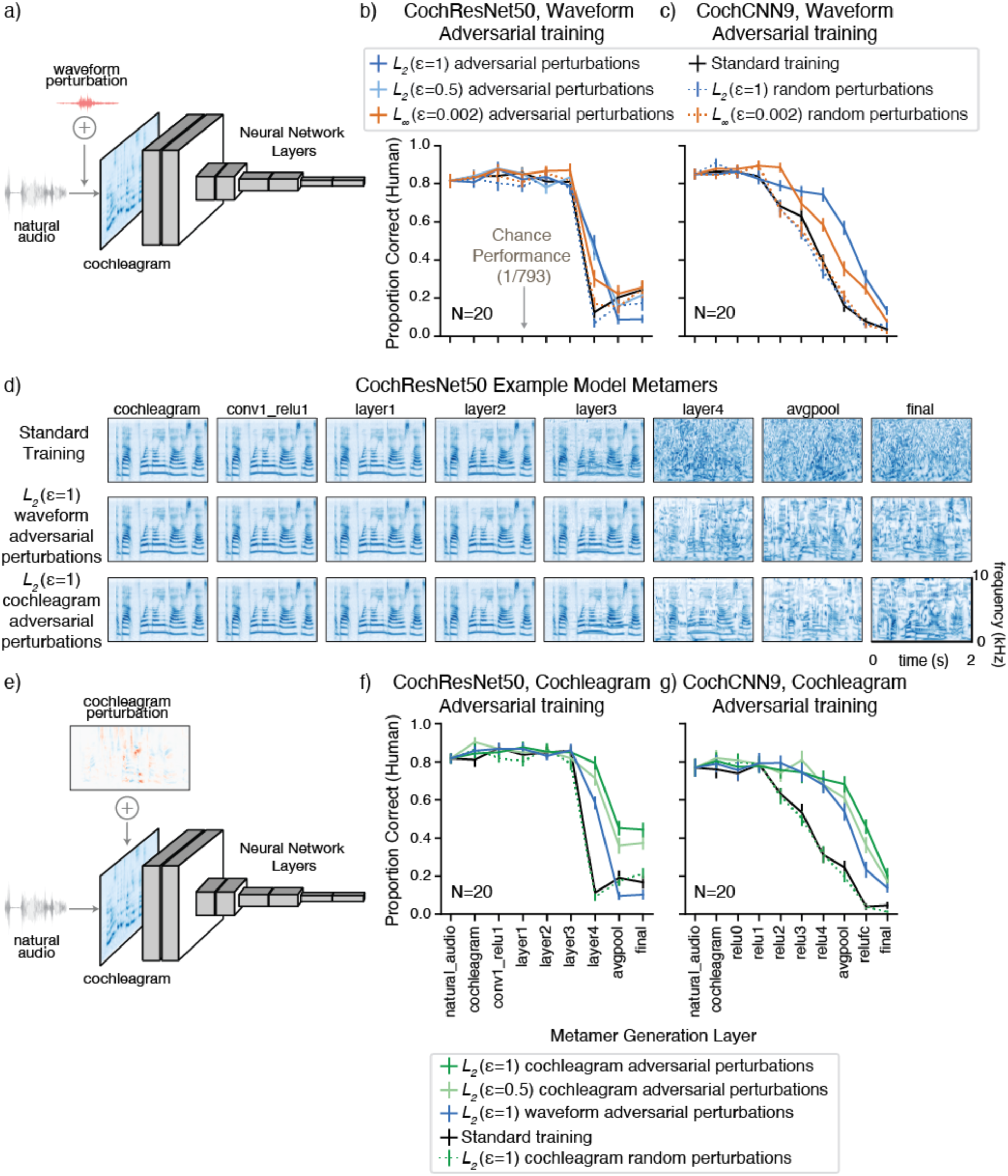
*(next page)*. Adversarial training increases human recognition of auditory model metamers. a) Schematic of auditory convolutional neural networks with adversarial perturbations applied to the waveform input. b) Human recognition of auditory model metamers from CochResNet50 (N=20) and c) CochCNN9 (N=20) models with adversarial perturbations generated in the waveform space (models trained with random perturbations are also included for comparison). When plotted here and in f&g, chance performance (1/793) is indistinguishable from the x-axis and error bars plot SEM across participants. *L2 (*ε=0.5) waveform adversaries were only included in the CochResNet50 experiment. d) Cochleagrams of example model metamers from CochResNet50 models trained with waveform and cochleagram adversarial perturbations. e) Schematic of auditory convolutional neural networks with adversarial perturbations applied to the cochleagram stage. f) Human recognition of auditory model metamers from models trained with cochleagram adversarial perturbations are more recognizable for CochResNet50 and g) CochCNN9 models compared to those from models trained with waveform perturbations.

We also trained models with adversarial perturbations to the cochleagram representation, whose fixed components enabled norm-based constraints on the perturbation size analogous to those used for input-based adversarial examples (Figure 5e). Models trained on cochleagram adversarial examples had significantly more recognizable metamers than both the standard models and the models adversarially trained on waveform perturbations (Figure 5f&g; ANOVAs comparing each model trained with cochleagram perturbations vs. the same architecture trained with waveform perturbations; significant main effect in each case: F(1, 19)>4.6, p<0.04, η^2^*p*>0.19; ANOVAs comparing each model trained with cochleagram perturbations to the standard model; significant main effect in each case: F(1, 19)>102.25, p<0.0001, η^2^*p*>0.84). The effect on metamer recognition was again specific to adversarial perturbations (ANOVAs comparing effect of training with adversarial vs. random perturbations with the same ε type and size: F(1, 19)>145.07, p<0.0001, η^2^*p*>0.88).

Although the perturbation sizes in the two model stages are not directly comparable, in each case we chose the size to be large enough that the model showed robustness to adversarial perturbations while not being so large that the model could not perform the task (as is standard for adversarial training). Further, training models with perturbations generated at the cochleagram stage resulted in substantial robustness to adversarial examples generated at the waveform (Supplementary Figure 6e&f). These results suggest that the improvements from intermediate-stage perturbations may in some cases be more substantial than those from perturbations to the input representation. They also highlight the utility of model metamers for evaluating model modifications, in this case adversarial training with perturbations to intermediate model representations.

Overall, these results suggest that training alterations intended to mitigate undesirable characteristics of artificial neural networks (adversarial vulnerability) can cause their invariances to become more like those of humans in both the visual and auditory domain. However, substantial discrepancies remain even with these alterations. In both modalities, many model metamers from deep model stages remain unrecognizable even after adversarial training.

### Human recognition of metamers dissociates from adversarial vulnerability

Although we found that adversarial training increased human recognizability of model metamers, the degree of robustness that resulted from the training was not itself predictive of metamer recognizability. We first examined all the visual models from Figures 2-5, comparing their adversarial robustness with the recognizability of their metamers from the final model stage (this stage was chosen as it exhibited considerable variation in recognizability across models). As shown in Figure 6a, there is a correlation between robustness and metamer recognizability for visual models (ρ=0.73, p<0.001), but it is mostly driven by the overall difference between two groups of models: those that were adversarially trained and those that were not.

**Figure 6.**
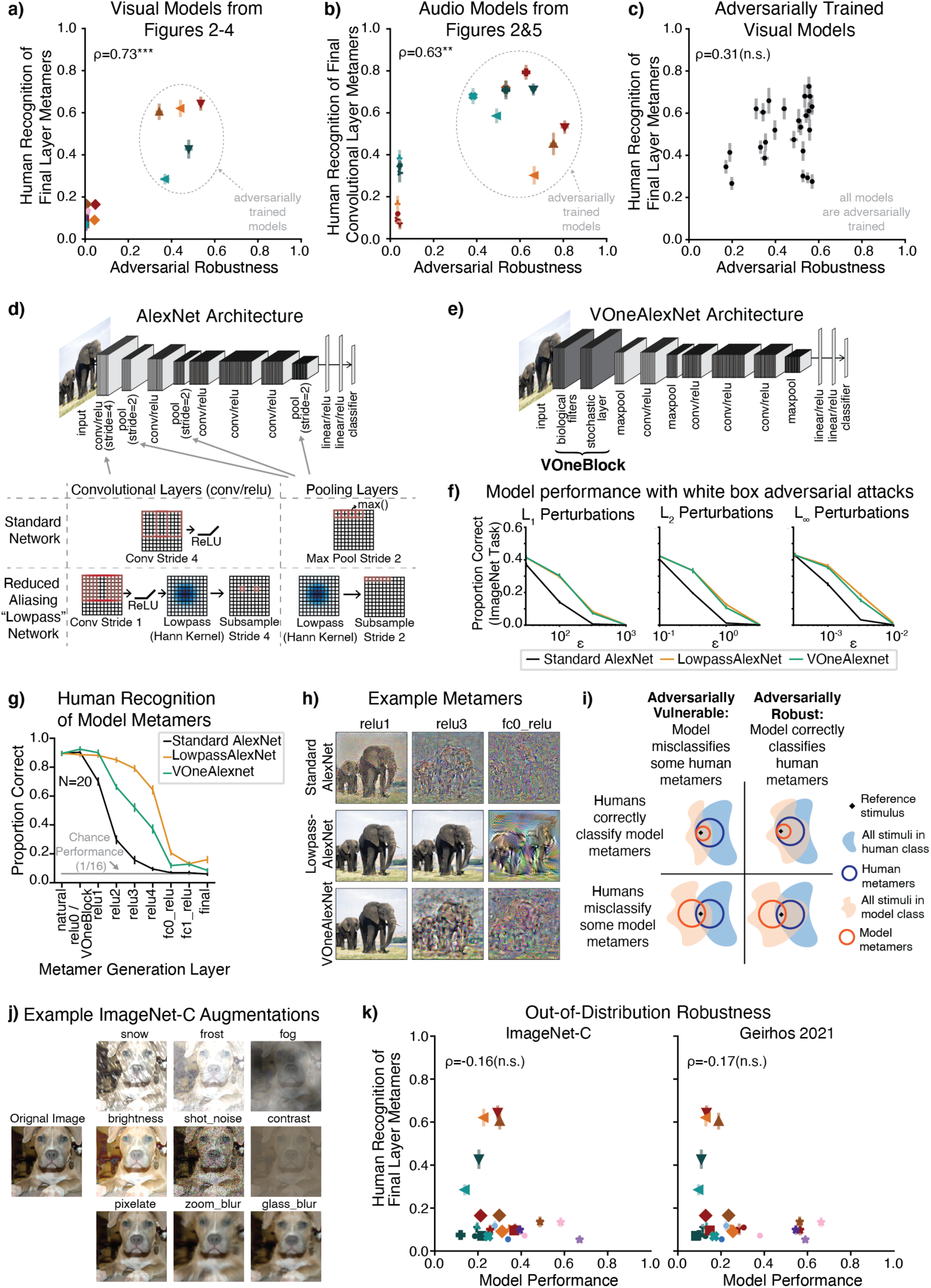
(previous page). Human recognition of model metamers dissociates from adversarial vulnerability. a) Scatter plot of adversarial robustness of visual models vs. human recognizability of final model stage model metamers (N=26 models). Adversarial robustness is quantified as the average robustness to L2 (ε=3) and L∞ (ε=4/255) adversarial examples, normalized by the performance on natural images. Symbols follow Figure 7a. b) Same as a, but for auditory models (N=17 models). Adversarial robustness is quantified as the average robustness to L2 (ε=10^-0.5^) and L∞ (ε=10^-2.5^) adversarial perturbations of the waveform, normalized by the performance on natural audio. Symbols follow Figure 7c. c) Scatter plot of adversarial robustness of a set of visual adversarially trained models vs. human recognizability of the final model stage model metamers (N=25 models). d) Operations included in AlexNet architecture to reduce aliasing. Strided convolutions were replaced with a sequence of four operations: convolution with a learnable kernel using a stride of 1, ReLU, convolution with a fixed lowpass filter, and subsampling with the original convolution stride. Max Pooling was replaced with a sequence of two operations: convolution with a fixed lowpass filter, and subsampling with the original pooling stride. The model with these modified operations more closely obeys the sampling theorem and is referred to as “LowpassAlexNet”. e) Schematic of VOneBlock input to AlexNet backbone architecture. The VOneBlock includes a stage of fixed Gabor filters, and a stochastic stage with Gaussian noise. The stochastic stage was turned on for training but turned off for evaluations of adversarial robustness and for metamer generation. f) Adversarial vulnerability, assessed via recognition accuracy on 1000-way ImageNet classification task with adversarial perturbations of different sizes added to the images. LowpassAlexNet and VOneAlexNet both produce a modest increase in adversarial robustness (greater accuracy for large adversarial perturbations) compared to the standard-trained AlexNet model. Error bars plot SEM across five subsets of training images. g) Human recognition of model metamers generated from LowpassAlexNet, VoneAlexNet and standard AlexNet models on the 16-way classification task (N=20). LowpassAlexNet has substantially more recognizable model metamers than the VOneNet, demonstrating a dissociation between human recognition of model metamers and a model’s susceptibility to adversarial examples. Error bars plot SEM across participants. h) Example model metamers from experiment in (d). i) Human-recognizability of model metamer is dissociable from adversarial robustness. Adversarial examples are cases where a model misclassifies stimuli that are metameric to humans (left row). This schematic depicts a reference stimulus for which the model gives the correct class label, such that the adversarial perturbation induces a classification error. Model metamers may be unrecognizable to humans even if the model is robust to adversarial examples (bottom right). Tests involving model metamers are thus complementary to those involving adversarial examples. j) Example augmentations applied to images in tests of out-of-distribution robustness. k) Scatter plot of out-of-distribution robustness vs. human recognizability of final stage model metamers (N=26 models). Large-scale training often produces improved robustness (models denoted with ★ symbols), but not an improvement in metamer recognizability. Symbols follow Figure 7a.

The auditory models showed a similar relationship as the visual models (Figure 6b). When both standard and adversarially trained models were analyzed together, there was a significant correlation between metamer recognizability and robustness (ρ=0.63, p=0.004) that was driven by the overall difference between the two groups of models. This analysis was performed using intermediate stage metamers (layer4 for CochResNet50, and ReLU4 for CochCNN9) because the final stage metamer recognizability was sufficiently low overall (because the auditory recognition task was much harder than the visual task: 793 vs. 16 possible classes) that a floor effect plausibly explained the absence of a significant correlation for the final stage metamers (ρ=0.21, p=0.215). As with the visual models, there is no obvious relationship when considering just the adversarially trained models.

To better assess whether variations in robustness produce variation in metamer recognizability, we compared the robustness of a large set of adversarially trained models (taken from a well-known robustness evaluation (65)) to the recognizability of their metamers from the final model stage. As shown in Figure 6c, despite the considerable variation in both robustness and metamer recognizability, the correlation between these two measures was low and non-significant (ρ=0.31, p=0.099). Overall, it thus seems that something about the adversarial training procedure leads to more recognizable metamers, but that robustness per se does not drive the effect.

Adversarial training is not the only means of making models adversarially robust. But when examining other sources of adversarial robustness, we again found examples where a model’s susceptibility to adversarial examples was not predictive of the recognizability of its metamers. Here we present results for two models that had similar adversarial robustness, one of which had much more recognizable metamers than the other.

The first model was a CNN that was modified to reduce aliasing (LowpassAlexNet). Because many traditional neural networks contain downsampling operations (e.g. pooling) without a preceding lowpass filter, they violate the sampling theorem (36, 66) (Figure 6d). It is nonetheless possible to modify the architecture to reduce aliasing, and such modifications have been suggested to improve model robustness to small image translations (21, 22). The second model was a CNN that contained an initial processing block inspired by primary visual cortex in primates (67, 68) featuring hard-coded Gabor filters and a stochastic response component present during training (VOneAlexNet, Figure 6e). The inclusion of this block has been previously demonstrated to increase adversarial robustness. It was a priori unclear whether either model modification would improve human recognizability of the model metamers.

Both architectures were comparably robust to standard (*Lp*-bounded) adversarial perturbations (Figure 6f; no significant main effect of architecture for perturbations, F(1, 8)<4.5, p>0.10, η^2^*p*<0.36 for all perturbation types), and both were more robust than the standard AlexNet (main effect of architecture for perturbations F(1, 8)>137.4, p<0.031, η^2^*p*>0.94, for all adversarial perturbation types for both VOneAlexNet and LowPassAlexNet). Both were also more robust than the standard model to “fooling images” (23) and “feature adversaries” (69), and LowpassAlexNet was not more robust than VOneAlexNet to these types of adversaries (Supplementary Figure 9). However, metamers generated from LowpassAlexNet were substantially more recognizable than metamers generated from VOneAlexNet (Figure 6g,h; main effect of architecture: F(1, 19)=71.7, p<0.0001, η^2^*p*>0.79; interaction of architecture and model stage: F(8, 152)=21.8, p<0.0001, η^2^*p*>0.53). This result further demonstrates that human recognizability of model metamers and susceptibility to adversarial examples are dissociable. The model metamer test thus provides a complementary comparison of model invariances that can differentiate models even when adversarial robustness does not.

These adversarial robustness-related results may be understood in terms of the four types of stimulus sets originally shown in Figure 1b, four configurations of which are depicted in Figure 6i Adversarial examples are stimuli that are metameric to a reference stimulus for humans but that are classified differently from the reference stimulus by a model. Adversarial robustness thus corresponds to a situation where the human metamers for a reference stimulus fall completely within the set of stimuli that are given the reference class label by a model (blue outline contained within orange shaded region in 6i, right column). This condition does not imply that all model metamers will be recognizable to humans (orange outline contained within the blue shaded region, top row). These theoretical observations motivate the use of model metamers as a complementary model test, and are confirmed by the empirical observations of this section. The effect of adversarial training on model metamers (Figures 4&5) thus appears to be somewhat specific to the training method – a model’s adversarial robustness is not always predictive of the human recognizability of its metamers.

### Human recognition of metamers also dissociates from other forms of robustness

Neural network models have also been found to be less robust than humans to images that fall outside their training distribution (e.g. line drawings, silhouettes, high-pass filtered images that qualitatively differ from the photos that form the basis of the common ImageNet1K training set, Figure 6j) (18, 70, 71). This type of robustness has been found to be improved by training models on substantially larger data sets (72). We compared model robustness to such “out-of-distribution” images to the recognizability of their metamers from the final model stage (the model set included several models trained on large-scale data sets, taken from Figure 2d, along with all the other models from Figures 2c, 3 & 4). As shown in Figure 6k, this type of robustness (measured by two common benchmarks) was again not correlated with metamer recognizability (ImageNet-C: ρ=-0.16, p=0.227, Geirhos 2021: ρ=-0.17, p=0.215).

#### Current model-brain comparisons do not capture metamer differences between models

Are the differences between models shown by the metamer test similarly evident when using standard brain comparison benchmarks? To address this question, we used such benchmarks to evaluate the visual and auditory models described above in Figures 2-5. For the visual models, we used the Brain-Score platform to measure the similarity of model representations to neural benchmarks for visual areas V1, V2, V4 and IT (38, 42). The platform’s similarity measure combines a set of model-brain similarity metrics, primarily measures of variance explained by regression-derived predictions. For each model, the score was computed for each visual area using the model stage that gave highest similarity in held-out data for that visual area. We then compared this neural benchmark score to the recognizability of the model’s metamers from the same stage used to generate the neural predictions. This analysis showed modest correlations between the two measures for V4 and IT (Figure 7b; V1: ρ=0.10, p=0.62; V2: ρ=0.13, p=0.51; V4: ρ=0.40, p=0.04; IT: ρ=0.45, p=0.02). We note that none of these would survive Bonferroni correction, and that these correlations were well below the presumptive noise ceiling (given the split-half reliability of both the metamer recognizability scores and the model-brain similarity scores (73), the noise ceiling of the correlation was ρ=0.92 for IT). Thus, most of the variation in metamer recognizability was not captured by standard model-brain comparison benchmarks.

**Figure 7.**
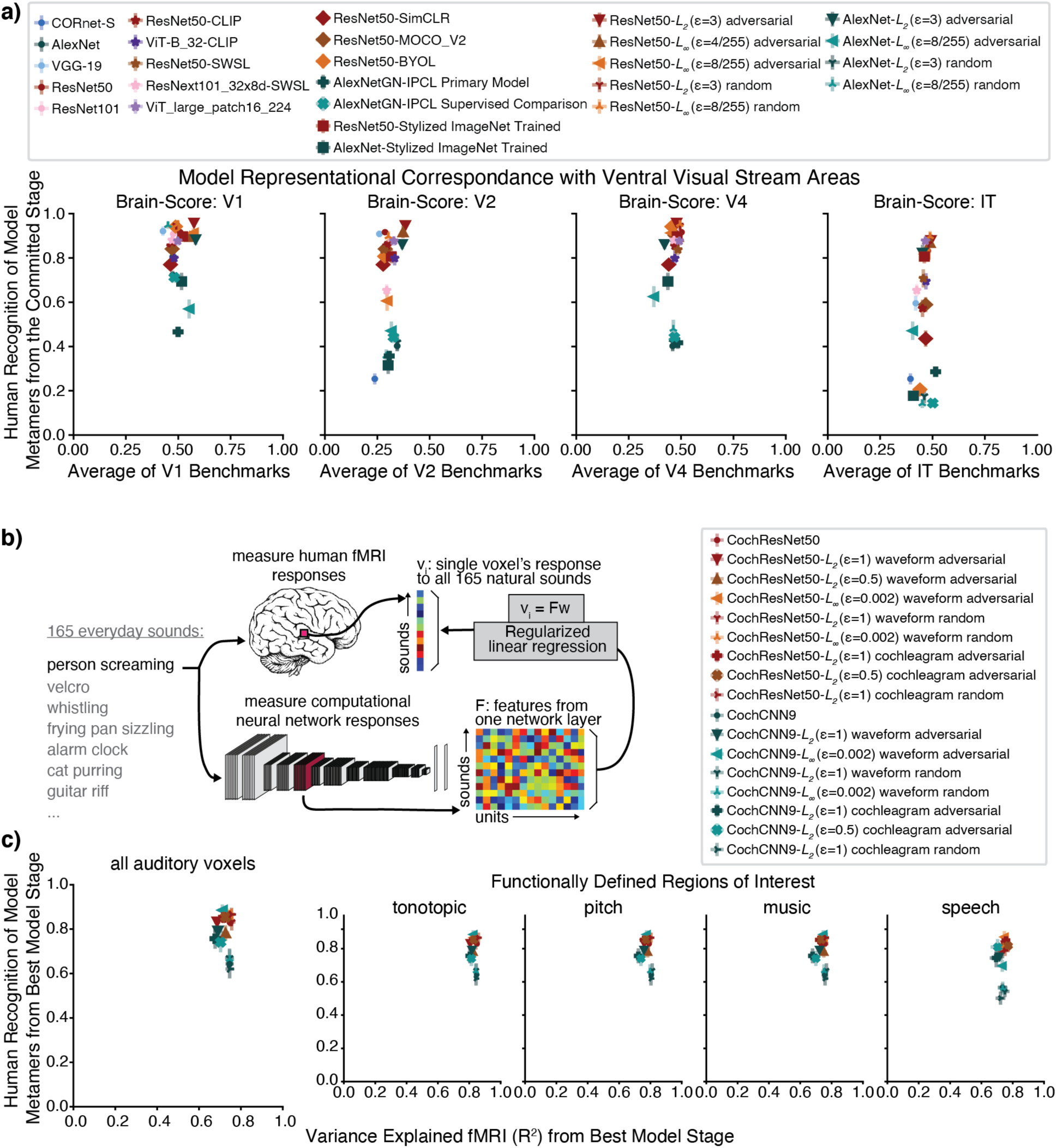
Human recognition of model metamers dissociates from model predictions of brain responses. a) Scatter plots of the human recognizability of a model’s metamers vs. the model-brain similarity for four areas of the ventral stream as assessed by a commonly used set of benchmarks (38, 42). The benchmarks mostly consisted of the neurophysiological variance explained by model features via regression. A single model stage was chosen for each model and brain region that produced highest similarity in a test data set. Graph plots metamer recognizability and model-brain similarity for this stage. b) To obtain auditory brain predictions, 165 natural sounds presented to humans in an fMRI experiment (74) were also presented to neural network models. The model’s time-averaged unit responses in each stage were used to predict each auditory cortical voxel’s response by a linear mapping fit to the responses to a subset of the sounds with ridge regression. For each neural network model, the best-predicting stage was selected for each participant using independent data. Model predictions were evaluated on a set of held-out sounds. Schematic of procedure is reproduced from (5). c) Average voxel response variance explained by the best-predicting stage of each auditory model from Figures 2 and 5, plotted against the metamer recognizability for that model stage. We performed this analysis across all voxels in auditory cortex (left) as well as within a selection of auditory functional ROIs (right). The variance explained (R^2^) was measured for the best-predicting stage of the neural network models, chosen individually for each participant and each voxel subset. For each participant, the other participants’ data was used to choose the stage that yielded the best predictions, and then the R^2^ from this stage for the held-out participant was included in the average. Error bars on each data point plot SEM across participants.

We performed an analogous analysis for the auditory models, using a large data set of human auditory cortical responses (74) (fMRI responses to a large set of natural sounds) that had previously been used to evaluate neural network models of the auditory system (5, 75). We analyzed voxel responses within four regions of interest in addition to all of auditory cortex, in each case again choosing the best-predicting model stage, measuring the variance it explained in held-out data, and comparing that to the recognizability of the metamers from that stage (Figure 7b). The correlation between metamer recognizability and explained variance in the brain response was not significantly different from 0 when all voxels were considered (ρ=0.06, p=0.81; Figure 7c). We did find a modest correlation within one of the ROIs (tonotopic: ρ=-0.13, p=.63; pitch: ρ=-0.31, p=.22; music: ρ=-0.19, p=.47; speech: ρ=0.58, p=.016), but it was again well below the presumptive noise ceiling (which ranged from ρ=0.78 to ρ=0.87, depending on the ROI), and would not survive Bonferroni correction.

We conducted analogous analyses using representational similarity analysis instead of regression-based explained variance to evaluate model-brain similarity, with similar conclusions (Supplementary Figure 10). Overall, the results indicate that the metamer test is complementary to traditional metrics of model-brain fit.

### Human recognition of a model’s metamers is predicted by their recognizability to other models

Are one model’s metamers recognizable by other models? We addressed this issue by taking all the models we trained for one modality, holding one model out as the “generation” model, and then presenting its metamers to each of the other models (“recognition” models), measuring the accuracy of their class predictions (Figure 8a). We repeated this procedure with each model as the generation model. Accuracy for all combinations of recognition and generation models is shown in Supplementary Figures 11 and 12. As a summary measure for each generation model, we averaged the accuracy across the recognition models (Figure 8a, right). Because we had trained the ResNet50 and CochResNet50 architectures with several variants of self-supervised and adversarial training in addition to standard supervised training, we performed further analysis of these architectures (Figure 8b&c).

**Figure 8.**
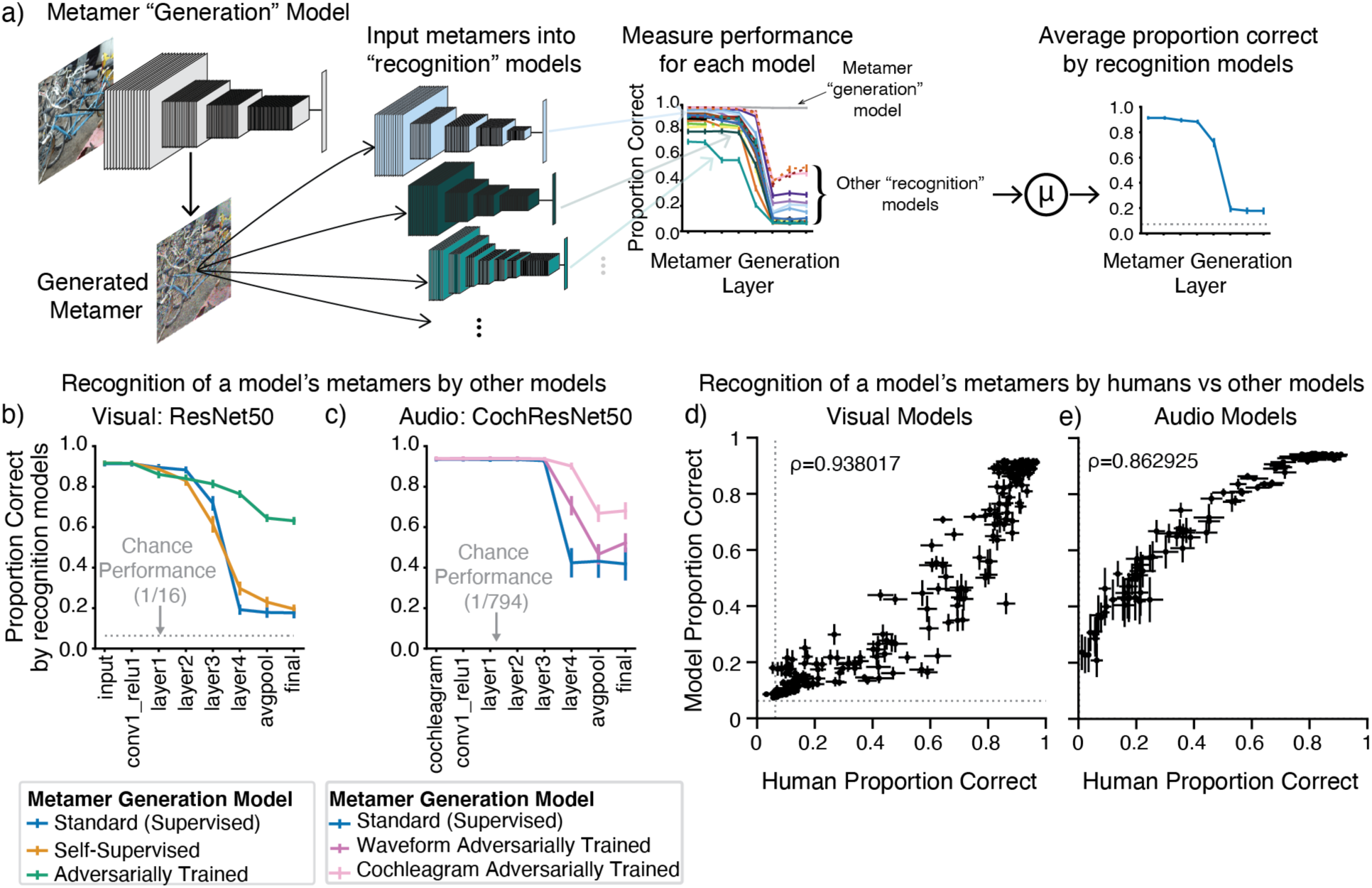
Human recognition of a model’s metamers is correlated with their recognition by other models. a) Model metamers were generated for each stage of a “generation” model (one of the models from Figures 2c-d, 3, 4, and 6g for visual model analysis, and from Figures 2f and 5 for auditory model analysis). The associated metamers are input into “recognition” models (all other models from Figures 2c-d, 3, 4, and 6g for visual model analysis, and from Figures 2f and 5 for auditory model analysis). We measured recognition of the generating model’s metamers by each recognition model, averaging the accuracy over all recognition models (excluding the generation model). This analysis is shown here for a standard-trained ResNet50 image model. Error bars are SEM over recognition models. b) Average model recognition of metamers generated from the standard ResNet50, the three self-supervised ResNet50 models, and the three ResNet50 models trained with adversarial perturbations. In model groups with multiple generating models, we averaged each recognition model’s accuracy curve across all of the generating models, then averaged these curves across the recognition models. Error bars are the SEM over recognition models. Model metamers from deep stages tend to be unrecognizable to other models, but models trained with adversarial perturbations have metamers that are more recognizable by other models. c) Same as b, but for auditory models, with metamers generated from the standard CochResNet50, three CochResNet50 models with waveform adversarial perturbations, and two CochResNet50 models with cochleagram adversarial perturbations. Chance performance is 1/794 for the models because (unlike the human experimental participants) they had a “null” (no speech) class label as a possible prediction in addition to the 793 word labels. d, e) Correlation between human and model recognition of another model’s metamers for visual (d) and auditory (e) models. The abscissa of each data point plots the average human recognition accuracy of metamers generated from one stage of a model. The ordinate of each data point plots the average recognition by other models of those metamers. In both modalities, human recognition of a model’s metamers is highly correlated with the accuracy of other models at recognizing those same model metamers (quantified by Spearman rank-ordered correlation coefficient).

In the vision models, metamers from deep stages of the standard supervised trained ResNet50 were generally not recognized by other models (Figure 8b, blue), suggesting that many of the invariances in this model are not present in the other models. A similar trend held for the models trained with self-supervision (Figure 8b, orange). By contrast, metamers from the adversarially trained models were more recognizable to other models (Figure 8b, green). We saw an analogous metamer transfer boost from the model with reduced aliasing (LowPassAlexNet), for which metamers for intermediate stages were more recognizable to other models (Supplementary Figure 12). Similar results held for auditory models (Figure 8c), though metamers from the standard supervised CochResNet50 transferred better to other models than did those for the supervised vision model, perhaps due to the shared cochlear representation present in all auditory models, which could increase the extent of shared invariances.

These results suggest that models tend to contain idiosyncratic invariances, in that their metamers vary in ways that render them unrecognizable to other models. The results also clarify the effect of adversarial training. Specifically, they suggest that adversarial training is removing some of the idiosyncratic invariances of standard-trained deep neural networks, rather than learning new invariances that are not shared with other models (in which case their metamers would not have been better recognized by other models). The architectural change that reduced aliasing had a similar effect, albeit limited to the intermediate model stages.

The average model recognition of metamers generated from a given stage of another model is strikingly similar to human recognition of the metamers from that stage (compare Figures 8b and 8c to Figures 3, 4, and 5, and Supplementary Figure 13 to Figure 6d). To quantify this similarity, we plotted the average model recognition for metamers from each stage of each generating model against human recognition of the same stimuli, revealing a strong correlation for both visual (Figure 8d) and auditory (8e) models. This result suggests that the human-model discrepancy revealed by model metamers reflects invariances that are often idiosyncratic properties of a specific neural network, leading to impaired recognition by both other models and human observers. One consequence of this result is that a collection of other models may be a reasonable proxy for a human observer -- rather than running a new psychophysical experiment any time there is a new model to evaluate, one could test the model’s metamers on a set of other models, facilitating the automation of searches for better models of perception.

## Discussion

We used model metamers to reveal invariances of deep artificial neural networks, and compared these invariances to those of humans by measuring human recognition of visual and auditory model metamers. We found that metamers of standard deep neural networks are dominated by invariances that are absent from human perceptual systems, in that metamers from deep model stages are typically completely unrecognizable to humans. This was true across modalities (both visual and auditory) and across training methods (supervised vs. self-supervised training). We identified ways to make model metamers more human-recognizable in both the auditory and visual domains, including a new type of adversarial training for auditory models using perturbations at an intermediate model stage. Although there was a substantial effect on metamer recognizability from one common training method to reduce adversarial vulnerability, we found that metamers could reveal model differences that were not evident by measuring adversarial vulnerability alone. Moreover, the model improvements revealed by model metamers were not obvious from standard brain prediction metrics. These results show that metamers provide a model comparison tool that complements the standard benchmarks that are in widespread use.

Even though some models produced significantly more recognizable metamers than others, metamers generated from late model stages remained less recognizable than natural images or sounds in all cases we tested, suggesting that substantial further improvements are needed to align model representations with those of biological sensory systems. The fact that one model generally cannot recognize another model’s metamers indicates that the discrepancy is at least partially caused by invariances that are idiosyncratic to a model (being absent in both humans and other models). Might humans analogously have their own invariances that are specific to an individual? This possibility is difficult to explicitly test given that we cannot currently sample human metamers (model metamer generation relies on having access to the model’s parameters and responses, which are currently beyond reach for biological systems). If idiosyncratic invariances were present in humans as well, the phenomenon we have described here might not represent a human-model discrepancy, and could instead be a general property of recognition systems. The main argument against this interpretation is that several model modifications motivated by engineering considerations – different forms of adversarial training, and architectural modifications to reduce aliasing – substantially reduced the idiosyncratic invariances present in standard deep neural network models. These results suggest that idiosyncratic invariances are not unavoidable in a recognition system. Moreover, the set of modifications explored here was far from exhaustive, and we see no reason why the idiosyncratic model invariances could not be further alleviated with alternative training or architecture changes in the future.

### Relation to previous work

The methods we use to synthesize model metamers are not new. Previous neural network visualizations have also used gradient descent on the input to visualize representations (76), in some cases matching the activations at individual stages as we do here (64). However, the significance of these visualizations for evaluating neural network models of biological sensory systems has received relatively little attention. One contributing factor may be that model visualizations have often been constrained by added natural image priors or other forms of regularization (77) that help make model visualizations look more natural, but mask the extent to which they otherwise diverge from a perceptually meaningful stimulus. For this reason, we intentionally avoided priors or other regularization when generating model metamers, as they defeat the purpose of the metamer test. When we explicitly measured the benefit of regularization, we found that it did boost recognizability somewhat, but that it was not sufficient to render model metamers fully recognizable, or to reproduce the benefits of model modifications that improve metamer recognizability (Figure 4e).

Another reason the discrepancies we report here have not been widely discussed within neuroscience is that most studies of neural network visualizations have relied on small numbers of examples rather than systematically measuring recognizability to human observers (in part because these visualizations are primarily reported within computer science, where such experiments are not the norm). By contrast, our work used human perceptual experiments to quantify the divergence between human and model invariances, revealing a widespread mismatch between humans and current state-of-the-art computational models. We found controlled experiments to be essential. Prior to running full-fledged experiments we always conducted the informal exercise of generating examples and evaluating them subjectively. Although the largest effects were evident informally, the variability of natural images and sounds made it difficult to predict with certainty how an experiment would turn out. It is thus critical to substantiate informal observation with controlled experiments in humans.

Metamers are also related to a type of adversarial example generated by adding small perturbations to an image from one class such that the activations of a classifier match those of a reference image from a different class. These “invariance-based adversarial examples” yield the same model activations at the classification stage (sometimes referred to as “feature adversaries” (69) or “feature collisions” (78)), but are seen as different classes by humans (albeit again tested informally with a small number of examples rather than a systematic experiment) (79, 80). Our method differs in probing the model invariances without any explicit bias to cause the metamers to appear different to humans, but our results may reflect some of the same properties evident in these alternative adversarial stimuli. We innovate on this prior work by quantifying human recognition with experiments, in probing invariances across model stages, by exploring a diverse set of models (e.g. those trained with self-supervision), by showing that the relevant phenomena generalize across auditory and visual domains, and by identifying many non-human invariances as model-specific. The finding that models tend to learn idiosyncratic invariances may be related to findings that the representational dissimilarity matrices for natural images can vary between individual neural network models (81).

Metamers have previously been used to validate models of visual and auditory texture (30, 82), and visual crowding (34, 35, 83, 84). A related type of model-matched stimulus has been used to test whether models can account for brain responses (58). In these previous lines of work the models instantiated hypotheses about specific types of invariances that may be present in human sensory systems, and the experiments assess human perception or brain responses to test the hypotheses. Our work differs in that the models are learned, and could in principle instantiate many different types of invariances (which are a priori unknown).

Our work also differs from prior work on metamers in that our test of model metamers is a recognition test rather than a same/different metamerism test that has classically been used for evaluating models of early sensory stages (measuring the ability to discriminate two stimuli). We view the recognition test as most appropriate for current deep neural network models that are typically trained on one or two tasks at most and that model components of high-level sensory perception, such as object or speech recognition. A same/different discrimination judgment might be performed using any representations in the observer’s perceptual system, rather than just those that are relevant for a particular task. There is thus no reason to expect that metamers for representations that support a particular task would be completely indistinguishable in every respect to a human (in contrast to metamers of color vision, for instance, for which the entire visual system is constrained by the cone inputs). The classical metamer test is thus likely to be too sensitive for our purposes – models may fail the test even if they succeeded in capturing the invariances of a particular task. We note that because an unrecognizable metamer would obviously be distinguishable from the natural image or sound to which its activations are matched, a failure in our recognition test also implies a failure in the classical test of metamerism.

### Effects of unsupervised training

One common criticism of task-optimized models is that supervised training on classification tasks is inconsistent with biological learning (48). Recent advances in unsupervised learning have enabled useful representations to be learned from large quantities of natural data without explicit labels, potentially providing a more biologically plausible computational theory of learning (52, 85, 86). A priori it seemed plausible that the invariances of deep neural network models could be strongly dependent on supervised training for classification tasks, in which case models trained without supervision might be substantially more human-like according to the metamers test. However, we found that the invariances learned from self-supervised learning also diverged from the invariances of human perceptual systems.

The metamer-related model discrepancies we saw for self-supervised models are particularly striking because these models are trained with the goal of invariance, being explicitly optimized to become invariant to the augmentations performed on the input. But the metamers of these models reveal that they nonetheless encode invariances that diverge from those of human perception. This finding is consistent with evidence that the classification errors of self-supervised models are no more human-like than those of supervised models (87). We also found that the divergence with human recognition had a similar dependence on model stage irrespective of whether models are trained with or without supervision. These findings raise the possibility that factors common to supervised and unsupervised neural networks underlie the divergence with humans.

### Differences in metamers across stages

The metamer test differs from some other model metrics (e.g. behavioral judgments of natural images or sounds, or measures of adversarial vulnerability) in that metamers can be generated from every stage of a model, with the resulting discrepancies associated with particular model stages. Because invariances reflect information that is discarded, and because information that is discarded at one stage of a feedforward system cannot be recovered, the set of metamers for a reference stimulus cannot shrink from one model stage to the next. But apart from this constraint it is otherwise not obvious how discrepancies might develop across stages, and we observed variation from model to model.

One application of this stage-specificity is to characterize intermediate model stages. In some cases metamers revealed that intermediate stages were more human-like in some models than others. For example, the effects of reducing aliasing produced large improvements in the human-recognizability of metamers from intermediate stages (Figure 6g), consistent with the idea that biological systems also avoid aliasing. By contrast, metamers from the final stages showed little improvement. This result indicates that this model change produces intermediate representations with more human-like invariances despite not resolving the discrepancy introduced at the final model stages.

For most models, the early stages produced model metamers that were fully recognizable, but that also resemble the original image or sound they were matched to. By contrast, metamers from late stages physically deviated from the original image or sound but for some models nonetheless remained recognizable. This difference highlights two ways that a model’s metamers can pass the recognition test used here – either by being perceptually indistinguishable to humans, or by being recognizable to humans as the same class despite being perceptually distinct. This distinction could be quantified in future work by combining a traditional metamer test with our recognition test.

Across all the models we considered, the final model stages tended to produce metamers that were less recognizable than natural images to humans as well as to other models. This was true irrespective of how the models were trained. This result highlights these deep stages as targets for model improvements (55).

### Limitations

Although a model that fails our metamer test is ruled out as a description of human perception, passing the test, on its own, reveals little. For instance, a model that instantiates the identity mapping would pass our test despite not being able to account for human perceptual abilities. Traditional metrics thus remain critical, but on their own are also insufficient (as shown in Figures 6 and 7). Failing the test also does not imply that the model representations are not present in the brain, only that they are not sufficient to account for the recognition behavior under consideration. For instance, there is considerable evidence for time-averaged auditory statistics in auditory perception (30, 88) even though they do not produce human-recognizable metamers for speech (Supplementary Figure 5f). We thus argue that a large suite of test metrics, including but not limited to the model metamer test, is needed for model comparison.

Model metamers are generated via iterative gradient-based optimization of a non-convex loss function, and only approximately reproduce the activations of the natural stimulus to which they are matched. We attempted to improve on previous neural network visualization work (64, 76) by setting explicit criteria for the optimization success. For models that perform classification tasks, we verified that the model metamer produced the same class label as the reference stimulus to which it was matched. Failures of the metamer test represent an unambiguous dissociation of model and human behavior for such models. We also verified that the residual error between the metamer and reference activations was much smaller than would be expected by chance, and that the residual error did not predict metamer recognizability (Supplementary Figure 4). However, the reliance on optimization may be a limitation in some contexts and with some models.

The metamer optimization process is also not guaranteed to sample uniformly from the set of a model’s metamers. Non-uniform sampling cannot explain the human-model discrepancies we observed, but could in principle contribute to differences between the magnitude of discrepancies for some models compared to others. For instance, the greater human-recognizability of metamers from some models could in principle be caused by differences in the optimization landscape that make it less likely that the metamer generation process samples along a model’s idiosyncratic invariances. We are not aware of any reason to think that this might be the case, but it is not obvious how to fully exclude this possibility.

### Adversarial training reduces idiosyncratic invariances

Why does adversarial training produce models with more recognizable metamers? One interpretation is that it prevents models from “cheating” on tasks by using data set artifacts to make decisions (61) and as a consequence develops more human-like invariances. This interpretation is supported by findings that adversarial examples often transfer across models (60, 89). Our results suggest that the effect of adversarial training on metamers likely has a distinct explanation. First, we found that not all model changes that improve adversarial robustness have comparable effects on metamer recognizability (Figure 6). In addition, robustness differences between different adversarially trained models did not predict recognizability differences between their metamers. These two findings suggest that it is the adversarial training procedure, rather than adversarial robustness per se, that underlies the increased human recognizability of model metamers. Second, metamers from standard-trained models generally did not transfer across models trained on the same data set (unlike adversarial examples). Metamers from adversarially trained models were more recognizable to humans but also transferred substantially better to other models. This finding suggests that adversarial training serves to avoid the model-specific invariances that are otherwise learned by deep neural network models. This set of results highlights the relationship between adversarial robustness and model metamers as a promising direction for future study.

### Future directions

The underlying causes of the human-model discrepancies demonstrated here seem important to understand, in part because we will not fully understand biological sensory systems until we can replicate them in a model, and deep neural networks are currently the most promising candidates. In addition, many otherwise promising model applications, such as model-based signal enhancement (90, 91), are likely to be hindered by human-discrepant model invariances.

The supervised models we explored here were trained on a single speech or image recognition task. It is possible that the use of a single task causes models to discard information that is preserved by biological sensory systems. One possibility is that biological sensory systems do not instantiate invariances per se, in the sense of mapping multiple different inputs onto the same representation (92, 93). Instead, they might learn representations that untangle behaviorally relevant variables. For instance, a system could represent word labels and talker identity, or object identity and pose, via independent directions in a representational space. Such a representational scheme could enable invariant classification without invariant representations. Some model architectures may be able to address this hypothesis, for instance by factoring out different latent variables that, once combined, can reconstruct the input representation. But as of yet we lack methods for building such models that can support human-level recognition at scale. Training on multiple tasks could be another approach that would serve this goal. The results of Figure 8 – showing that the recognition judgments of a set of models for another model’s metamers are predictive of human recognition – suggest a way to efficiently test any model for discrepant metamers, which should facilitate evaluation of these and any other model class.

The discrepancies shown here for model metamers contrast with a growing number of examples of human-model similarities for behavioral judgments of natural stimuli. Models optimized for object recognition (94), speech recognition (5), sound localization (95), and pitch recognition (96) all exhibit qualitative and often quantitative similarities to human judgments when run in traditional psychophysical experiments with natural or relatively naturalistic stimuli. Neural network models trained in naturalistic conditions thus often match human behavior for signals that fall within their training distribution, but not for some signals derived from the model that fall outside the distribution of natural sounds and images.

Current deep neural network models are overparametrized, such that training produces one of many functions consistent with the training data. From this perspective it is unsurprising that different systems can perform similarly on natural signals while exhibiting different responses to signals outside the training distribution of natural images or sounds. And yet we nonetheless found that sensible engineering modifications succeeded in bringing models into better alignment with human invariances. These results demonstrate that divergence between human and model invariances is not inevitable, and show how metamers can be a useful metric to guide and evaluate the next generation of brain models.

## METHODS

### Model training and evaluation

Models were trained and evaluated with the PyTorch deep learning library (97), and the Robustness library (98), modified to accommodate metamer generation and auditory model training. Model code and instructions for downloading checkpoints are available online at https://github.com/jenellefeather/model_metamers_pytorch. Model architecture descriptions are provided as Supplemental Information. All models were trained on the OpenMind computing cluster at MIT using NVIDIA GPUs with a minimum of 11GB memory. In the methods sections that follow we often refer to model stages as “layers”, to be consistent with how they are named in PyTorch. “Stage” and “layer” should be taken as synonymous.

#### Image training dataset

Unless otherwise noted, all visual neural network models were trained on the ImageNet1K Large Scale Visual Recognition Challenge dataset (37). This classification task consists of 1000 classes of images with 1,281,167 images in the training set and 50,000 images in the validation set. All classes were used for training the neural network models. Accuracy on ImageNet1K task and additional training parameters are reported in Table 1.

**Table 1:**
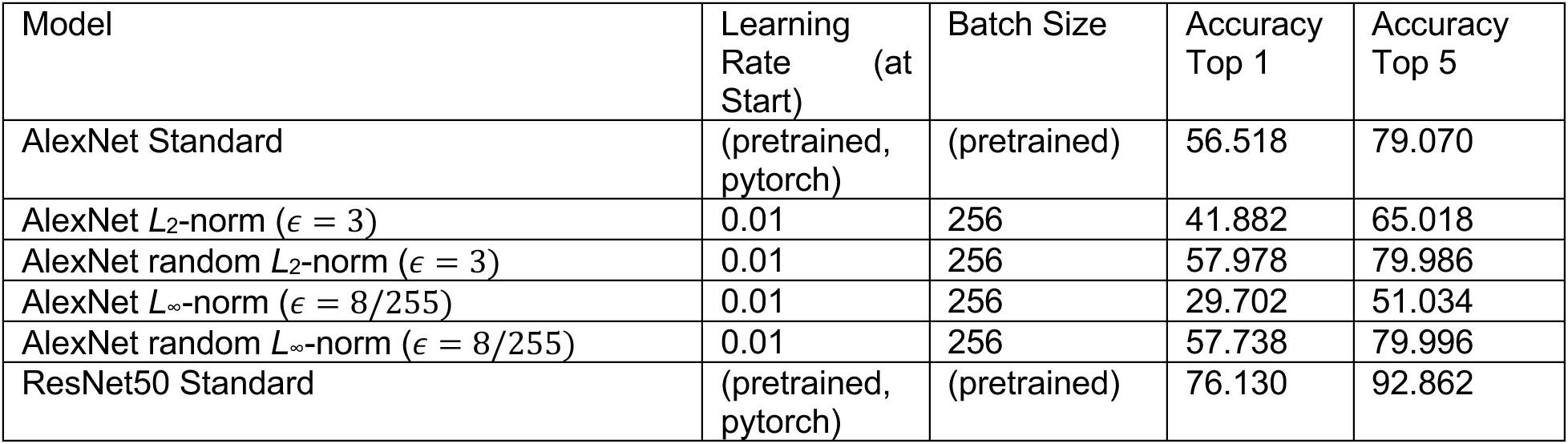
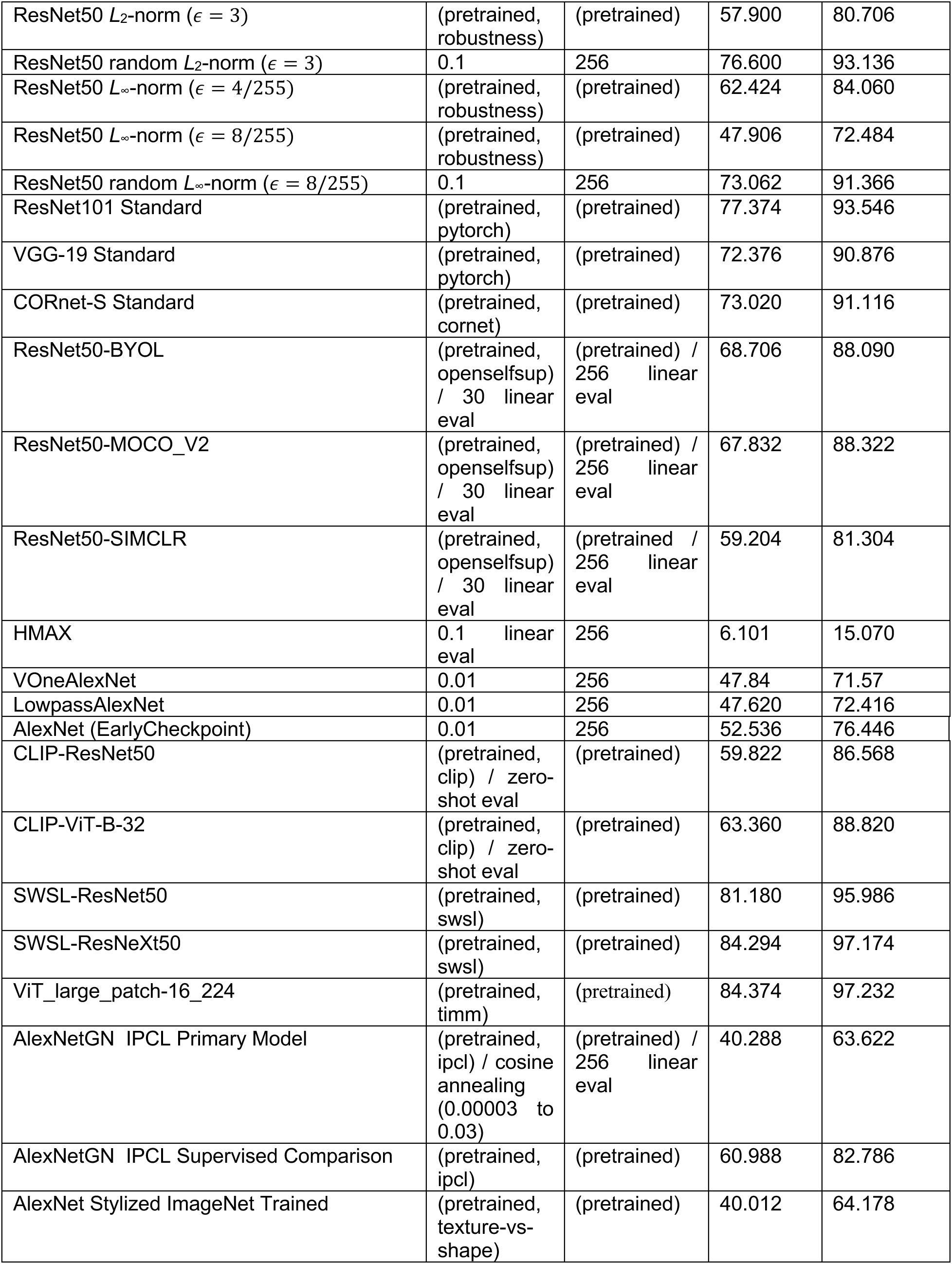
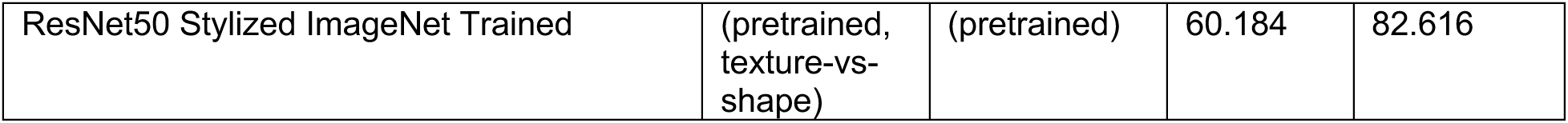
Vision architecture training parameters and ImageNet1K accuracy.

#### ImageNet1K model training and evaluation

Unless otherwise described below, visual models trained on ImageNet1K consisted of publicly available checkpoints. Standard supervised models used the pretrained PyTorch checkpoints from torchvision.models (documentation https://pytorch.org/vision/stable/models.html, referred to as “pytorch” in Table 1). Visual model performance was evaluated as the model accuracy on the ImageNet1K validation set, implemented by resizing the images so the smallest dimension was 256 pixels (or 250 in the case of HMAX) and taking a center crop of 224×224 (or 250 in the case of HMAX) pixels of the image. Train, test, and metamer images were all normalized by subtracting channel means and dividing by channel standard deviations before being passed into the first stage of the neural network backbone (except in the case of HMAX, where this normalization was not applied). Channel means were set to [0.485, 0.456, 0.406] and channel standard deviations were set to [0.229, 0.224, 0.225] unless otherwise noted for the architecture. ImageNet1K model training used data-parallelization to split batches across multiple GPUs.

#### Visual models trained on large-scale datasets

We also tested five visual models that were pretrained on datasets larger than the ImageNet1K dataset described above. Two of these were the visual encoders from Contrastive Language-Image Pre-Training (CLIP) models (one with a ResNet50 visual encoder and one with a ViT-B_32 visual encoder), obtained from the publicly available checkpoints at https://github.com/openai/CLIP (referred to as “clip” in Table 1) (43). CLIP models were trained on a dataset of 400 million (image, text) pairs collected from the internet. ImageNet1K performance from the CLIP models is evaluated with a zero-shot prediction using the list of prepared 80 image template prompts and modified ImageNet labels from (43). We found empirically that we could not synthesize metamers from this classifier, and so only included model stages from the visual encoder in our experiments. CLIP models used a custom channel mean of [0.48145466, 0.4578275, 0.40821073] and a channel standard deviation of [0.26862954, 0.26130258, 0.27577711]. The third and fourth of these models were Semi-Weakly Supervised (SWSL) ImageNet models (ResNet50 and ResNeXt101-32×8d architectures), obtained from the publicly available checkpoints at https://github.com/facebookresearch/semi-supervised-ImageNet1K-models (referred to as “swsl” in Table 1) (44). SWSL Models are pre-trained on 940 million public images with 1.5K hashtags matching with the 1000 synsets in the ImageNet dataset, followed by fine-tuning on the ImageNet1K training images described above. The fifth model was a Vision Transformer (ViT_large_patch-16_224, (45)), obtained from the publicly available checkpoint at https://github.com/rwightman/pytorch-image-models (referred to as “timm” in Table 1). ViT_large_patch-16_224 was pre-trained on the ImageNet-21K dataset, consisting of approximately 14 million images with about 21000 distinct object categories, and then fine-tuned on the ImageNet1K training images described above.

#### Self-supervised ResNet50 vision models

Self-supervised ResNet50 models were downloaded from the OpenSelfSup Model Zoo, and the training details that follow are taken from the documentation (https://github.com/open-mmlab/OpenSelfSup, referred to as “openselfsup” in Table 1). Three models, each with a ResNet50 architecture, were used: MoCo_V2, SimCLR and BYOL. MoCo_V2 self-supervised training had a batch size of 256, with data augmentations consisting of random crop (224×224 pixels), random horizontal flip (probability=0.5), random color jitter (brightness=0.4, contrast=0.4, saturation=0.4, hue=0.1, probability=0.8, all values uniformly chosen), random greyscale (probability=0.2), and random gaussian blur (sigma_min=0.1, sigma_max=0.2, probability=0.5). SimCLR self-supervised training had a batch size of 256, with augmentations consisting of random crop (224×224 pixels), random horizontal flip (probability=0.5), random color jitter (brightness=0.8, contrast=0.8, saturation=0.8, hue=0.2, probability=0.8, all values uniformly chosen), random greyscale (probability=0.2), and random gaussian blur (sigma_min=0.1, sigma_max=0.2, probability=0.5). BYOL self-supervised training had a batch size of 4096, with augmentations consisting of random crop (224×224 pixels), random horizontal flip (probability=0.5), random color jitter (brightness=0.4, contrast=0.4, saturation=0.2, hue=0.1, probability=0.8, all values uniformly chosen), random greyscale (probability=0.2), and random gaussian blur (sigma_min=0.1, sigma_max=0.2, probability=0.5). For all self-supervised ResNet50 models, a linear readout consisting of a fully connected layer with 1000 units applied to the average pooling layer of the model was trained using the same augmentations used for supervised training of the other ImageNet1K-trained models described above. The linear readout was trained for 100 epochs of ImageNet1K (while the model backbone up to avgpool remained unchanged). For MoCo_V2 and SimCLR models, the accuracy was within 1% of that reported on the OpenSelfSup (BYOL average pooling evaluation was not posted at the time of training). The linear readout served as a check that the downloaded models were instantiated correctly, and was used to help verify the success of the metamer generation optimization procedure, as described below. Linear evaluations from each model stage were obtained by training a fully connected layer with 1000 units and a softmax classifier applied to a random subsample of 2048 activations. If the number of activations was less than or equal to 2048 all activations were maintained.

#### Self-supervised IPCL AlexNetGN models

A model trained with Instance-Prototype Contrastive Learning (IPCL) and a supervised model with the same augmentations were downloaded from the IPCL github (https://github.com/harvard-visionlab/open_ipcl, referred to as “ipcl" in Table 1) (52). Both models used an AlexNet architecture with group-normalization layers. As described in the original publication (52), the models were trained with a batch size of 128×5 (128 images with 5 augmentations each), for 100 epochs, with data augmentations consisting of a random resize crop (random crop of the image resized with a scale range of [0.2,1] and aspect ratio [3/4,4/3], and resized to 224×224 pixels), random horizontal flip (probability=0.5), random grayscale conversion (probability=0.2), random color jitter (brightness=0.6, contrast=1, saturation=0.4, and hue +/-144 degrees).

Using the procedure described in (52), a linear readout consisting of a fully connected layer with 1000 units was used to evaluate each model stage. The readout was trained using a batch size of 256, with input augmentations of a random resize crop (random crop of the image resized with a scale range of [0.08,1] and aspect ratio [3/4,4/3], and resized to 224×224 pixels) and a random horizontal flip (probability=0.5). The linear readout was trained for 10 epochs with the one-cycle learning rate policy (99), with cosine annealing to vary the learning rate from 0.00003, increasing to a maximum of .3 after 3 epochs, then decreasing with a cosine annealing function toward zero (3e-09) by 10 epochs.

#### Models trained on Stylized ImageNet

Models trained on a “Stylized” ImageNet were downloaded from publicly available checkpoints (https://github.com/rgeirhos/texture-vs-shape, referred to as “texture-vs-shape” in Table 1) and training details that follow are taken from the documentation (53). Stylized ImageNet is constructed by taking the content of an ImageNet1K image and replacing the style of the image with that of a randomly selected painting using AdaIN style transfer (100). A single stylized version of each image in the ImageNet1K training dataset was used for training. The models were trained with a batch size of 256 for 60 epochs, with input augmentations of a random resize crop (random crop of the image resized with a scale range of [0.08,1] and aspect ratio [3/4,4/3], and resized to 224×224 pixels) and a random horizontal flip (probability=0.5).

#### HMAX vision model

The hand-engineered HMAX vision model was based off of a publicly availably implementation in PyTorch (https://github.com/wmvanvliet/pytorch_hmax) which follows the model documented in a previous publication (4). A gaussian activation function was used, and boundary handling was added to match the MATLAB implementation provided by the original HMAX authors (https://maxlab.neuro.georgetown.edu/hmax.html). For full comparison to the other models, we trained a linear classifier consisting of 1000 units to perform the ImageNet1K recognition task on the final C2 output of the HMAX model. This fully connected layer was trained for 30 epochs of the ImageNet1K training dataset, and the learning rate was dropped after every 10 epochs. Inputs to HMAX during the classifier training consisted of random crops (250×250 pixels), random horizontal flip (p=0.5), random color jitter (brightness=0.1, contrast=0.1, saturation=0.1, probability=1, all values uniformly chosen), and lighting noise (alpha standard deviation of 0.05, an eigenvalue of [0.2175, 0.0188, 0.0045], and channel eigenvectors of [[-0.5675, 0.7192, 0.4009], [-0.5808, -0.0045, -0.8140], [-0.5836, -0.6948, 0.4203]]). HMAX performance was evaluated by measuring the model accuracy on the ImageNet1K validation set after resizing the images so that the smallest dimension was 250 pixels, taking a center crop of 250×250 pixels of the image, converting to greyscale, and multiplying by 255 to scale the image to the 0-255 range. As expected, the performance on this classifier was low, but it was significantly above chance and could thus be used for the metamer optimization criteria described below.

#### Adversarial training – vision models

Adversarially trained ResNet50 models were obtained from the robustness library (https://github.com/MadryLab/robustness, referred to as “robustness” in Table 1). Adversarially trained AlexNet architectures and the random perturbation ResNet50 and AlexNet architectures were trained for 120 epochs of the ImageNet1K dataset, with image pixel values scaled between 0-1, using data parallelism to split batches across multiple GPUs. Learning rate was decreased by a factor of 10 after every 50 epochs of training. During training, data augmentation consisted of random crop (224×224 pixels), random horizontal flip (probability=0.5), color jitter (brightness=0.1, contrast=0.1, saturation=0.1, probability=1, all values uniformly chosen), and lighting noise (alpha standard deviation of 0.05, an eigenvalue of [0.2175, 0.0188, 0.0045], and channel eigenvectors of [[-0.5675, 0.7192, 0.4009], [-0.5808, -0.0045, -0.8140], [-0.5836, -0.6948, 0.4203]]). An adversarial or random perturbation was then added. All adversarial examples were untargeted, such that the loss used to generate the adversarial example pushed the input away from the original class by maximizing the cross-entropy loss, but did not push the prediction towards a specific target class. For the *L*2-norm (є = 3) model, adversarial examples were generated with a step size of 1.5 and 7 attack steps. For the *L*∞-norm (є = 8/255) model, adversarial examples were generated with a step size of 4/255 and 7 attack steps. For both ResNet50 and AlexNet random-perturbation *L*2-norm models, a random sample on the *L*2 ball with width є = 3 was drawn and added to the input, independently for each training example and dataset epoch. Similarly, for both Resnet50 and AlexNet random perturbation *L*∞-norm models, a random sample on the corners of the *L*∞ ball was selected by randomly choosing a value of ±8/255 to add to each image pixel, independently chosen for each training example and dataset epoch. After the adversarial or random perturbation was added to the input image, the new image was clipped between 0-1 before being passed into the model.

#### VOneAlexNet vision model

The VOneAlexNet architecture was downloaded from the VOneNet GitHub repository (https://github.com/dicarlolab/vonenet) (67). Modifications were then made to use Gaussian noise rather than Poisson noise as the stochastic component, as in (68), and to use the same input normalization as in our other models (rather than a mean of 0.5 and standard deviation of 0.5 as used in (67)). The VOneAlexNet architecture was trained for 120 epochs using the same data augmentations and training procedure described for the adversarially trained AlexNet model (but without adversarial or random pertubrations). The model was trained with stochastic responses (Gaussian noise with standard deviation of 4) in the “VOne” model stage, but for the purposes of metamer generation we fixed the noise by randomly drawing one noise sample when loading the model and using this noise sample for all metamer generation and adversarial evaluation. Although “fixing” the noise reduces the measured adversarial robustness compared to when a different sample of noise is used for each iteration of the adversarial example generation, the model with a single noise draw was still significantly more robust than a standard model, and allowed us to perform the metamer experiments without having to account for the stochastic representation during metamer optimization.

#### LowpassAlexNet vision model

The LowpassAlexNet architecture was trained for 120 epochs using the same augmentations and training procedure described for the adversarially trained AlexNet models (but without adversarial or random perturbations). To approximately equate performance on natural stimuli with the VOneNetAlexNet, we chose an early checkpoint that was closest, but did not exceed, the Top 1% performance of the VOneAlexNet model (to ensure that the greater recognizability of the metamers from LowpassAlexNet could not be explained by higher overall performance of that model). This resulted in a comparison model trained for 39 epochs of the ImageNet1K dataset.

#### AlexNet vision model, early checkpoint

We trained an AlexNet architecture for 120 epochs using the same augmentations and training procedure described for the adversarially trained AlexNet models (but without adversarial or random perturbations). After training, to approximately equate performance on natural stimuli with the VOneNetAlexNet and LowpassAlexNet, we chose an early checkpoint that was closest, but not lower than, the performance of the VOneAlexNet model. This resulted in a comparison model trained for 51 epochs of the ImageNet1K dataset.

#### Pre-trained adversarially robust models for final stage evaluation

To compare the relationship between metamer recognizability and adversarial robustness of adversarially trained models (Figure 6c), we evaluated a large set of adversarially trained models. We evaluated all of the models included in a well-known robustness evaluation as of November 2022 – these comprised the ImageNet- *L*∞ evaluation of robustbench (65) (8 models), as well as additional models from each of the repositories from which these models were chosen (17 additional models), for a total of 25 models. Five of these models were from https://github.com/dedeswim/vits-robustness-torch (101), and were trained with *L*∞-norm, є = 4/255, (ViT-XCiT-L12, ViT-XCiT-M12, ViT-XCit-S12, ConvNeXt-T, and GELUResNet-50). Another 16 of these models were from https://github.com/microsoft/robust-models-transfer (102), with 3 trained with *L*∞-norm, є = 4/255, (ResNet18, ResNet50, and Wide-ResNet-50-2), 3 trained with *L*∞-norm, є = 1/255, (ResNet18, ResNet50, and Wide-ResNet-50-2), and 10 trained with *L*2- norm, є = 3.0, (ResNet18, ResNet50, Wide-ResNet-50-2, Wide-ResNet-50-4, DenseNet, MNASNET, MobileNet-v2, ResNeXt50_32×4d, ShuffleNet, VGG16_bn). Another 3 models were ResNet50 architectures from https://github.com/MadryLab/robustness (98); one was trained on *L*∞-norm, є = 4/255, one was trained on *L*∞-norm, є = 8/255, and was one trained on *L*2- norm, є = 3.0). Lastly, 1 model was the ResNet50 checkpoint available from https://github.com/locuslab/fast_adversarial (103). Only the final layer was used for metamer generation for these models. These models are omitted from Table 1 as they were used for only a single analysis (that of Figure 6b).

#### Adversarial evaluation -- visual models

The adversarial robustness of visual models was evaluated with white-box untargeted adversarial attacks (i.e., in which the attacker has access to the model’s parameters when determining an attack that will cause the model to classify the image as any category other than the correct one). All 1000 classes of ImageNet1K were used for the adversarial evaluation. Attacks were computed with *L1, L2, and L*∞ maximum perturbation sizes (ε) added to the image, with 64 gradient steps each with size ε/4 (pilot experiments suggested that this step size and number of steps were sufficient to produce adversarial examples for most models). We randomly chose images from the ImageNet1K evaluation dataset to use for adversarial evaluation, applying the evaluation augmentation described above (resizing so that the smallest dimension was 256 pixels, followed by a center crop of 224×224 pixels). Five different subsets of 1024 stimuli were drawn to compute error bars.

For the detailed investigation of adversarial vulnerability shown in Supplemental Figure 9, we measured robustness to two additional types of white-box adversarial attacks. “Fooling Images” (23) were constructed by first initializing the input image as a sample from a normal distribution with standard deviation of 0.05 and a mean of 0.5. We then randomly chose a target label from the 1000 classes of ImageNet1K and derived a perturbation to the image that would cause the noise to be classified as the target class. Performance was evaluated as the percent of perturbed images that had the target label. Attacks were computed with *L1, L2, and L*∞ maximum perturbation sizes (ε) added to the image, with 64 gradient steps each with size ε/4. Error bars were computed using five different random samples of 1024 target labels. “Feature Adversaries” (69) were constructed by deriving small perturbations to a natural “source” image to yield model activations (at a particular model stage) that are close to those evoked by a different natural “target” image, by minimizing the L2 distance between the perturbed source image activations and the target activations. The source and target images were randomly selected from the ImageNet1K validation dataset. Evaluation was performed by measuring the percent of perturbed images that had the same label as the target image. Attacks were computed with *L1, L2, and L*∞ maximum perturbation sizes (ε) added to the image, with 128 gradient steps each with size ε/16. Error bars were computed using five different subsets of 1024 “source” and “target” stimuli.

For the statistical comparisons between the adversarial robustness of architectures for Figure 6f we performed a repeated measure ANOVA with within-group factors of architecture and perturbation size !. A separate ANOVA was performed for each adversarial attack type. The values of є included in the ANOVA were constrained to a range where the VOneAlexNet and LowPassAlexNet showed robustness over the standard AlexNet (four values for each attack type,, so that any difference in clean performance did not affect the comparisons. We computed statistical significance for the main effect of architecture by a permutation test, randomly permuting the architecture assignment, independent for each subset of the data. We computed a p-value by comparing the observed F-statistic to the null distribution of F-statistics from permuted data (i.e., the p-value was one minus the rank of the observed F-statistic divided by the number of permutations). In cases where the maximum possible number of unique permutations was less than 10,000, we instead divided the rank by the maximum number of unique permutations.

We performed the same type of ANOVA analysis for the statistical comparisons in Supplementary Figure 9, using adversarial attack strengths for which VOneAlexNet and LowPassAlexNet showed robustness over the standard AlexNet for the specific attack being evaluated. For the “Fooling Images” in Supplementary Figure 9a we used attack strengths of *ε_L1_ ε{10^1.5^, 10^2^, 10^2.5^, 10^3^}, *ε_L2_* *ε*{10^-1^, 10^-0.5^, 10^2^, 10^0.5^}, *ε_L1_*{10^-3.5^, 10^-3^, 10^-2.5^, 10^-2^}, and for the feature adversaries in Supplementary Figure 9b we used attack strengths of *ε_L1_*{10^-2.5^, 10^3^, 10^-3.5^, 10^4^}, *ε*_L2_ ε*{10^0^, 10^0.5^, 10^1^, 10^1.5^}, ε{10^-2.5^, 10^-2^, 10^-1.5^, 10^-1^}). Supplementary Figure 9b we used attack strengths of *ε_L1_*{10^-2.5^, 10^-3^, 10^-3.5^, 10^-4^}, *ε_L2_* ε {10^0^, 10^0.5^, 10^1^, 10^1.5^}, *ε_L2_* ε {10^-2.5^, 10^-2^, 10^-1.5^, 10^-1^}

#### Out of distribution evaluation – visual models

We evaluated model performance on out of distribution images using two publically available benchmarks. For both benchmarks, we utilized the BrainScore (42) implementations of the behavioral benchmarks for the models.

The ImageNet-C benchmark (70) measures model top 1 accuracy on distorted images derived from the ImageNet test set, using the labels for the original image. The accuracy is averaged over stimulus sets consisting of the following distortions: Gaussian noise, shot noise, impulse noise, defocus blur, frosted glass blur, motion blur, zoom blur, snow, frost, fog, brightness, contrast, elastic, pixelate, and JPEG compression.

The Geirhos 2021 benchmark (72) measures whether the model gets the same stimuli correct as human observers (error consistency), using 16-way recognition decisions from human observers for comparison. The error consistency is averaged over stimulus sets consisting of the following stimulus manipulations: colour/grayscale, constrast, high-pass, low-pass (blurr), phase scrambling, power equalization, false colour, rotation, Eidolon I, Eidolon II, Eidonlon III, uniform noise, sketch, stylized, edge, silhouette, and texture-shape cue conflict.

#### Audio training dataset

All auditory neural network models were trained on the Word-Speaker-Noise (WSN) dataset. This dataset was first presented in (36) and was constructed from existing speech recognition and environmental sound classification datasets. The dataset is approximately balanced to enable performance of three tasks on the same training exemplar: (1) recognition of the word at the center of a two second speech clip (2) recognition of the speaker and (3) recognition of environmental sounds, that are superimposed with the speech clips (serving as “background noise” for the speech tasks while enabling an environmental sound recognition task). Although the dataset is constructed to enable all three tasks, the models described in this paper were only trained to perform the word recognition task. The speech clips used in the dataset were excerpted from the Wall Street Journal (104) (WSJ) and Spoken Wikipedia (105) (SWC).

To choose speech clips, we screened WSJ, TIMIT (106) and a subset of articles from SWC for appropriate audio clips (specifically, clips that contained a word at least four characters long and that had one second of audio before the beginning of the word and after the end of the word, to enable the temporal jittering augmentation described below). Some SWC articles were left out of the screen due to a) potentially offensive content for human listening experiments; (29/1340 clips), b) missing data; (35/1340 clips), or c) bad audio quality (for example, due to computer generated voices of speakers reading the article or the talker changing mid-way through the clip; 33/1340 clips). Each segment was assigned the word class label of the word overlapping the segment midpoint and a speaker class label determined by the speaker. With the goal of constructing a dataset with speaker and word class labels that were approximately independent, we selected words and speaker classes such that the exemplars from each class spanned at least 50 unique cross-class labels (e.g., 50 unique speakers for each of the word classes). This exclusion fully removed TIMIT from the training dataset. We then selected words and speaker classes that each contained at least 200 unique utterances, and such that each class could contain a maximum of 25% of a single cross-class label (e.g., for a given word class, a maximum of 25% of utterances could come from the same speaker). These exemplars were subsampled so that the maximum number in any word or speaker class was less than 2000. The resulting training dataset contained 230,356 unique clips in 793 word classes and 432 speaker classes, with 40,650 unique clips in the test set. Each word class had between 200 and 2000 unique exemplars. A “null” class was used as a label when a background clip was presented without the added speech.

The environmental soundtrack clips that were superimposed on the speech clips were a subset of examples from the AudioSet dataset (a set of annotated YouTube video soundtracks) (107). To minimize ambiguity for the two speech tasks, we removed any sounds under the “Speech” or “Whispering” branch of the AudioSet ontology. Since a high proportion of AudioSet clips contain music, we achieved a more balanced set by excluding any clips that were only labeled with the root label of “Music”, with no specific branch labels. We also removed silent clips by first discarding everything tagged with a “Silence” label and then culling clips containing more than 10% zeros. This screening resulted in a training set of 718,625 unique natural sound clips spanning 516 categories. Each AudioSet clip was a maximum of 10 seconds long, from which a 2-second excerpt was randomly cropped during training (see below).

#### Auditory model training

During training, the speech clips from the Word-Speaker-Noise dataset were randomly cropped in time and superimposed on random crops of the AudioSet clips. Data augmentations during training consisted of 1) randomly selecting a clip from AudioSet to pair with each labeled speech clip, 2) randomly cropping 2 seconds of the AudioSet clip and 2 seconds of the speech clip, cropped such that the labeled word remained in the center of the clip (due to training pipeline technicalities, we used a pre-selected set of 5,810,600 paired speech and natural sound crops which spanned 25 epochs of the full set of speech clips and 8 passes through the full set of AudioSet clips), 3) superimposing the speech and the noise (i.e., the AudioSet crop) with a Signal-to-Noise-Ratio (SNR) sampled from a uniform distribution between -10dB SNR and 10dB SNR, augmented with additional samples of speech without an AudioSet background (i.e. with infinite SNR, 2464 examples in each epoch) and samples of AudioSet without speech (i.e. with negative infinite SNR, 2068 examples in each epoch) and 4) setting the root-mean-square (RMS) amplitude of the resulting signal to 0.1. Evaluation performance is reported on one pass through the speech test set (i.e., one crop from each of the 40,650 unique test set speech clips) constructed with the same augmentations used during training (specifically, variable SNR and temporal crops, paired with a separate set of AudioSet test clips, same random seed used to test each model such that test sets were identical across models). Audio model training used data-parallelization to split batches across multiple GPUs.

Each auditory model was trained for 150 epochs (where an epoch is defined as a full pass through the set of 230,356 speech training clips). The learning rate was decreased by a factor of 10 after every 50 epochs (see Table 2).

**Table 2:**
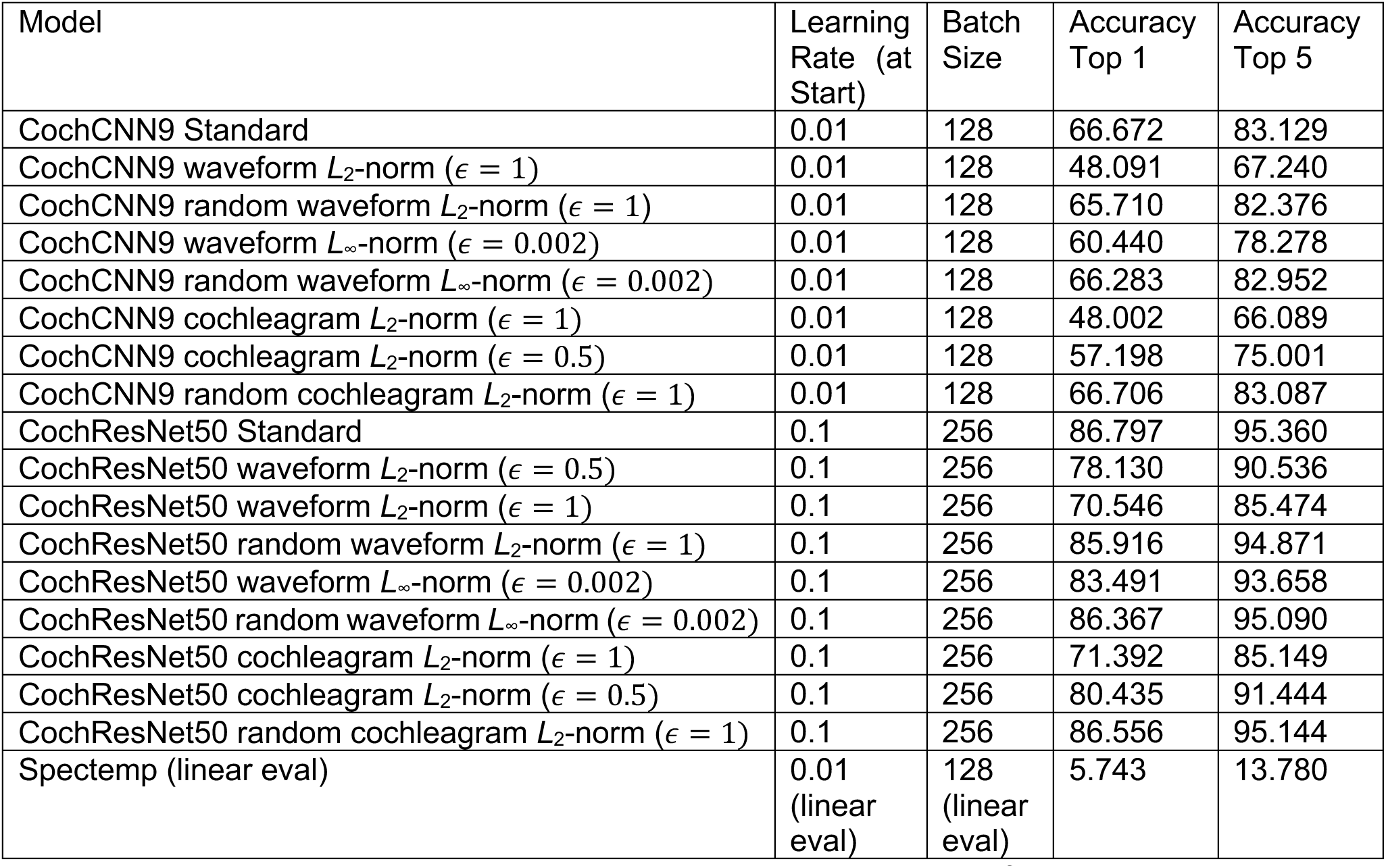
Auditory model architecture training parameters and word classification accuracy.

#### Auditory model cochlear stage

The first stage of the auditory models produced a “cochleagram” – a time-frequency representation of audio with frequency tuning that mimics the human ear, followed by a compressive nonlinearity (47). This stage consisted of the following sequence of operations. First, the 20kHz audio waveform passed through a bank of 211 bandpass filters with center frequencies ranging from 50Hz to 10kHz. Filters were zero-phase with frequency response equal to the positive portion of a single period of a cosine function, implemented via multiplication in the frequency domain. Filter spacing was set by the Equivalent Rectangular Bandwidth (ERB N) scale (46). Filters perfectly tiled the spectrum such that the summed squared response across all frequencies was flat (four low-pass and four high-pass filters were included in addition to the bandpass filters in order to achieve this perfect tiling). Second, the envelope was extracted from each filter subband using the magnitude of the analytic signal (via the Hilbert transform). Third, the envelopes were raised to the power of 0.3 to simulate basilar membrane compression. Fourth, the compressed envelopes were lowpass-filtered and downsampled to 200Hz (1d convolution with a Kaiser-windowed Sinc filter of size 1001 in the time domain, applied with a stride of 100 and no zero padding, i.e. “valid” convolution), resulting in a final “cochleagram” representation of 211 frequency channels by 390 time points. The first stage of the neural network “backbone” of the auditory models operated on this cochleagram representation. Cochleagram generation was implemented in PyTorch such that the components were differentiable for metamer generation and adversarial training. Cochleagram generation code will be released upon acceptance of the paper.

#### Spectemp model

The hand-engineered Spectro-Temporal filter model (Spectemp) was based on a previously published model (56). Our implementation differed from the original model in specifying spectral filters in cycles/ERB rather than cycles/octave (because our implementation operated on a cochleagram generated with ERB-spaced filters). The model consisted of a linear filter bank tuned to spectro-temporal modulations at different frequencies, spectral scales, and temporal rates. The filtering was implemented via 2D convolution with zero padding in frequency (211 samples) and time (800 samples). Spectro-temporal filters were constructed with spectral modulation center frequencies of [0.0625, 0.125, 0.25, 0.5, 1, 2] cycles/ERB and temporal modulation center frequencies of [0.5, 1, 2, 4, 8, 16, 32, 64] Hz, including both upward and downward frequency modulations (resulting in 96 filters). An additional 6 purely spectral and 8 purely temporal modulation filters were included for a total of 110 modulation filters. This filterbank operated on the cochleagram representation (yielding the ‘filtered_signal’ stage in Figure 4d-f). We squared the output of each filter response at each time step (‘power’) and took the average across time for each frequency channel (‘average’), similar to previous studies (5, 57, 74). To be able to use model classification judgments as part of the metamer generation optimization criteria (see below), we trained a linear classifier after the average pooling layer (trained for 150 epochs of the speech training set with a learning rate that started at 0.01 and decreased by a factor of 10 after every 50 speech epochs, using the same data augmentations as for the neural networks). Although performance on the word recognition task for the Spectemp model was low, it was significantly above chance, and thus could be used to help verify the success of the metamer generation optimization procedure.

#### Adversarial training – auditory models – waveform perturbations

CochResNet50 and CochCNN9 were adversarially trained with perturbations in the waveform domain. We also included a control training condition in which random perturbations were added to the waveform. For both adversarial and random waveform perturbations, after the perturbation was added, the audio signal was clipped to fall between -1 and 1. As with the adversarially trained vision models, all adversarial examples were untargeted. The *L*2-norm (!= 0.5 and є = 1.0) model adversarial examples were generated with a step size of 0.25 and 0.5, respectively, and 5 attack steps. *L*∞-norm (є = 0.002) model adversarial examples in the waveform space were generated with a step size of 0.001 and 5 attack steps. For random perturbation *L*2-norm models (both CochResNet50 and CochCNN9), a random sample on the *L*2 ball with width є = 1.0 was selected and added to the waveform, independently for each training example and dataset epoch. Similarly, for random perturbation *L*∞-norm models, a random sample on the corners of the *L*∞ ball was selected by randomly choosing a value of ±0.002 to add to each image pixel, chosen independently for each training example and dataset epoch.

We estimated the SNR_*+_ for the perturbations of the waveform using:

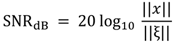

where x is the input waveform and ξ is the adversarial perturbation. As described above, the input waveforms, x, to the model were RMS normalized to 0.1, and thus ||:|| = 0.1 ∗ √?, where *n* is the number of samples in the waveform (40,000). For *L*2-norm perturbations to the waveform, the norm of the perturbation is just the є value, and so є = 0.5 and є = 1.0 correspond to ||ξ|| = 0.5 and ||ξ|| = 1, resulting in SNR*+ values of 32.04 and 26.02, respectively. For *L*∞-norm perturbations, the worst case (lowest) SNR_*+_ is achieved by a perturbation that maximally changes every input value. Thus, an *L*∞ perturbation with є = 0.002 has ||ξ|| = 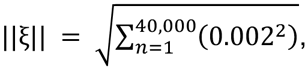 corresponding to a SNR_*+_ value of 33.98. These SNR_*+_ values do not guarantee that the perturbations were always fully inaudible to humans, but they confirm that the perturbations are relatively minor and unlikely to be salient to a human listener.

#### Adversarial training – auditory models – cochleagram perturbations

CochResNet50 and CochCNN9 were adversarially trained with perturbations in the cochleagram domain. We also included a control training condition in which random perturbations were added to the cochleagram. Adversarial or random perturbations were added to the output of the cochleagram stage, after which the signal was passed through a ReLU so that no negative values were fed into the neural network backbone. All adversarial examples were untargeted. The *L*2-norm (є = 0.5 and є = 1.0) model adversarial examples were generated with a step size of 0.25 and 0.5 respectively, and 5 attack steps. For random perturbation *L*2-norm models (both CochResNet50 and CochCNN9), a random sample on the *L*2 ball with width є = 1.0 was selected, independently for each training example and dataset epoch.

We estimated the SNR_*+_ of the cochleagram perturbations using the average cochleagram from the test dataset, whose *L*_2_-norm was 40.65. Using this value with the SNR*+ equation yielded estimates of 38.20 and 32.18 dB for cochleagram perturbation models trained with є = 0.5 and є = 1.0, respectively. We again cannot guarantee that the perturbations are inaudible to a human, but they are fairly low in amplitude and thus unlikely to be salient.

#### Adversarial evaluation – auditory models

As in visual adversarial evaluation, the adversarial vulnerability of auditory models was evaluated with untargeted white-box adversarial attacks. Attacks were computed with *L_1_, L_2_, and L*_∞_ maximum perturbation sizes (ε) added to the waveform, with 64 gradient steps each with size ε/4 (pilot experiments and previous results (68) suggested that this step size and number of steps were sufficient to attack most auditory models). We randomly chose audio samples from the WSN evaluation dataset to use for adversarial evaluation, including the evaluation augmentations described above (additive background noise augmentation with SNR randomly chosen between -10 to 10 dB SNR, and RMS normalization to 0.1). Five different subsets of 1024 stimuli were drawn to compute error bars.

#### Metamer generation

##### Optimization of metamers

Gradient descent was performed to minimize the normalized squared error between all activations at a particular model stage (for instance, each *x, y,* and *channel* value from the output of a convolutional layer) for the model metamer and the corresponding activations for a natural signal:

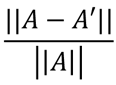

where *A* represents the activations from the natural signal and *A’* represents the activations from the model metamer. The weights of the model remained fixed during the optimization. Optimization was performed with gradient descent on the input signal, which was otherwise unconstrained. Each step of gradient descent was constrained to have a maximum *L2* norm of η, where η was initialized at 1 and was dropped by a factor of 0.5 after every 3000 iterations. Optimization was run for a total of 24000 steps for each generated metamer. For all vision models other than HMAX, the input signal was initialized as a sample from a normal distribution with standard deviation of 0.05 and a mean of 0.5 (natural inputs to the models were scaled between 0-1). The size of the input stimuli, range of pixel values, and normalization parameters were matched to the test data for the model. For all auditory models other than the spectrotemporal model, the input signal was initialized from a random normal distribution with standard deviation of 10^-7^ and a mean of zero. The perturbation *s* was initialized to 0. Normalization that occurred after data augmentation in the visual models (subtracting channel means and dividing by channel standard deviations) was included as a model component during metamer generation (i.e., gradients from these operations contributed to the metamer optimization along with all other operations in the model).

The two hand-engineered models (HMAX and the Spectemp model) had different initialization settings. For the HMAX model, whose inputs were scaled to fall between 0-255, metamers were initialized with a random normal distribution with a mean of 127.5 and standard deviation of 10. For the Spectemp model, metamers were initialized with a random normal distribution with a mean of zero and standard deviation of 10^-5^. Empirically, we found that both of these models contained stages that were difficult to optimize with the same strategy used for the neural networks. For both models, we maintained the learning rate schedule of 24000 total optimization steps with learning rate drops of 0.5 after every 3000 iterations (initialized at a learning rate of 1). However, in both cases we found that optimization was aided by selectively optimizing for subsets of the units (channels) in the early iterations of the optimization process. For the HMAX model, the subsets that were chosen depended on the model stage. For the S1 stage, we randomly choose activations from a single Gabor filter channel to include in the optimization. For the C1 stage, we randomly selected a single scale. And for the S2 and C2 stages we randomly chose a single patch size. The random choice of subset was changed after every 50 gradient steps. This subset-based optimization strategy was used for the first 2000 iterations at each learning rate value. All units were then included for the remaining 1000 iterations for that learning rate value.

For the Spectemp model, we observed that the higher frequency modulation channels were hardest to optimize. We set up a coarse-to-fine optimization strategy by initially only including the lowest frequency spectral and temporal modulation filters in the loss function, and adding in the filters with the next lowest modulation frequencies after every 400 optimization steps (with 7 total sets of filters defined by center frequencies in both temporal and spectral modulation, and the remaining 200/3000 steps continuing to include all of the filters from the optimization). The temporal modulation cutoffs for each of the 7 sets were [0, 0.5, 1, 2, 4, 8, 16] Hz and the spectral modulation cutoffs were [0, 0.0625, 0.125, 0.25, 0.5, 1, 2] cycles/ERB; a filter was included in the n^th^ set if it had either a temporal or spectral scale that was equal to or less than the n^th^ temporal or spectral cutoff, respectively. This strategy was repeated for each learning rate.

#### Criteria for optimization success

Because metamers are derived via a gradient descent procedure, the activations that they produce approach those of the natural signal used to generate them, but never exactly match. It was thus important to define criteria by which the optimization would be considered sufficiently successful for the result to be considered a model metamer and included in the behavioral experiments.

The first criterion was that the models had to produce the same class label for the model metamer and natural signal. For visual models, the model metamer had to result in the same 16-way classification label as the natural signal to which it was matched. For the auditory models, the model metamer had to result in the same word label (out of 794 possible labels, including “null”) as the natural speech signal to which it was matched. For models that did not have a classifier stage (the self-supervised models, HMAX, and the spectrotemporal filter model), we trained a classifier as described above for this purpose. The classifier was included to be conservative, but in practice could be omitted in future work, as very few stimuli pass the matching fidelity criterion but not the classifier criterion.

The second criterion was that the activations for the model metamer had to be matched to those for the natural signal better than would be expected by chance. We measured the fidelity of the match between the activations for the natural stimulus and its model metamer at the matched model stage using three different metrics: Spearman ρ, Pearson R^2^, and the Signal-To-Noise Ratio:

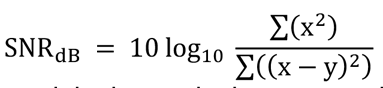

where *x* is the activations for the original sound when comparing metamers, or for a randomly selected sound for the null distribution and *y* is activations for the comparison sound (the model metamer or another randomly selected sound). We then ensured that for each of the three measures, the value for the model metamer fell outside of a null distribution measured between 1,000,000 randomly chosen image or audio pairs from the training dataset. Metamers that did not pass the null distribution test for any of the Spearman M, Pearson R^2^, or Signal-To-Noise Ratio measured at the stage used for the optimization were excluded from the set of experimental stimuli. The only exception to this was the HMAX model, for which we only used the Signal-To-Noise Ratio for the matching criteria (we found empirically that after the S2 stage of the HMAX model, activations from pairs of random images became strongly correlated due to the different offsets and scales in the natural image patch set, such that the correlation measures were not diagnostic of the match fidelity).

#### Handling gradients through the ReLU operation

Many neural networks use the ReLU nonlinearity, which yields a partial derivative of zero if the input is negative. We found empirically that it was difficult to match ReLU layers due to the initialization producing many activations of zero. To improve the optimization when generating a metamer for activations immediately following a ReLU, we modified the derivative of the metamer generation layer ReLU to be 1 for all input values, including values below zero (36). ReLU layers that were not the metamer generation layer behaved normally, with a gradient of 0 for input values below zero.

#### Metamer generation with regularization

To investigate the effects of regularization on metamer recognizability, in one experiment we generated metamers with additional constraints on the optimization procedure. We followed the procedures of an earlier paper by Mahendran and Vedaldi (64). Two regularization terms were included: (1) a total variation (TV) regularizer, and (2) an ɑ-norm regularizer.

The resulting objective function minimized to generate metamers was:

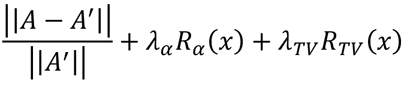

using the 6-norm for the alpha-norm regularizer:

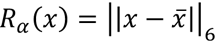

and using the TV regularizer:

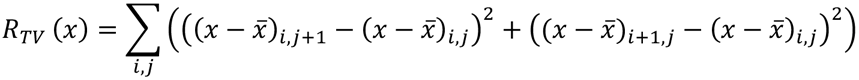

where A are the activations evoked by the natural signal at the generation layer, A’ are the activations evoked by the model metamer at the generation layer, *λ_α_ λ_TV_* and are scaling coefficients for the regularizers, and :̅ is the mean of the input signal : (: is normalized according to the typical normalization for the model, ie subtracting the dataset mean and dividing by the dataset standard deviation for each channel).

For the TV regularizer we generated metamers for 3 different coefficient values with O_12"_=0.000005, O_12%_ =10O_12"_, and O_12&_ =100 O_12"_. As was also observed by Mahendran and Vedaldi (64), we found that larger TV regularizers impaired optimization at early model stages, with resulting stimuli often not passing our metamer optimization success criteria (for instance, only 2/400 metamers generated from relu_0 of AlexNet passed this criteria for the largest regularizer coefficient value). Thus, for the behavioral experiments that we ran, we chose separate coefficient values for each model stage. Specifically, in AlexNet we used O_12"_ for relu_0, and relu_1, O_12%_ for relu_2 and relu_3 and O_12&_ for relu_4, fc_0, fc_1, and final (this is exactly what was done in (64) except that the lambda values are different due to differences in how the input is normalized, 0-255 in (64) compared to 0-1 in our models).

For the Y-norm regularizer, we followed the methods used in (64), with Y=6, and using a single coefficient of O_0_ =0.005 for all stages. This coefficient was chosen based on the logic proposed in (64) for the starting value, with a small sweep around the values for a small number of examples (10x up and 10x down), in which we subjectively judged which value produced the largest visual recognizability benefit.

We also observed that when these regularizers were used, the default step sizes (initial learning rate η = 1) used in our metamer generation method resulted in stimuli that looked qualitatively more “gray” than expected, i.e. stayed close to the mean. Thus, to maximize the chances of seeing a benefit from the regularization, in a separate condition we increased the initial step size for metamer generation to be 16x the default value (initial η = 16).

We note that regularization typically interferes with the generation of metamers. We found empirically that there was a tradeoff between satisfying the goal of matching the metamer activations and of minimizing the regularization term. Imposing the regularization generally resulted in worse metamer optimization. As described above, it was necessary to hand-tune the

regularization weights to obtain something that met our convergence criteria, but even when these criteria were met, metamers generated with regularization tended to have worse activation matches than metamers generated without regularization. This observation is consistent with the idea that there is not an easy fix to the discrepancies revealed by metamers that simply involves adding an additional term to the optimization. And in some domains (such as audio) it is not obvious what to use for a regularizer. While the use of additional criteria to encourage the optimization to stay close to the manifold of “natural” examples is an interesting direction to explore further (and likely has useful applications), we emphasize that it is at odds with the goal of testing whether a model on its own replicates the properties of a biological sensory system.

#### Online behavioral experiments

All behavioral experiments presented in the main text were run on Amazon Mechanical Turk. To increase data quality, Amazon Turk qualifications were set to relatively stringent levels: the “HIT Approval Rate for all Requesters’ HITs” had to be at least 97% and the “Number of HITs Approved” had to exceed 1000. Example code to run the online experiments is available at https://github.com/jenellefeather/model_metamers_pytorch. All experiments with human participants (both online and in-lab) were approved by the Committee On the Use of Humans as Experimental Subjects at the Massachusetts Institute of Technology and were conducted with the informed consent of the participants.

#### Stimuli - image experiments

Each stimulus belonged to one of the 16 entry-level MS COCO (Microsoft Common Objects in Context) categories. We used a mapping from these 16 categories to the corresponding ImageNet1K categories (where multiple ImageNet1K categories can map onto a single MS COCO category), used in a previous publication (18). For each of the 16 categories, we selected 25 examples from the ImageNet1K validation dataset for a total of 400 natural images that were used to generate stimuli. A square center crop was taken for each ImageNet1K image (with the smallest dimension of the image determining the size) and the square image was rescaled to the necessary input dimensions for each ImageNet1K trained model. Metamers were generated for each of the 400 images to use for the behavioral experiments.

#### Stimuli - auditory experiments

A set of two-second speech audio excerpts with no background noise was used to generate stimuli. These clips were randomly chosen from the test set of the Word-Speaker-Noise dataset described above, constrained such that only the WSJ corpus was used. We further ensured that the clips were taken from unique sources within WSJ, and that the sounds were cropped to the middle two seconds of the training set clip such that the labeled word was centered at the one-second mark. To reduce ambiguity about the clip onset and offset (which were helpful in judging whether a word was centered on the clip), we also screened the chosen clips to ensure that the beginning 0.25s or end 0.25s of the clip was no more than -20dB quieter than the full clip. 400 clips were chosen subject to these constraints and such that each clip contained a different labeled word. Metamers were generated for each of the 400 clips to use for the behavioral experiments.

#### Image behavioral experiment

We created a visual experiment in JavaScript similar to that used in a previous publication (18). Participants were tasked with classifying an image into one of 16 presented categories (airplane, bear, bicycle, bird, boat, bottle, car, cat, chair, clock, dog, elephant, keyboard, knife, oven, truck). Each category had an associated image icon that participants chose from during the experiment. Each trial began with a fixation cross at the center of the screen for 300ms, followed by a natural image or a model metamer presented at the center of the screen for 300ms, followed by a pink noise mask presented for 300ms, followed by a 4×4 grid containing all 16 icons. Participants selected an image category by clicking on the corresponding icon. To minimize effects of internet disruptions, we ensured that the image was loaded into the browser cache before the trial began. We note that the precise timing of the image presentation for web experiments via JavaScript is less controlled compared to in-lab experiments, but any variation in timing should be similar across conditions and should average out over trials. To assess whether any timing variation in the online experiment set up might have affected overall performance, we compared recognition performance on natural images to that measured during in-lab pilot experiments (with the same task but different image examples) reported in an earlier conference paper (36). The average online performance across all natural images was on par or higher than that measured in-lab (in-lab proportion correct = 0.888 ± 0.0240) for all experiments.

The experimental session began with 16 practice trials to introduce participants to the task. There was one trial for each category, each presenting a natural image pulled from the ImageNet1K training set. Participants received feedback for these first 16 trials which included their answer and the correct response. Participants then began a 12-trial demo experiment which contained some natural images and some model metamers generated from the ImageNet1K training set. The goal of this demo experiment was two-fold – first, to introduce participants to the types of stimuli they would see in the main experiment and second, to be used as a screening criterion to remove participants who were distracted, misunderstood the task instructions, had browser incompatibilities, or were otherwise unable to complete the task. Participants were only allowed to start the main experiment if they correctly answered 7/12 correct on the demo experiment, which was the minimum that were correctly answered for these same demo stimuli by 16 in-lab participants in a pilot experiment (36). 341/417 participants passed the demo experiment and chose to move onto the main experiment.

There were 6 different main image experiments, each including a set of conditions (model stages) to be compared. The stimuli for each main experiment were generated from a set of 400 natural images. Participants only saw one natural image or metamer for each of the 400 images in the behavioral stimulus set. Participants additionally completed 16 catch trials. These catch trials each consisted of an image that exactly matched the icon for one of the classes. Participant data was only included in the analysis if the participant got 15/16 of these catch trials correct (270/341 participants were included). Of these participants, 125 self-identified as female, 143 as male, and 2 did not report; mean age= 42.1, minimum age=20, maximum age=78. For all but the HMAX experiment, participants completed 416 trials – one for each of the 400 original images, plus the 16 catch trials. The 400 images were randomly assigned to the experiment conditions subject to the constraint that each condition had approximately the same number of trials. Specifically, each condition was initially allocated ceiling(400/N) trials, and then trials were removed at random until the total number of trials was equal to 400. In addition, if the stimulus for a condition did not pass the metamer optimization criteria (and thus, had to be omitted from the experiment), the natural image was substituted for it as a placeholder, and analyzed as an additional trial for the natural condition. These two constraints resulted in the number of trials per condition varying somewhat across participants (see table below). The resulting 416 total trials were then presented in random order across the conditions of the experiment. The HMAX experiment used only 200 of the original 400 images, for a total of 216 trials. The experiment was run with a smaller number of stimuli than other experiments because it contained only 6 conditions (metamer optimization was run for all 400 stimuli and a subset of 200 images was randomly chosen from the subset of the 400 images for which metamer optimization was successful in every stage of the model). HMAX metamers were black and white, while all metamers from all other models were in color.

Model performance on this 16-way classification task was evaluated by measuring the predictions for the full 1000-way ImageNet classification task and finding the maximum probability for a label that was included in the 16-class dataset (231 classes).

**Table 3:**
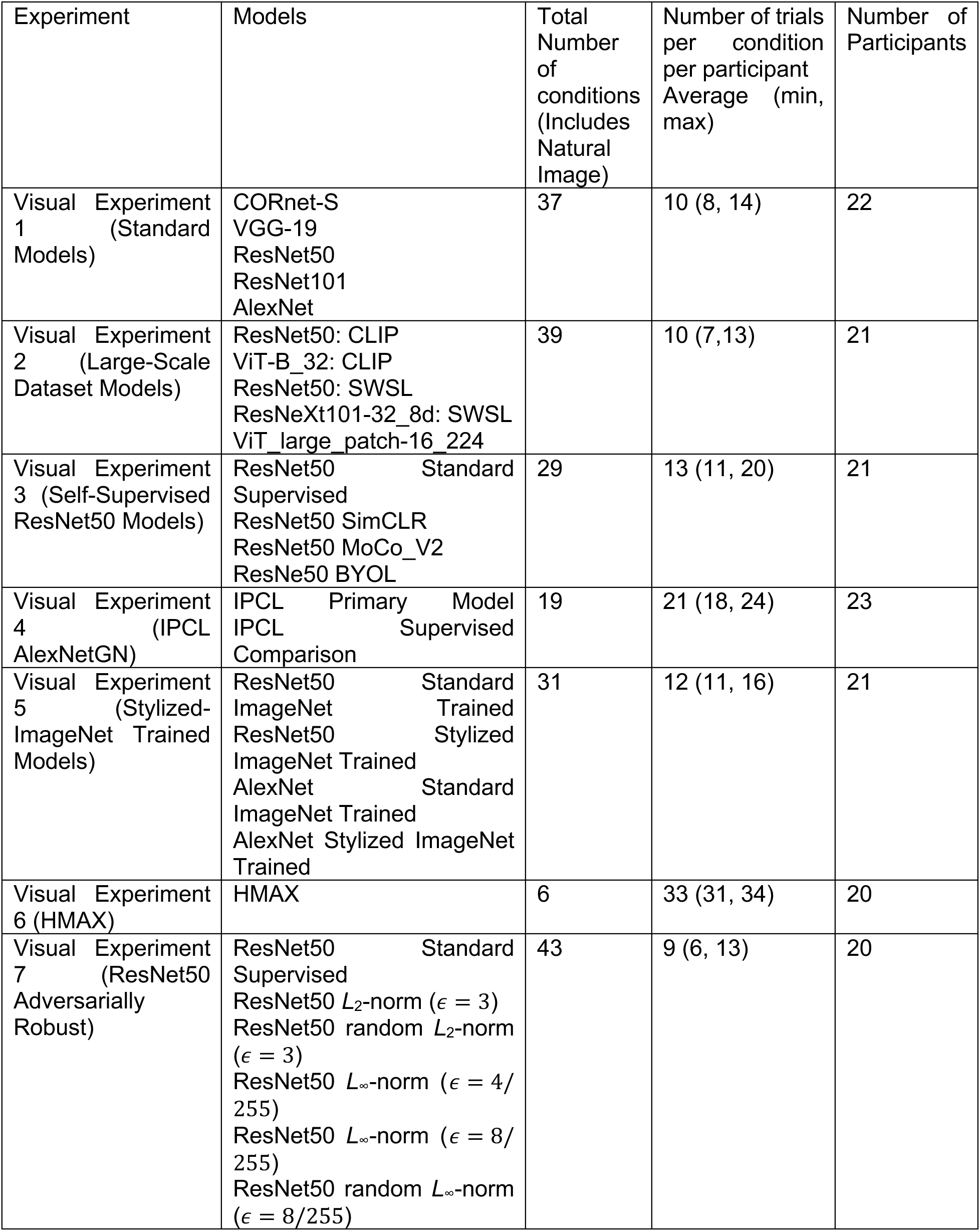
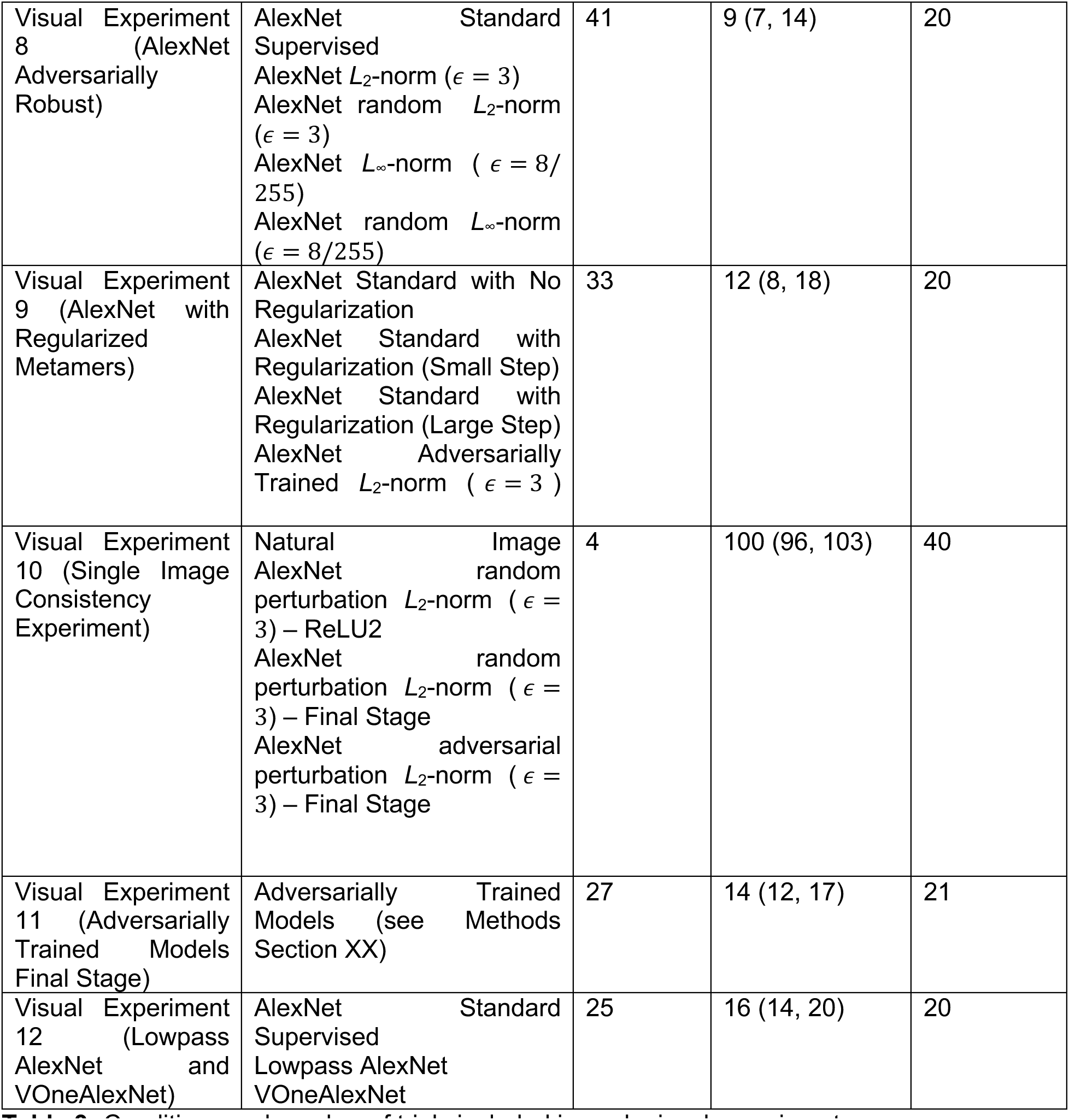
Conditions and number of trials included in each visual experiment.

#### Auditory behavioral experiment

We developed an auditory experiment in JavaScript that was similar to an experiment used in earlier publications from our lab (5, 36). Each human participant listened to a two-second audio clip and had to choose one of 793 word-labels corresponding to the word in the middle of the clip (centered at the one-second mark of the clip). Each trial began with the participant hearing the audio clip and typing the word they thought they heard into a response box. As participants typed, words matching the letter string they were typing appeared below the response box to help participants identify words that were in the response list. Once a word was typed that matched one of the 793 responses, participants could move onto the next trial.

To increase data quality, participants first completed a short experiment (six trials) that screened for the use of headphones (108). If participants scored 5/6 or higher on this screen (224/377 participants), they moved onto a practice experiment consisting of 10 natural audio trials with feedback (drawn from the training set), designed to introduce the task. This was followed by a demo experiment of 12 trials without feedback. These 12 trials contained both natural audio and model metamers, using the same set of 12 stimuli as in a demo experiment used for earlier in-lab experiments (36). As with the visual demo experiment, the goal of the audio demo experiment was to introduce participants to the type of stimuli they would hear in the main experiment and to screen out poorly performing participants. A screening criteria was set at 5/12, which was the minimum for 16 in-lab participants in earlier work (36). 154/224 participants passed the demo experiment and chose to move onto the main experiment. We have repeatedly found that online auditory psychophysical experiments qualitatively and quantitatively reproduce in-lab results, provided that steps such as these are taken to help ensure good audio presentation quality and attentive participants (109–112). The average online performance on natural stimuli was comparable to in-lab performance reported in (36) on natural stimuli using the same task with different audio clips (in-lab proportion correct = 0.863 ± 0.0340).

There were 6 different main auditory experiments, each including a set of conditions (multiple models and multiple model stages) to be compared. The design of these experiments paralleled the image experiments. The stimuli for each main experiment were generated from the set of 400 natural speech behavioral stimuli described above. Participants only heard one natural speech or metamer stimulus for each of the 400 excerpts in the behavioral stimulus set. Participants additionally completed 16 catch trials. These catch trials each consisted of a single word corresponding to one of the classes. Participant data was only included in the analysis if the participant got 15/16 of these trials correct (this inclusion criterion removed 8/154 participants). As the auditory experiment was long, some participants chose to leave the experiment early and their data was excluded from analysis (23/154). An additional 3 participants were excluded due to self-reported hearing loss, for a total of 120 participants across all auditory experiments. Of these participants, 45 self-identified as female, 68 as male, and 7 chose not to report; mean age=39.0, minimum age=22, maximum age=77. For all but the Spectemp experiment, participants completed 416 trials – one for each of the 400 original excerpts, plus the 16 catch trials. The 400 excerpts were randomly assigned to the experiment conditions subject to the constraint that each condition had approximately the same number of trials. As in the visual experiments, each condition was initially allocated ceiling(400/N) trials, and then trials were removed at random until the total number of trials was equal to 400. If the model stage selected for an image produced a metamer that did not pass the metamer optimization criteria (and thus, was omitted from experiment stimuli), the natural audio was used instead, but was not included in the analysis. As in the visual experiments, these two constraints resulted in the number of trials per condition varying somewhat across participants (see Table 4). The resulting 416 total trials were then presented in random order across the conditions of the experiment. The Spectemp experiment used only 200 of the original 400 excerpts, for a total of 216 trials. This experiment was run with a smaller number of stimuli because it contained only 6 conditions (metamer optimization was run for all 400 stimuli and a subset of 200 was randomly chosen from the subset of the 400 original excerpts for which metamer optimization was successful in every stage of the model). We collected online data in batches until we reached the target number of participants for each experiment.

**Table 4:**
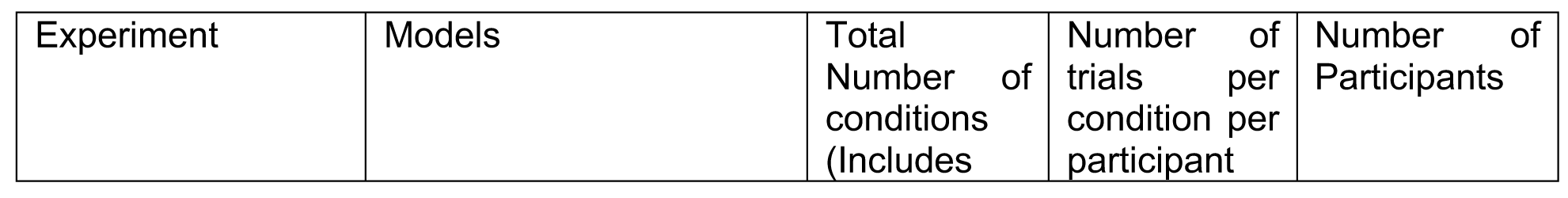
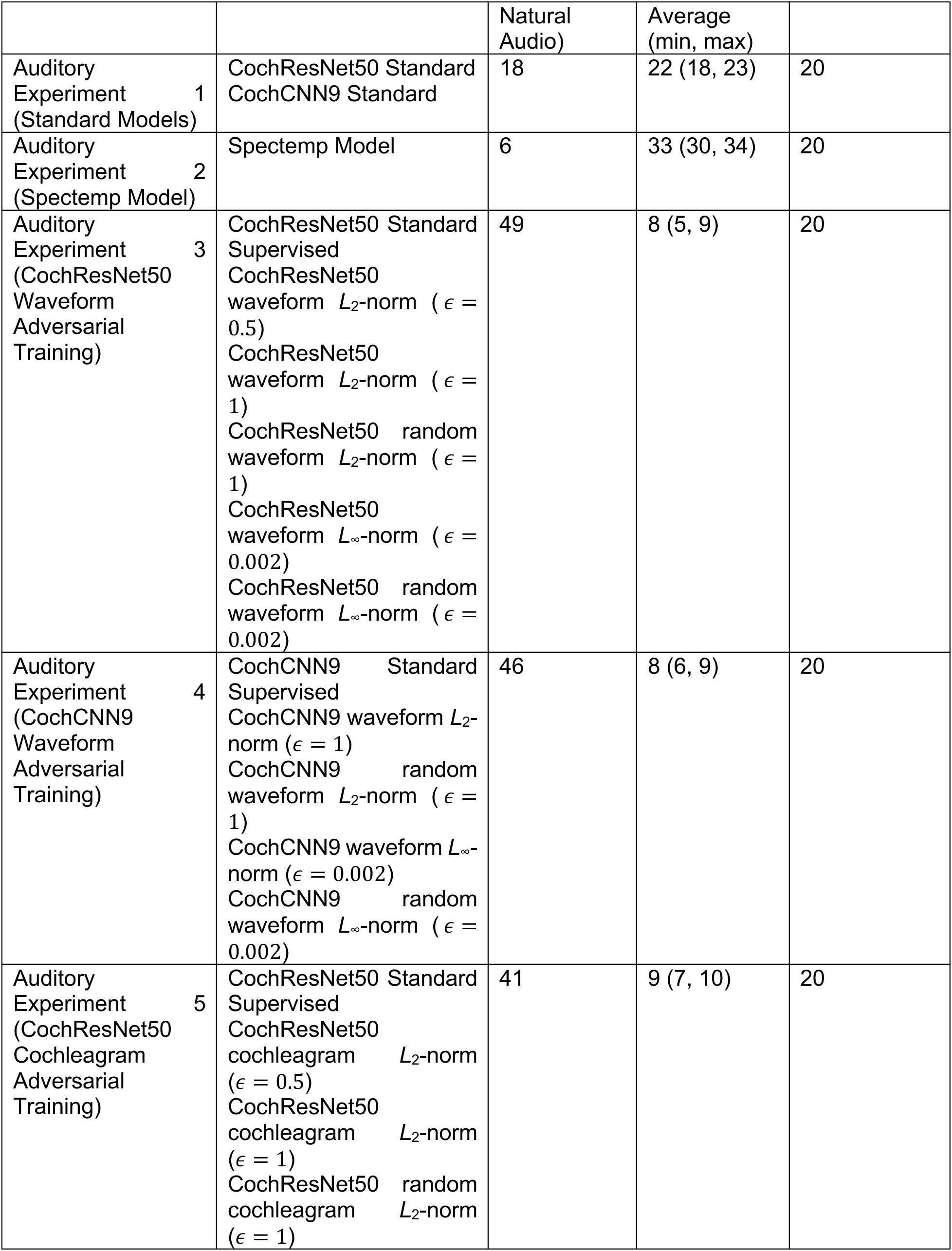
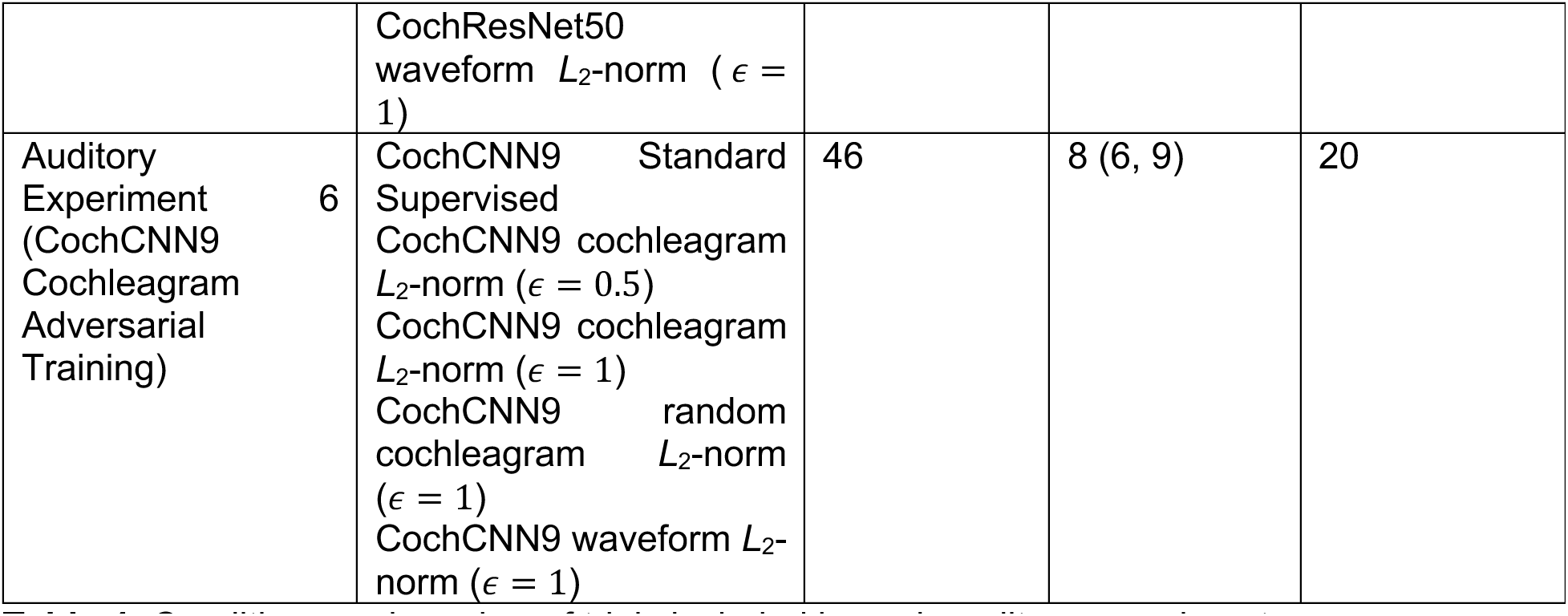
Conditions and number of trials included in each auditory experiment.

#### Statistical tests – difference between human and model recognition accuracy

All human recognition experiments involving neural network models were analyzed by comparing human recognition of a generating model’s metamers to the generating model’s recognition of the same stimuli (its own metamers). Each human participant was run on a distinct set of model metamers; we presented each set to the generation model and measured its recognition performance for that set. Thus, if N human participants performed an experiment, we obtained N model recognition curves. We ran mixed model repeated measures ANOVAs with a within-group factor of metamer generation model stage and a between-group factor of observer (human or model observer), testing for both a main effect of observer and an interaction between observer and model stage. Data were non-normal due to a prevalence of values close to 1 or 0 depending on the condition, and so we evaluated statistical significance non-parametrically, using permutation tests in which we compared the observed F-statistic to that obtained after randomly permuting the data labels. To test for main effects, we permuted observer labels (model vs. human). To test for interactions of observer and model stage, we randomly permuted both the observer labels and the model stage labels, independently for each participant. In each case we used 10,000 random permutations and computed a p value by comparing the observed F-statistic to the null distribution of F-statistics from permuted data (i.e., the p value was one minus the rank of the observed F-statistic divided by the number of permutations).

Because the classical models did not themselves perform recognition judgments, rather than comparing human and model recognition as in the experiments involving neural network models, we instead tested for a main effect of model stage on human observer recognition. We performed a single-factor repeated measure ANOVA using a within-group factor of model stage, again evaluating statistical significance non-parametrically. We randomly permuted the model stage labels of the recognition accuracy data, independently for each participant (with 10,000 random permutations).

#### Statistical tests – difference between human recognition of metamers generated from different models

To compare human recognition of metamers generated from different models, we ran a repeated measures ANOVA with within-group factors of model stage and generating model. This type of comparison was only performed in cases where the generating models had the same architecture (so that the model stages were shared between models). We again evaluated statistical significance non-parametrically, by comparing the observed F-statistic to a null distribution of F-statistics from permuted data (10,000 random permutations). To test for a main effect of generating model, we randomly permuting the generating model label, independently for each participant. To test for an interaction between the generating model and model stage, we permuted both the generating model label and the model stage label, independently for each participant.

#### Power analysis to determine sample sizes

We planned to run ANOVAs examining the interaction between human performance and model performance, the interaction between model and stage, the interaction between standard models and adversarially trained models, the interaction between different types of adversarial training, and the interaction between standard supervised models and self-supervised models. To estimate the number of participants necessary to be well powered for these analyses, we ran a pilot experiment comparing the standard versus adversarially trained ResNet50 and CochResNet50 models, as this experiment included the largest number of conditions and we expected that the differences between different types of adversarially trained models would be subtle, and would put an upper bound on the sample sizes that would be needed across experiments.

For the vision experiment, we ran 10 participants in a pilot experiment on Amazon Mechanical Turk. The format of the pilot experiment was identical to that of the main experiments in this paper, with the exception that we used a screening criteria of 8/12 correct for the pilot, rather than the 7/12 correct used for the main experiment. In this pilot experiment, the smallest effect size out of those we anticipated analyzing in the main experiments was the comparison between the *L*∞-norm (є = 8/256) adversarially trained ResNet50 and the *L*2-norm (є = 3) adversarially trained Resnet50, with a partial eta squared value of 0.10 for the interaction. A power analysis was performed with g*power (113); 18 participants were needed to have a 95% chance of seeing an effect of this size at a p<.01 significance level. We thus set a target of 20 participants for each main vision experiment.

For the auditory experiments we ran 14 participants in a pilot experiment on Amazon Mechanical Turk. The format of the pilot experiment was identical to that of the main experiments in this paper with the exception that 8 of the 14 participants only received 6 original audio trials with feedback, while in the main experiment 10 trials with feedback were used. As in the vision pilot experiment, in this pilot the smallest effect size of interest was that for the comparison between the *L*∞-norm (є = 0.002) adversarially trained CochResNet50 and the *L*2-norm (є = 1) waveform adversarially trained CochResNet50, yielding a partial eta squared value of 0.37 for the interaction. A power analysis was performed with g*power; this indicated that 12 participants were needed to have a 95% change of seeing an effect for this size at a p<0.01 significance level. To match the image experiments, we set a target of 20 participants for each main auditory experiment.

#### Split-Half Reliability Analysis of Metamer Confusion Matrices (***Supplementary Figure 2***)

To assess whether human participants had consistent error patterns, particularly in metamer conditions with poor recognizability, we compared confusion matrices from split halves of participants. Each row of the confusion matrix (corresponding to a category label) was normalized by the number of trials for that label. We then computed the Spearman correlation between the confusion matrices from each split and compared this correlation to that obtained from confusion matrices from permuted participant responses for the condition. We computed the correlation for 1000 random splits of participants (splitting the participants in half) and using a different permutation of the response for each split. We counted the number of times that the difference between the true split-half correlation and the shuffled correlation was less than or equal to 0 (η_overlap_), and the p-value was computed as:

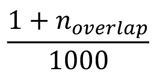

#### Experiment testing effect of optimization algorithm (Supplementary Figure 3)

An in-lab visual experiment was conducted to analyze whether differences in the optimization strategy would have an effect on the human-recognizability of model metamers. This experiment used one example visual model (a ResNet50 architecture). The optimization code and techniques used for this small in-lab experiment differed from any of the experiments described in the main text of this paper (they followed the methods used in our previous work (36)) but showed a similar main effect of model stage for standard neural network models, where model metamers generated from late stages of ImageNet1K task-optimized ResNet50 models were unrecognizable to humans.

#### Visual models included in the experiment

The models and metamer generation for this experiment were implemented in TensorFlow v1.12 (114). For these experiments, we converted the ResNet50 model available via the PyTorch Model Zoo to TensorFlow using ONNX (version 1.6.0).

The experiment was run together with an unrelated pilot experiment comparing four other models, the data for which are not analyzed here. To reduce the number of conditions, the stage corresponding to ResNet50 “layer1” was not included in metamer generation or in the experiment (from ad-hoc visual inspection of a few examples, the model metamers for conv1_relu1, layer1, and layer2 looked similar enough to the natural image that we expected ceiling performance for all conditions).

#### Model metamer optimization

Metamers were optimized using TensorFlow code based on that used for a previous conference paper (36). Unless otherwise noted the methods were identical to those used in the main experiments of this paper.

We tested two different optimization schemes for generating model metamers. The first used stochastic gradient descent with 15000 iterations, where each step of gradient descent was constrained to have an *L2* norm of 1. The second method used the Adam optimizer (115), which uses an adaptive estimation of first and second order moments of the gradient. Metamers were generated with 15000 iterations of the Adam optimizer with an exponentially decaying learning rate (initial learning rate of 0.001, 1000 decay steps, and a decay rate of 0.95).

#### Stimuli – Image dataset for optimization strategy experiment

Similar to the other visual experiments in this paper, each stimulus belonged to one of the 16 entry-level MS COCO categories. For each of the 16 categories, we randomly selected 16 examples from the ImageNet1K training dataset using the list of images provided by (18) for a total of 256 natural images that were used to generate stimuli. Metamers were generated for each of the 256 images to use for the behavioral experiments.

#### In-lab visual experiment

The experimental setup was the same as the experiment described for our main experiments (but implemented in MATLAB Psychotoolbox rather than JavaScript), in which participants had to classify an image into one of the 16 presented categories. Each trial began with a fixation cross at the center of the screen for 300ms, followed by a natural image or a model metamer presented at the center of the screen for 200ms, followed by a pink noise mask presented for 200ms, followed by a 4×4 grid containing all 16 icons. Stimuli were presented on a 20” ACER LCD (backlit LED) monitor with a spatial resolution of 1600×900 and a refresh rate of 60Hz. Stimuli spanned 256×256 pixels and were viewed at a distance of approximately 62 cm (set by the chair position, which was fixed; participants were free to position themselves in the chair as was most comfortable, which introduced minor variation in viewing distance).

Before the experiment, each participant was shown a printout of the 16 category images with labels and the experimenter pointed to and read each category. This was followed by a demo experiment with 12 trials without feedback (same stimuli as in the main experiments, but performance was not used to exclude participants).

Each participant saw 6 examples from each condition, chosen such that each natural image or metamer was from a unique image from the 256-image behavioral set. Ten participants completed the experiment.

#### Human consistency of errors for individual stimuli (Supplementary Figure 7)

In the experiment to evaluate the consistency of errors for individual stimuli (Supplementary Figure 7), we only included four conditions in order to collect enough data to analyze performance on individual images: natural images, metamers from the ReLU2 and final stages for the random perturbation trained AlexNet *L*2-norm (є = 1) model, and metamers from the final stage of the adversarial perturbation trained AlexNet *L*2-norm (є = 1) model. The rationale for the inclusion of these stages was that the ReLU2 stage of the Random perturbation AlexNet and the Final stage of the Adversarial perturbation AlexNet had similarly recognizable metamers (Figure 4c), whereas metamers from the Final stage of the Random perturbation AlexNet were recognized no better than chance by humans.

To first assess the reliability of the recognizability of individual stimuli (Supplementary Figure 7a), we measured the Spearman correlation of the recognizability (proportion correct) of each stimulus across splits of participants, separately for each of the four conditions. We averaged this correlation over 1000 random splits of participants. P-values were computed non-parametrically by shuffling the participant responses for each condition and each random split, and computing the number of times the true average Spearman ρ was lower than the shuffled correlation value. We only included images in the analysis that had at least 4 trials in each split of participants, and when there were more than 4 trials in a split, we only included four of the trials, randomly selected, in the average, to avoid having some images exert more influence on the result than others.

#### Most and least recognizable images

To analyze the consistency of the “most” and “least” recognizable metamers in each condition (Supplementary Figure 7b), we used one split of participants to select 50 images that had highest recognition score and 50 images with lowest recognition score. We then measured the recognizability of these images in the second split of participants, and assessed whether the “most” recognizable images had a higher recognition score than the “least” recognizable images. P-values for this comparison were computed by using 1000 splits of participants and measuring the proportion of splits in which the difference between the two scores was greater than 0.

To select examples of the most and least recognizable images (Supplementary Figure 7c&d) we only included example stimuli with at least 8 responses for both the natural image condition and the model metamer stage under consideration, and which had 100% correct responses on the natural image condition. From this set we selected the “most” recognizable images (as those with scores of 100% correct for the considered condition) and the “least” recognizable images (as those with scores of 0% correct).

#### Model-Brain Comparison Metrics for Visual Models

We utilized the Brain-Score (42) platform to obtain metrics of neural similarity in four visual cortical areas of the macaque monkey brain: V1, V2, V4, and IT. For each model considered, we analyzed only the stages that were included in our human metamer recognition experiments. We note that some models may have had higher brain similarity scores had we analyzed all stages. Each of these model stages was fit to a public data split for each visual region, with the best-fitting stage for that region selected for further evaluation. The match of this model stage to brain data was then evaluated on a separate set of evaluation data for that region. Evaluation data for V1 consisted of the average of 23 benchmarks: 22 distribution-based comparison benchmarks from (116) and the V1 partial-least-squares regression (PLS) benchmark from (117). Evaluation data for V2 consisted of the V2 PLS benchmark from (117). Evaluation data for V4 consisted of the average of four benchmarks: the PLS V4 benchmark from (118), the PLS V4 benchmark from (119), the PLS V4 benchmark from (120), and the PLS V4 benchmark from (121). Evaluation data for IT consisted of the average of 4 benchmarks: the PLS IT benchmark from (118), the PLS IT benchmark from (119), the PLS IT benchmark from (120), and the PLS IT benchmark from (121). When comparing the metamer recognizability to the Brain-Score results, we used the human recognition of metamers from the model stage selected as the best match for each visual region.

We used Spearman correlations to compare metamer recognizability to the Brain-Score results. We note that the analogous Pearson correlations were lower, and that none reached statistical significance. We report Spearman correlations on the grounds that the recognizability was bounded by 0 and 1, and to be conservative with respect to our conclusion that metamer recognizability is not explained by standard model-brain comparison metrics.

We estimated the noise ceiling of the correlation between Brain-Score results and human recognizability of model metamers as the geometric mean of the reliabilities of each quantity. To estimate the reliability of the metamer recognizability, we split the participants for an experiment in half and measured the recognizability of metamers for each model stage used to obtain the Brain-Score results (i.e., the best-predicting stage for each model for the brain region under consideration). We then calculated the Spearman correlation between the recognizability for the two splits, and Spearman-Brown corrected to account for the 50% reduction in sample size from the split. This procedure was repeated for 1000 random splits of participants. We then took the mean across the 1000 splits as an estimate of the reliability. This estimated reliability was 0.917 for V1, 0.956 for V2, 0.924 for V4, and 0.97 for IT. As we did not have access to splits of the neural data used for Brain-Score, we estimated the reliability of the Brain-Score results as the Pearson correlation of the score for two sets of neural responses to the same images, Spearman-Brown corrected, reported in (73). This estimated reliability was only available for IT (r=0.87), but we assume that the reliability would be comparable for other visual areas.

#### Model-Brain Comparison Metrics for Auditory Models

The auditory fMRI analysis closely followed that of a previous publication (5) using the fMRI dataset collected in another previous publication (74).The key components of the dataset and analysis methods are replicated here, but for additional details see (5, 74). The text from sections “*fMRI data acquisition and preprocessing*” and “*Voxel and ROI selection”* is replicated from a previous publication with minor edits (5).

#### Natural sound stimuli

The stimulus set was composed of 165 two-second natural sounds spanning eleven categories (instrumental music, music with vocals, English speech, foreign speech, non-speed vocal sounds, animal vocalization, human-non-vocal sound, animal non-vocal sound, nature sound, mechanical sound, or environment sound). The sounds were presented in a block design with five presentations of each two-second sound. To prevent sounds from being played at the same time as scanner noise, a single fMRI volume was collected following each sound presentation (“sparse scanning”). This resulted in a 17-second block. Blocks were grouped into 11 runs with 15 stimuli each and four blocks of silence. Silence blocks were the same duration as the stimulus blocks and were used to estimate the baseline response. Participants performed a sound-intensity discrimination task to increase attention. One sound in the block of five was presented 7dB lower than the other four (the quieter sound was never the first sound) and participants were instructed to press a button when they heard the quieter sound. Sounds were presented with MR-compatible earphones (Sensimetrics S14) at 75 dB SPL for the louder sounds and 68dB SPL for the quieter sounds.

#### fMRI data acquisition and preprocessing

MR data was collected on a 3T Siemens Trio scanner with a 32-channel head coil at the Athinoula Martinos Imaging Center of the McGovern Institute for Brain Research at MIT. Data was first published in (74), and was re-analyzed for this paper. Each functional volume consisted of fifteen slices oriented parallel to the superior temporal plane, covering the portion of the temporal lobe superior to and including the superior temporal sulcus. Repetition time (TR) was 3.4 s (acquisition time was only 1 s due to sparse scanning), echo time (TE) was 30 ms, and flip angle was 90 degrees. For each run, the five initial volumes were discarded to allow homogenization of the magnetic field. In-plane resolution was 2.1 x 2.1 mm (96 x 96 matrix), and slice thickness was 4 mm with a 10% gap, yielding a voxel size of 2.1 x 2.1 x 4.4 mm. iPAT was used to minimize acquisition time. T1-weighted anatomical images were collected in each subject (1mm isotropic voxels) for alignment and surface reconstruction.

Functional volumes were preprocessed using FSL and in-house MATLAB scripts. Volumes were corrected for motion and slice time. Volumes were skull-stripped, and voxel time courses were linearly detrended. Each run was aligned to the anatomical volume using FLIRT and BBRegister. These preprocessed functional volumes were then resampled to vertices on the reconstructed cortical surface computed via FreeSurfer, and were smoothed on the surface with a 3mm FWHM 2D Gaussian kernel to improve SNR. All analyses were done in this surface space, but for ease of discussion we refer to vertices as ‘‘voxels’’ in this paper. For each of the three scan sessions, we estimated the mean response of each voxel (in the surface space) to each stimulus block by averaging the response of the second through the fifth acquisitions after the onset of each block (the first acquisition was excluded to account for the hemodynamic lag). Pilot analyses showed similar response estimates from a more traditional GLM (74). These signal-averaged responses were converted to percent signal change (PSC) by subtracting and dividing by each voxel’s response to the blocks of silence. These PSC values were then downsampled from the surface space to a 2mm isotropic grid on the FreeSurfer-flattened cortical sheet.

#### Voxel and ROI selection

We used the same voxel selection criterion as Norman-Haignere et al. (2015), selecting voxels with a consistent response to sounds from a large anatomical constraint region encompassing the superior temporal and posterior parietal cortex. Specifically, we used two criteria: (1) a significant response to sounds compared with silence (p < 0.001); and (2) a reliable response to the pattern of 165 sounds across scans. The reliability measure was as follows:

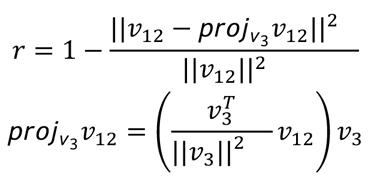

where *ν_12_* is the response of a single voxel to the 165 sounds averaged across the first two scans (a vector), and *ν_3_* is that same voxel’s response measured in the third. The numerator in the second term in the first equation is the magnitude of the residual left in *ν_12_* after projecting out the response shared with *ν_3_*. This ‘‘residual magnitude’’ is divided by its maximum possible value (the magnitude of *ν_12_*). The measure is bounded between 0 and 1, but differs from a correlation in assigning high values to voxels with a consistent response to the sound set, even if the response does not vary substantially across sounds. We found that using a more traditional correlation-based reliability measure excluded many voxels in primary auditory cortex because some of them exhibit only modest response variation across natural sounds. We included voxels with a value of this modified reliability measure of 0.3 or higher, which when combined with the sound responsive t-test yielded a total of 7694 voxels across the eight participants (mean number of voxels per subject: 961.75; range: 637-1221).

We localized four regions of interest (ROIs) in each participant, consisting of voxels selective for (1) frequency (i.e., tonotopy), (2) pitch, (3) speech, and (4) music. In each case we ran a ‘‘localizer’’ statistical test and selected the top 5% most significant individual voxels in each subject and hemisphere (including all voxels identified by the sound-responsive and reliability criteria described above). We excluded voxels that were identified in this way by more than one localizer. The frequency, pitch, and speech localizers required acquiring additional imaging data, and were collected either during extra time during the natural sound stimuli scan sessions or on additional sessions on different days. Scanning acquisition parameters were identical to those used to acquire the natural sounds data. Throughout this paper we refer to voxels chosen by these criteria as ‘‘selective,’’ for ease and consistency. There were a total of 379 voxels in the Tonotopic ROI, 379 voxels in the pitch ROI, 393 voxels in the music ROI and 379 voxels in the speech ROI.

To identify frequency-selective voxels, we measured responses to pure tones in six different frequency ranges (center frequencies: 200, 400, 800, 1600, 3200, 6400 Hz) (122, 123). For each voxel, we ran a one-way ANOVA on its response to each of these six frequency ranges and selected voxels that were significantly modulated by pure tones (top 5% of all selected voxels in each subject, ranked by p values). Although there was no spatial contiguity constraint built into our selection method, in practice most selected voxels were contiguous and centered around Heschl’s gyrus.

To identify pitch-selective voxels, we measured responses to harmonic tones and spectrally-matched noise (123). For each voxel we ran a one-tailed t-test evaluating whether the response to tones was greater than that to noise. We selected the top 5% of individual voxels in each subject that had the lowest p values for this contrast.

To identify speech-selective voxels, we measured responses to German speech and to temporally scrambled (‘‘quilted’’) speech stimuli generated from the same German source recordings (124). We used foreign speech to identify responses to speech acoustical structure, independent of linguistic structure. Note that two of the participants had studied German in school and for one of these participants we used Russian utterances instead of German. The other subject was tested with German because the Russian stimuli were not available at the time of the scan. For each voxel we ran a one-tailed t-test evaluating whether responses were higher to intact speech than to statistically matched quilts. We selected the top 5% of all selected voxels in each subject.

To identify music-selective voxels, we used the music component derived by (74). We inferred the ‘‘voxel weights’’ for each voxel to all six of the components from its response to the 165 sounds:

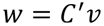

where c contains the inferred voxel weights (a vector of length 6), d′ is the Moore-Penrose pseudoinverse of the ‘‘response components’’ (a 6 by 165 matrix), and *ν* is the measured response of a given voxel (vector of length 165). We assessed the significance of each voxel’s music component weight via a permutation test. During each iteration, we shuffled all the component elements, recomputed this new matrix’s pseudoinverse, and recomputed each voxel’s weights via the matrix multiply above. We performed this procedure 10,000 times, and fit a Gaussian to each voxel’s null distribution of music weights. We then calculated the likelihood of the empirically observed voxel weight from this null distribution, and took the top 5% of voxels with the lowest likelihood under this null distribution.

#### Voxelwise encoding analysis

Each of the 165 sounds from the fMRI experiment was resampled to 20,000Hz (the sampling rate of the training dataset) and passed through each auditory model. We measured the model response at each model stage for each sound. To compare the model responses to the fMRI response we averaged over the time dimension for all units that had a temporal dimension (all model stages except fully connected layers).

In the CochCNN9 architecture, this resulted in the following number of regressors for each stage: cochleagram (211), relu0 (6818), relu1 (4608), relu2 (4608), relu3 (9216), relu4 (4608), avgpool (2560), relufc (4096), final (794). In the CochResNet50 architecture, this resulted in the following number of regressors for each stage: cochleagram (211), conv1_relu1 (6784), layer1 (13568), layer2 (13824), layer3 (14336), layer4 (14336), avgpool (2048), final (794).

We used the features sets extracted from model stages to predict the fMRI response to the natural sound stimuli. Each voxel’s time-averaged responses were modeled as a linear combination of a stage’s time-averaged unit responses. Ten random train-test splits (83/82) were taken from the 165 sound dataset. For each split we estimated a linear mapping using L2-regularized (“ridge”) linear regression using RidgeCV from the scikit learn library version 0.23.1 (125). Ridge regression places a zero-mean gaussian prior on the regression coefficients, corresponding to solving the regression problem given by:

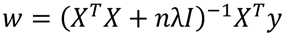

whereis the number of stimuli used for estimation (here equal to 83), c is the X-length column vector the length of the regressors (number of extracted features for the model stage), *y* is an *n*-length column vector containing the voxels response to each sound (length *n*), *X* is a matrix containing the regressors (*n* stimuli by *d* regressors, the extracted features for each sound), g is the identity matrix, and λ is the ridge regression regularization parameter. The mean of each column was subtracted from the regressor matrix before fitting.

The best ridge regression parameter for each voxel was independently selected using leave-one-out cross validation across the 83 training sounds in each split sweeping over 81 logarithmically spaced values (each power of 10 between 10^-40^ – 10^40^). For each held-out sound in the training set, the mean squared error of the prediction was computed using regression weights from the other 82 training set sounds, for each of the regularization coefficients. The regularization parameter that minimized the mean squared error on the held-out sounds was used to fit a linear mapping between model responses to all 83 training set sounds and the voxel responses. This mapping was used to predict the voxel response to the held-out 82 test sounds and fitting fidelity was evaluated with the squared Pearson correlation (r^2^) as a metric of explained variance of the predicted voxel response and the observed voxel response.

This measured explained variance was corrected for the effects of measurement noise by computing the reliability of the voxel responses and of the predicted voxel response. We use the correction for attenuation (126), which estimates the correlation between two variables independent of measurement noise, resulting in an unbiased estimator of the correlation coefficient that would be observed from noiseless data. The corrected variance explained is:

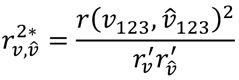

where *ν_123_* is the voxel response to the 82 test sounds averaged across all three scans, *ν_123_* is the predicted response to the 82 test sounds using the 83 training sounds to learn the regression weights, [is a function that computes the Pearson correlation coefficient, [*r^’^_12_* is the correction for reliability of the voxel responses, and [*r_ν_^’^* is the correction for reliability of the predicted voxel response. [*r_ν_^’^* is computed as the median Spearman-Brown corrected Pearson correlation

between the 3 pairs of scans (scan 0 with scan 1, scan 1 with scan 2, scan 0 with scan 2), where the Spearman-Brown correction accounts for increased reliability expected from tripling the amount of data (127). [*r_ν_^’^* is similarly computed by using the training data each of the three pairs of scans to predict the test data from the same scan, and calculating the median Spearman-Brown corrected correlation for the predicted voxel responses. If voxels and/or predictions are very unreliable, this can lead to large corrected variance explained measures (128). We set a minimum value of 0.182 for [*r_ν_^’^* (corresponding to the value at which the correlation of two 83-dimensional random variables reaches significance at a threshold of p<.05; 83 being the number of training data values) and a minimum value of 0.183 for [*r_ν_^’^* (the analogous value for 82-dimensional random variables, corresponding to the number test data values).

The above computation of corrected variance explained was computed for each voxel using each stage response for each of 10 train/test splits of data. We took the median variance explained across the 10 splits of data. To provide one summary metric across each of the ROIs (all auditory voxels, tonotopic voxels, pitch voxels, music voxels, speech voxels) we chose the best stage for each ROI using a hold-one-participant out analysis. (Figure 7c) A summary measure for each participant and model stage was computed by taking the median across all voxels of the voxel-wise corrected variance explained values within the ROI. Holding out one participant, we averaged across the remaining participant values to find the stage with the highest variance explained within the given ROI. We measured the corrected variance explained for this stage in the held-out participant, recorded the stage that was used as the “best” model stage, and repeated the procedure holding out one participant at a time. This cross validation avoids issues of non-independence when selecting the best stage. We report the mean corrected variance explained across the participants. Metamer recognition was measured from the model stage most frequently chosen as the best predicting model stage across participants (in practice, nearly all participants had the same “best” model stage). When plotting the variance explained for each model stage (Figure 7d) we took the median variance explained across the 10 splits of data, and then took the median across all participants.

#### Statistical tests – fMRI predictions

One-sided paired sample t-tests were performed to test for a difference in the variance explained summary metrics between different models. Significance values were computed by constructing a null-distribution of t-statistics by randomly permuting the model assignment independently for each participant (10,000 permutations). The p-value is reported as one minus the rank of the observed t-statistic divided by the number of permutations. In cases where the maximum possible number of unique permutations was less than 10,000, we instead divided the rank by the maximum number of unique permutations

#### Noise ceiling estimates for correlation between metamer recognizability and fMRI metrics

We estimated the noise ceiling of the correlation between auditory fMRI predictivity and human recognizability of model metamers as the geometric mean of the reliabilities of each quantity. To estimate the reliability of the metamer recognizability, we split the participants for an experiment in half and measured the recognizability of metamers for the model stage that was most frequently chosen (across all participants) as the best-predicting stage for the ROI under consideration (i.e., the stages used for Figure 7c). We then calculated the Spearman correlation between the recognizability for the two splits, and Spearman-Brown corrected to account for the 50% reduction in sample size from the split. This procedure was repeated for 1000 random splits of participants. We then took the mean across the 1000 splits as an estimate of the reliability. This estimated reliability of the metamer recognizability was 0.811 for the best predicting stage of all auditory voxels, 0.829 for the best predicting stage of the tonotopic ROI, 0.819 for the best predicting stage of the pitch ROI, 0.818 for the best predicting stage of the music ROI, and 0.801 for the best predicting stage of the speech ROI. To estimate the reliability of the fMRI prediction metric, we took two splits of the fMRI participants and calculated the mean variance explained for each model using the stage for which recognizability was measured. We then computed the Spearman correlation between the explained variance for the two splits and Spearman-Brown corrected the result. We then repeated this procedure for 1000 random splits of the participants in the fMRI study, and took the mean across the 1000 splits as the estimated reliability. This reliability of fMRI predictions was 0.923 for all auditory voxels, 0.768 for the tonotopic ROI, 0.922 for the pitch ROI, 0.796 for the music ROI, and 0.756 for the speech ROI.

#### Representational similarity analysis (Supplementary Figure 10)

To construct the model representational dissimilarity matrix (RDM) for a model stage, we computed the dissimilarity (1 minus the Pearson correlation coefficient) between the model activations evoked by each pair of the 165 sounds for which we had fMRI responses. Similarly, to construct the fMRI RDM, we computed the dissimilarity in voxel responses (1 minus the Pearson correlation coefficient) between all ROI voxel responses from a participant to each pair of sounds. Before computing the RDMs from the fMRI or model responses, we z-scored the voxel or unit responses. As a measure of fMRI and model similarity, we computed the Spearman correlation coefficient between the fMRI RDM and the model RDM.

To compute the RDM similarity for the model stage whose RDM best matched that of an ROI (Supplementary Figure 10a), we first generated 10 randomly selected train/test splits of the 165 sound stimuli into 83 training sounds and 82 test sounds. For each split, we computed the RDMs for each model stage and for each participant’s fMRI data for the 83 training sounds. We then chose the model stage that yielded the highest Spearman ρ measured between the model stage RDM and the participant’s fMRI RDM. Using this model stage, we measured the model and fMRI RDMs from the test sounds and computed the Spearman ρ. We repeated this procedure ten times, once each for 10 random train/test splits, and took the median Spearman ρ across the ten splits. We performed this procedure for all participants and computed the mean Spearman ρ across participants for each model. When comparing the RDM similarity from the best matching stage to metamer recognizability, we measured metamer recognizability from the model stage that was most frequently chosen as the best matching model stage across participants.

The representational similarity analysis is limited by measurement noise in the fMRI data. As an estimate of the upper bound for the RDM correlation that could be reasonably expected to be achieved between a model RDM and a single participant’s fMRI RDM, we calculated the correlation between one participant’s RDM and the average of all the other participant’s RDM. We held out one participant and averaged the RDMs across the remaining participants. The RDMs were measured from the same 10 train/test splits of the 165 sounds described in the previous paragraph, using the 82 test sounds for each split. We then calculated the Spearman ρ between the RDM from the held-out participant and the average participant RDM. We took the median Spearman ρ across the 10 splits of data to yield a single value for each participant. This procedure was repeated holding out each participant, and the upper bound shown in Supplementary Figure 10 is the mean across the measured value for each held out participant. This corresponds to the “lower bound” of the noise ceiling used in prior work (13). We plotted this upper bound on the results graphs rather than noise-correcting the human-model RDM correlation to be consistent with prior modeling papers that have used this analysis (13).

#### Model recognition of metamers generated from other models

To measure the recognition of a model’s metamers by other models, we took the generated image or audio that was used for the human behavioral experiments, provided it as input to a “recognition” model, and measured the 16 way image classification (for ImageNet1K models) or the 763-way word classification (for the auditory models).

The plots in Figure 8b show the average recognition by other models of metamers generated from a particular type of ResNet50 model. We used one “Standard” generation model (the standard supervised ResNet50). This curve is the average across all other vision recognition models (as shown in Figure 9a). The curve for self-supervised models shows results averaged across the three self-supervised generation models (SimCLR, MoCo_V2, and BYOL); the curve for adversarially trained models shows results averaged across the three adversarially trained ResNet50 models (trained with *L*2-norm (є = 3), *L*∞-norm (є = 4/255), and *L*∞-norm (є = 8/255) perturbations, respectively). For these latter two curves, we first computed the average curve for each recognition model across all three generation models, omitting the recognition model from the average if it was the same as the generation model (in practice, this meant that there was one fewer value included in the average for the recognition models that are part of the generation model group). We then averaged across the curves for each recognition model. The error bars on the resulting curves are the SEM computed across the recognition models.

Figure 8c was generated in an analogous fashion. We used one “Standard” generation model (the standard supervised CochResNet50). The curve for the waveform adversarially trained models shows results averaged across the three such CochResNet50 models (trained with *L*2- norm (є = 0.5), *L*2-norm (є = 1), and *L*∞-norm (є = 0.002) perturbations, respectively). The curve for the cochleagram adversarially trained models shows results averaged across the two such CochResNet50 models (trained with *L*2-norm (є = 0.5), and *L*2-norm (є = 1) perturbations, respectively). The group averages and error bars were computed as in Figure 8b.

## Data availability statement

Human data (auditory and visual human recognition performance and summarized fMRI data), and trained model checkpoints are available at https://github.com/jenellefeather/model_metamers_pytorch. The same repository also includes an interface to view/listen to the generated metamers used in the human recognition experiments. The Word-Speaker-Noise training dataset is available from the authors upon request.

## Code availability statement

Code for generating metamers, training models, and running online experiments is available at https://github.com/jenellefeather/model_metamers_pytorch. Auditory front-end (cochleagram generation) code is available at https://github.com/jenellefeather/chcochleagram.

## Supporting information

Supplemental Architecture Description

## Acknowledgements

We thank Ray Gonzalez for help constructing the Word-Speaker-Noise dataset used for training. We also thank Ray Gonzalez and Alex Durango for help running in-lab experiments, Joel Dapello for guidance on the VOneNet models, and Martin Schrimpf for help with Brain-Score evaluations. We thank Andrew Francl and Mark Saddler for advice on model training and evaluation, Alex Kell and Sam Norman-Haignere for help with the fMRI data analysis, and Malinda McPherson for help with Amazon Turk experiment design and statistics decisions. This work was supported by NSF grant no. BCS-1634050 to J.H.M., National Institutes of Health grant no. R01DC017970 to J.H.M., a DOE CSGF fellowship under grant no. DE-FG02-97ER25308 to J.F., and a Friends of the McGovern Institute Fellowship to J.F.

## Footnotes

1 A preliminary version of some of the experiments described here was presented in a conference paper (36).

