## Supplemental Figures for "Model metamers illuminate divergences between biological and artificial neural networks"

### Supplementary Figures

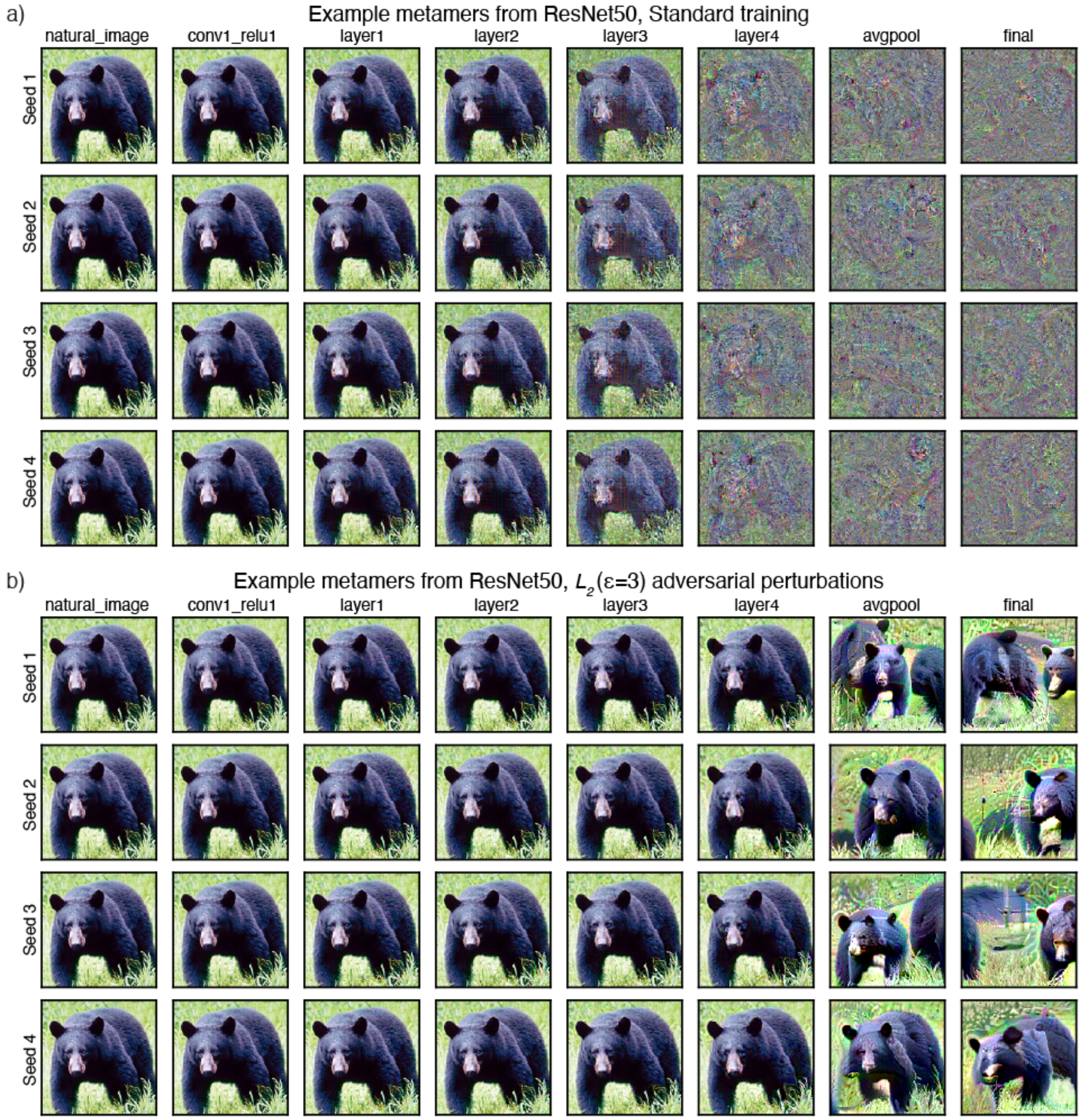


***Supplementary Figure 1.*** Additional model metamers generated from four different white noise initializations for the (a) Standard ResNet50 and (b) an adversarially trained ResNet50.

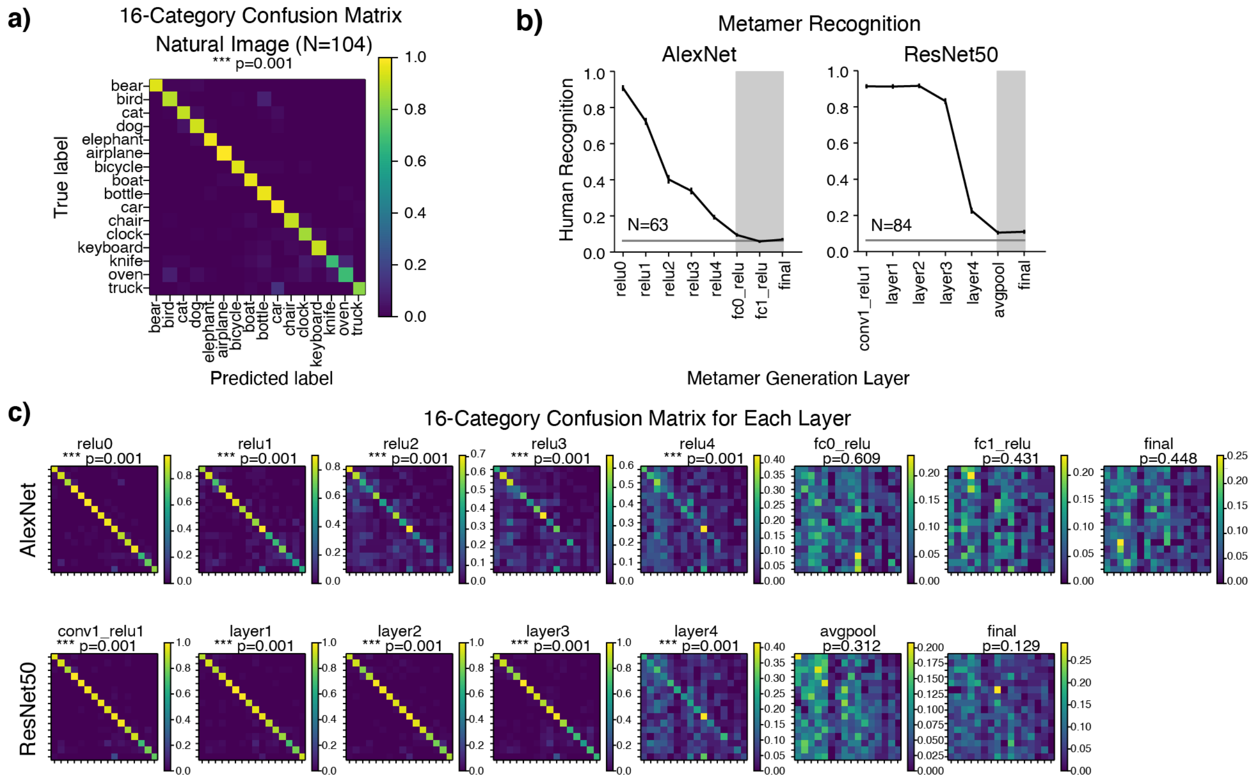


***Supplementary Figure 2.*** Analysis of consistency of human recognition errors for model metamers. (a) 16-way confusion matrix for natural images. Here and in b and c, results incorporate human responses from all experiments that contained the AlexNet Standard architecture or the ResNet50 Standard architecture. (b) Human recognizability of model metamers from different stages of AlexNet and ResNet50 models. Stages whose confusions were not consistent across splits of human observers are noted by the shaded region (see c for details of analysis). The stages for which recognition is near chance show inconsistent confusion patterns, ruling out the possibility that the chance levels of recognition are driven by systematic errors (e.g. consistently recognizing metamers for cats as dogs). (c) Confusion matrices for human recognition judgments of model metamers from each stage of the AlexNet and ResNet50 models (using data from all experiments that contained the AlexNet Standard architecture or the ResNet50 Standard architecture). We performed a split-half reliability analysis of the confusion matrices to determine whether the confusions were reliable across participants. We measured the correlation between the confusion matrices for splits of human participants, and assessed whether this correlation was significantly greater than 0. P-values from this analysis are given above each confusion matrix. For the later stages of each model, the confusion matrices are no more consistent than would be expected by chance, consistent with the metamers being completely unrecognizable (i.e., containing no information about the visual category of the natural image they are matched to).

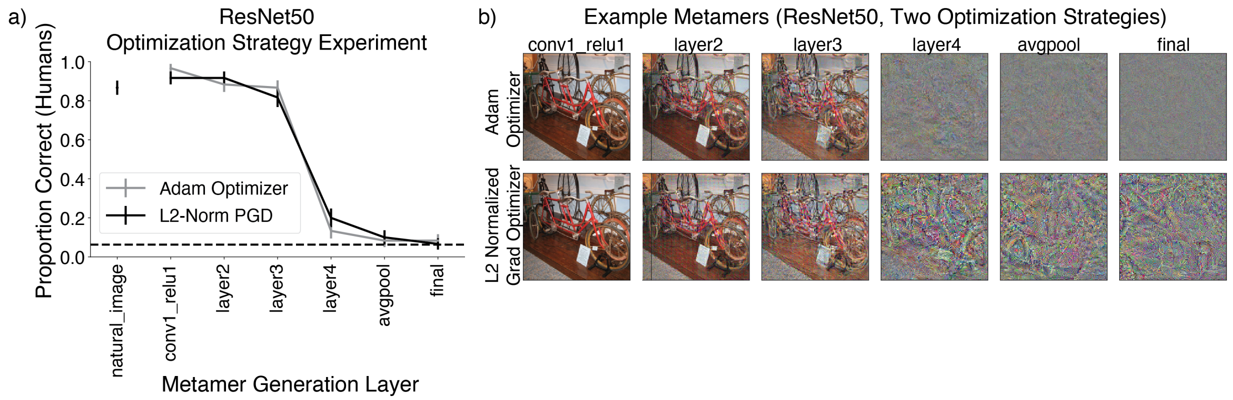


***Supplementary Figure 3***. Human recognition of visual model metamers generated from ResNet50 model with two different types of optimization strategies for metamer generation. a) Human recognition was similar for the two optimization strategies (N=10). b) Metamers are subjectively distinct for the two optimization methods, but not in ways that affect the recognition task. Experiment conducted in-lab. Error bars plot SEM across participants.

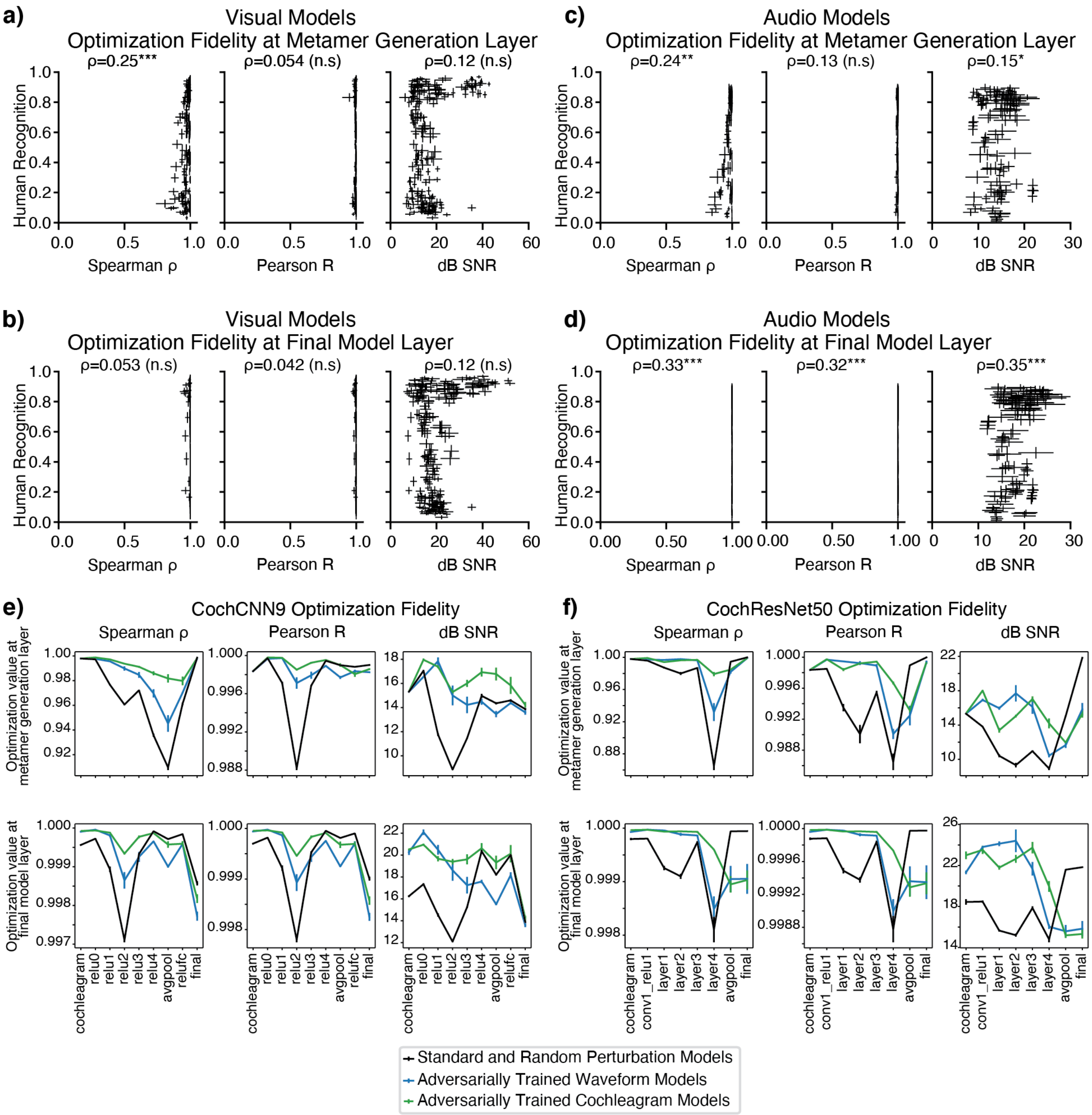


***Supplementary Figure 4.*** Optimization fidelity vs. human recognition of model metamers. (a) Optimization fidelity for visual model metamers at the metamer generation stage and (b) at the final model stage corresponding to a categorization decision (N=219). Visual models are those in Figures 2-4 and Figure 6g. Note that most data points are very close to 1 for the final stage correlation metrics (e.g. 209/219 stages exceed an average Spearman ρ of 0.99). Each point corresponds to a single stage of a single model. (c) Optimization fidelity for auditory model metamers at the metamer generation stage and (d) at the final model stage corresponding to a categorization decision. Auditory models are those in Figure 2 and Figure 5 (N=127). Note that most data points are again very close to 1 for the final stage correlation metrics (all 127 stages exceed an average Spearman ρ of 0.99). In all cases, optimization fidelity is high for both the metamer generation stage and the final stage, and human recognition is not predicted by the optimization fidelity, or only weakly correlated with the optimization fidelity for the generated model metamers (accounting for a very small fraction of the variance). Error bars on each data point are standard deviation across generated model metamers that passed the optimization criteria to be included in the psychophysical experiment. e) Final stage optimization fidelity plotted vs. model metamer generation stage for CochCNN9 auditory model and (f) CochResNet50 auditory model. Note the y axis limits, which differ across plots to show the small variations near 1 for the correlation measures. It is apparent that for any given model, some stages are somewhat less well optimized than others, but these variations do not account for the recognizability differences found in our experiments (compare these plots to the recognition plots in Figures 2 and 5).

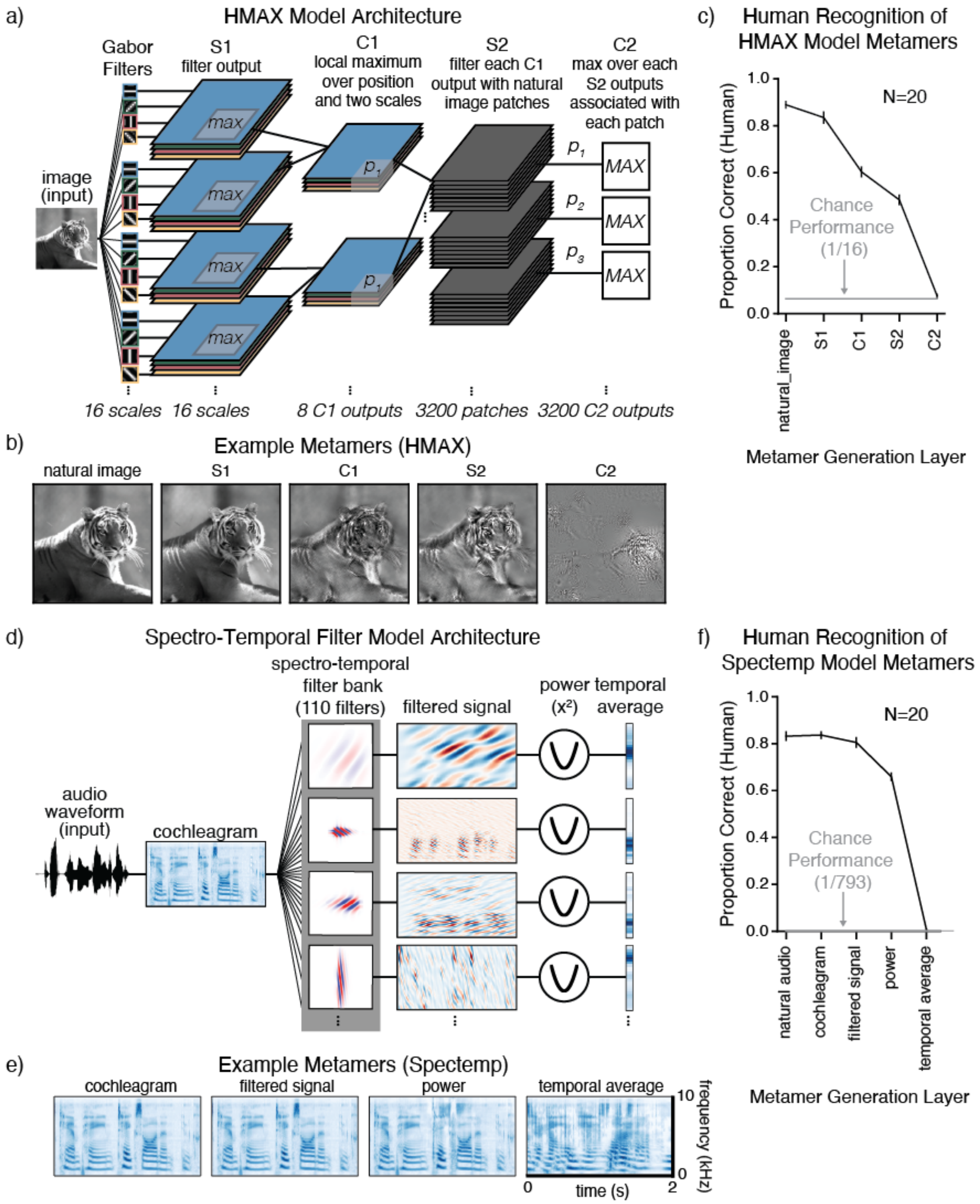


***Supplementary Figure 5.*** Metamers from classical models of sensory systems. a) Schematic of HMAX vision model, adapted from (4). b) Example HMAX model metamers. c) Human recognition of HMAX model metamers (N=20). Metamers generated from the HMAX model are recognizable at early model stages but become unrecognizable to humans by the C2 model stage. Error bars plot SEM across participants. d) Schematic of spectro-temporal auditory filterbank model (Spectemp), adapted from (56). e) Cochleagrams of example Spectemp model metamers. f) Human recognition of Spectemp model metamers (N=20). Metamers generated from the Spectemp model are recognizable at early model stages but become unrecognizable by the final (temporal average of filter power) stage. Error bars plot SEM across participants

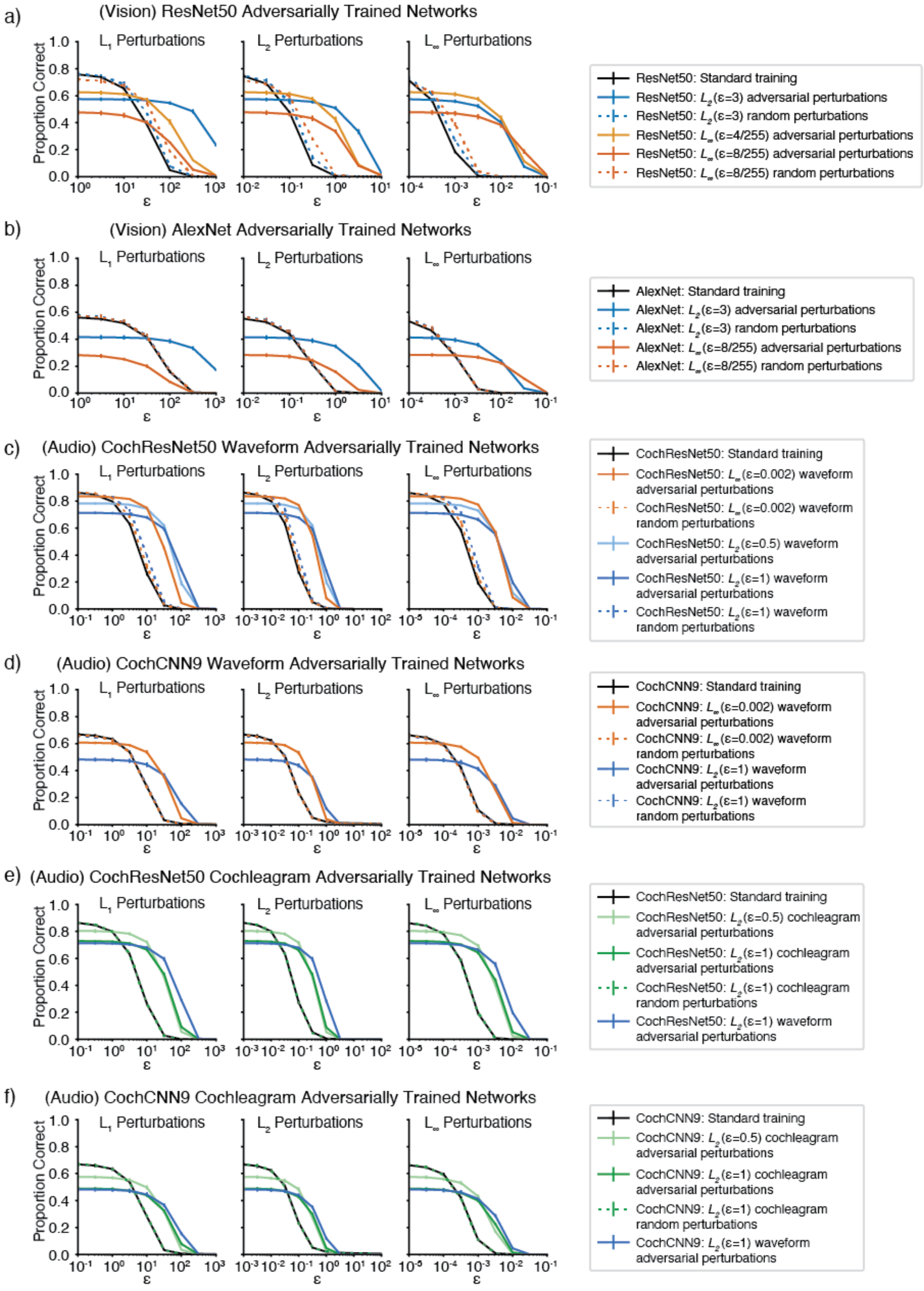
a)
(Vision) ResNet50 Adversarially Trained Networks
L1 Perturbations
L2 Perturbations
L∞ Perturbations
Proportion Correct
ε
ResNet50: Standard training
ResNet50: L2(ε=3) adversarial perturbations
ResNet50: L2(ε=3) random perturbations
ResNet50: L∞(ε=4/255) adversarial perturbations
ResNet50: L∞(ε=8/255) adversarial perturbations
ResNet50: L∞(ε=8/255) random perturbations
b)
(Vision) AlexNet Adversarially Trained Networks
AlexNet: Standard training
AlexNet: L2(ε=3) adversarial perturbations
AlexNet: L2(ε=3) random perturbations
AlexNet: L∞(ε=8/255) adversarial perturbations
AlexNet: L∞(ε=8/255) random perturbations
c)
(Audio) CochResNet50 Waveform Adversarially Trained Networks
CochResNet50: Standard training
CochResNet50: L∞(ε=0.002) waveform adversarial perturbations
CochResNet50: L∞(ε=0.002) waveform random perturbations
CochResNet50: L2(ε=0.5) waveform adversarial perturbations
CochResNet50: L2(ε=1) waveform adversarial perturbations
CochResNet50: L2(ε=1) waveform random perturbations
d)
(Audio) CochCNN9 Waveform Adversarially Trained Networks
CochCNN9: Standard training
CochCNN9: L∞(ε=0.002) waveform adversarial perturbations
CochCNN9: L∞(ε=0.002) waveform random perturbations
CochCNN9: L2(ε=1) waveform adversarial perturbations
CochCNN9: L2(ε=1) waveform random perturbations
e)
(Audio) CochResNet50 Cochleagram Adversarially Trained Networks
CochResNet50: Standard training
CochResNet50: L2(ε=0.5) cochleagram adversarial perturbations
CochResNet50: L2(ε=1) cochleagram adversarial perturbations
CochResNet50: L2(ε=1) cochleagram random perturbations
CochResNet50: L2(ε=1) waveform adversarial perturbations
f)
(Audio) CochCNN9 Cochleagram Adversarially Trained Networks
CochCNN9: Standard training
CochCNN9: L2(ε=0.5) cochleagram adversarial perturbations
CochCNN9: L2(ε=1) cochleagram adversarial perturbations
CochCNN9: L2(ε=1) cochleagram random perturbations
CochCNN9: L2(ε=1) waveform adversarial perturbations

***Supplementary Figure 6.*** Visual and auditory models become robust to adversarial attacks after adversarial training. Visual and auditory models were evaluated with $L_p$-norm white box adversarial attacks of varying strengths. a) ResNet50 models adversarially trained on ImageNet1K (same models and color scheme as Figure 5c,d). b) AlexNet models adversarially trained on ImageNet1K (same models and color scheme as Figure 5e,f). c) CochResNet50 models adversarially trained with waveform perturbations on word recognition (same models and color scheme as Figure 6b) d) CochCNN9 models adversarially trained with waveform perturbations on word recognition (same models and color scheme as Figure 6c) e) CochResNet50 models adversarially trained with cochleagram perturbations on word recognition (same models and color scheme as Figure 6f) f) CochCNN9 models adversarially trained with cochleagram perturbations on word recognition (same models and color scheme as Figure 6g). For each model and perturbation type, performance was evaluated for five random subsets of 1024 examples from the validation set. Error bars are SEM across the 5 subsets. For auditory models, adversarial perturbations for evaluation were always added to the waveform (because cochleagram perturbations are not necessarily realizable as audio signals due to overcompleteness). Adversarial training produces robustness to adversarial perturbations (better performance at large perturbation sizes), along with some reduction in clean accuracy, as is typical for adversarially trained models (129). Training with random perturbations typically produced similar results as standard training, as expected. In many plots, the results for random-perturbation training (dotted lines) overlap with those for standard training (black line).

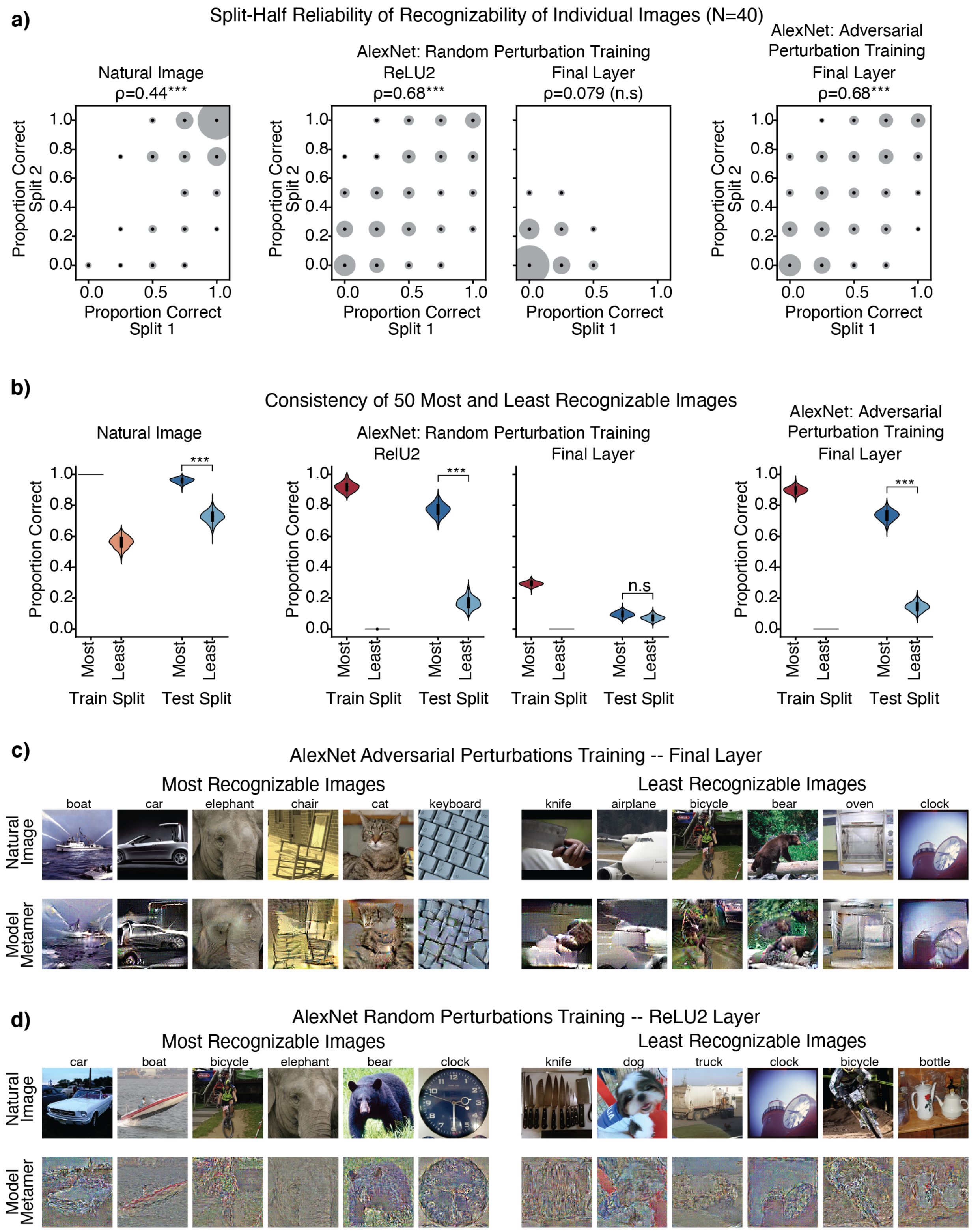


***Supplementary Figure 7.*** Analysis of human recognition of individual images from select stages of AlexNet trained with random or adversarial perturbations. To collect enough data to analyze performance on individual images, an experiment was run with only four conditions: natural images, the ReLU2 and final stages for random perturbation AlexNet and the final stage for adversarial perturbation AlexNet (N=40).

The ReLU2 stage of the Random perturbation AlexNet and the Final stage of the Adversarial perturbation AlexNet had similarly recognizable model metamers (Figure 4c) while metamers from the Final stage of the Random perturbation AlexNet were predominantly unrecognizable to humans. (a) Consistency of recognizability of individual images across splits of participants. Graph plots the proportion correct for individual stimuli for one random split. The size of the points represents the number of images at that particular value. Correlation values for each stage were determined by averaging the Spearman ρ over 1000 random splits of participants (p-values were computed non-parametrically by shuffling the participant responses for each condition, and computing the number of times the true Spearman ρ averaged across splits was lower than the shuffled correlation value). Images were only included that had at least 4 trials in each split of participants, and we only included four of the responses in the average, to avoid having some images exert more influence on the result than others (resulting in quantized values for proportion correct). Recognizability of individual images was reliable for the Natural Images, ReLU2 of AlexNet with random perturbation training and the Final stage of AlexNet with adversarial perturbation training. By contrast, recognizability of individual metamers from the Final stage of AlexNet with random perturbations showed no consistency across participants. (b) Analysis of responses for the most and least recognizable images. Using one half of the participants, we selected the 50 images that had the highest recognition score and the 50 images that had the lowest recognition score ("train" split). We then measured the recognizability for these most and least recognizable images in the second half of participants ("test" split). We analyzed 1000 random splits of the participants and p-values were computed by measuring the number of times the "most" recognizable images had a higher recognition score than the "least" recognizable images. Graph shows violin plots of the test split results for the 1000 splits. Images were only included in the analysis if at least 4 participants in each split responded to the example. The difference between the most and least recognizable metamers for the two model stages that have above-chance recognizability replicates across splits, indicating that human observers exhibit agreement on which metamers are recognizable. (c) Example natural images and model metamers for 5 images that were the "most" and "least" recognizable for the final stage of AlexNet with adversarial perturbation training and (d) for the ReLU2 stage of AlexNet with random perturbation training (as evaluated with data pooled across all participants). All images shown had at least 8 responses across participants for both the Natural Image and Model Metamer condition, had 100% correct responses for the natural image condition, and had 100% correct (for "most" recognizable images) or 0% correct (for "least" recognizable images).

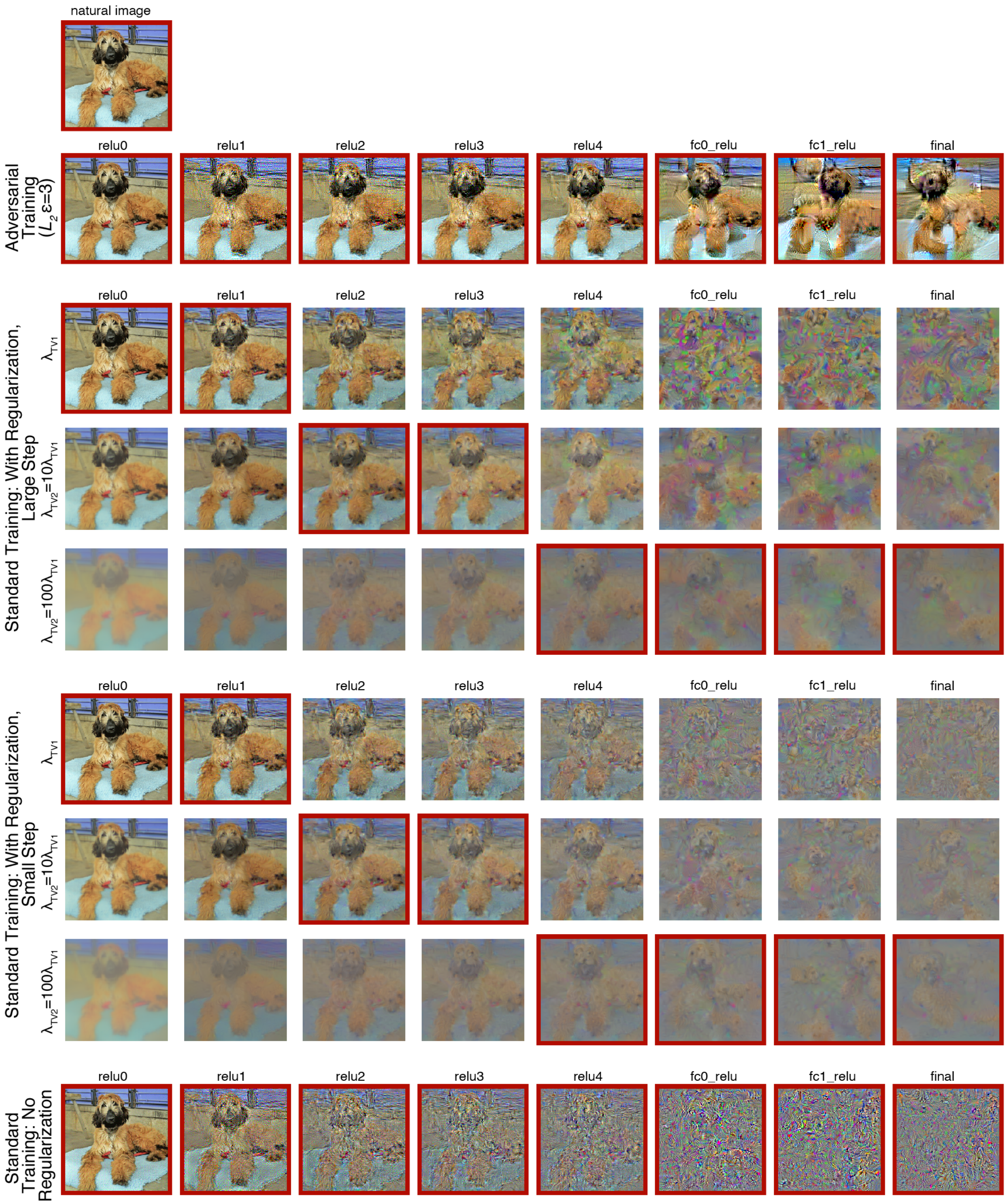


***Supplementary Figure 8.*** Examples of model metamers generated with regularization terms for smoothness and image range. Three coefficients for the smoothness regularizer were used. Red outlines are present on conditions that were used in the human classification experiment (chosen to maximize the recognizability, and to match the choices in the original paper that introduced this type of regularization (Mahendran et al. 2015)).

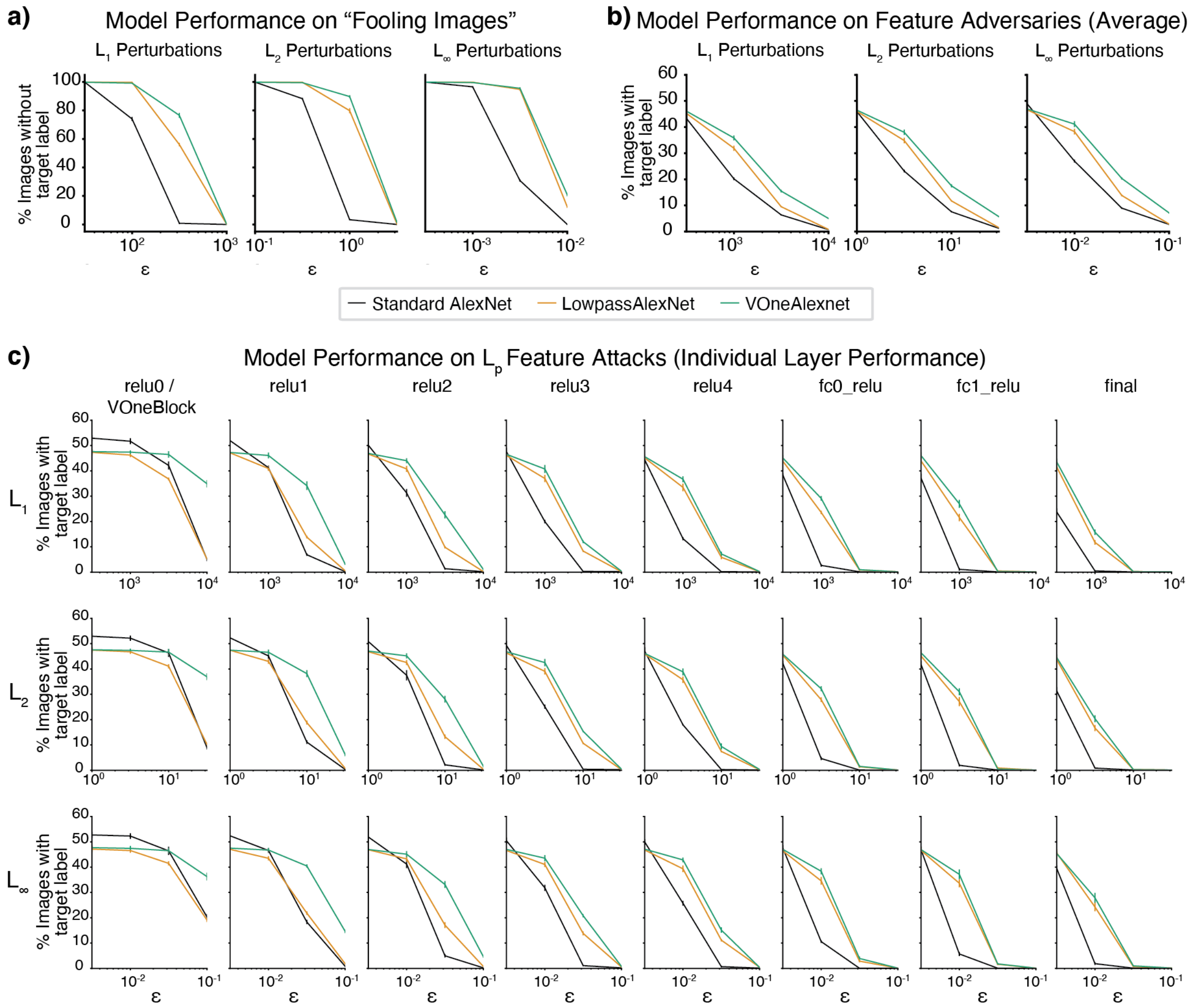


***Supplementary Figure 9.*** Adversarial robustness of VOneAlexNet and LowPassAlexNet to different types of adversarial attacks. (a) Adversarial robustness to “Fooling Images”. Fooling images are constructed by initializing with random noise (the same noise type used for initialization during model metamer generation) and making small $L_p$ constrained perturbations to cause the model to classify the noise as a particular target class. LowpassAlexNet and VOneAlexNet are more robust than the standard AlexNet for all perturbation types (main effect of architecture $F(1,8)>6787.0$, $p<0.031$, $\eta^2_p>0.999$, for all adversarial perturbation types for both VOneAlexNet and LowPassAlexNet), and although there was a main effect of architecture between LowPassAlexNet and VOneAlexNet, VOneAlexNet was more robust (main effect of architecture for perturbations, $F(1,8)>98.6$, $p<0.031$, $\eta^2_p>0.924$ for all perturbation types). Error bars plot SEM across five sets of target labels. (b) Adversarial robustness to feature adversaries. Feature adversaries are constructed by perturbing a natural “source” image so that it yields model activations (at a particular model stage) that are close to those evoked by a different natural “target” image, while constraining the perturbed image to remain within a small distance from the original natural image in pixel space. The robustness measure plotted here is averaged across adversaries generated for all stages of a model. LowpassAlexNet and VOneAlexNet are more robust than the standard AlexNet for all perturbation types (main effect of architecture $F(1,8)>90.8$, $p<0.031$, $\eta^2_p>0.919$, for all adversarial perturbation types for both VOneAlexNet and LowPassAlexNet), and although there was a main effect of architecture between LowPassAlexNet and VOneAlexNet, VOneAlexNet was more robust (main effect of architecture for perturbations, $F(1,8)>69.0$, p<0.031, $\eta^2_p$>0.895 for all perturbation types). Here and in (c), Error bars plot SEM across five samples of target and source images. (c) Performance on feature adversaries for each stage of the models, used to obtain the average curve in (b). LowpassAlexNet is not more robust to any type of adversarial example, even though it has more recognizable model metamers as shown in Figure 6g, suggesting that metamers reveal a different type of model discrepancy than that revealed with typical metrics adversarial robustness.

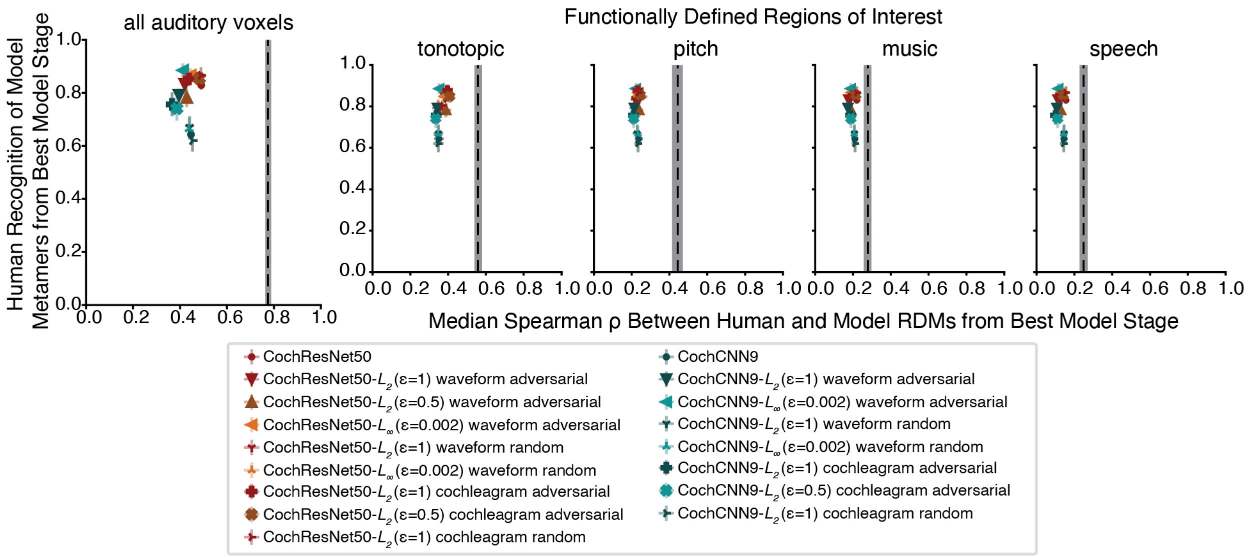


***Supplementary Figure 10.*** RSA analysis of auditory fMRI data. The median Spearman ρ between the RDM from fMRI activations to natural sounds (N=8)_and the RDM from model activations at the best model stage as determined with held out data, compared with the metamer recognition by humans at this chosen model stage. The dashed black line shows the upper bound on the RDM similarity that could be measured given the data reliability, estimated by comparing a participant's RDM with the average of the RDMs from each of the other participants. Error bars are SEM across participants. The correlation between metamer recognizability and the human-model RDM similarity was statistically significant for one ROI (all: ρ=0.02, p=.93; tonotopic: ρ=0.60, p=.011; pitch: ρ=0.06, p=.81; music: ρ=0.10, p=.70; speech: ρ=0.12, p=.66), but it was again well below the presumptive noise ceiling (which ranged from ρ=0.79 to ρ=0.89, depending on the ROI), and would not survive Bonferroni correction. We also note that the variation in metamer recognizability across models is substantially greater than the variation in RDM similarity, indicating that metamers better differentiate this set of models than does the RDM similarity with this fMRI dataset.

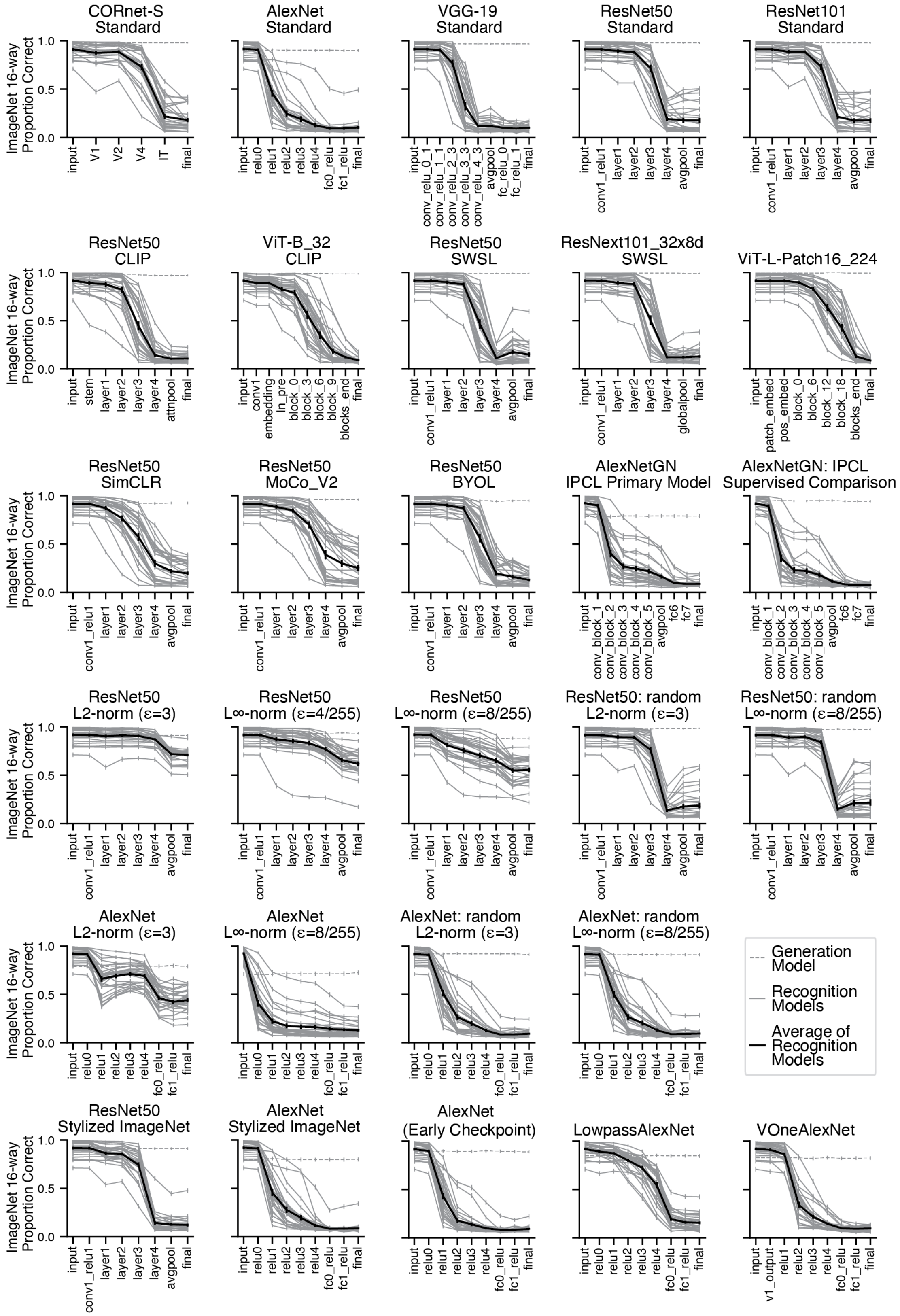
ImageNet 16-way Proportion Correct
CORnet-S Standard
AlexNet Standard
VGG-19 Standard
ResNet50 Standard
ResNet101 Standard
ResNet50 CLIP
ViT-B_32 CLIP
ResNet50 SWSL
ResNext101_32x8d SWSL
ViT-L-Patch16_224
ResNet50 SimCLR
ResNet50 MoCo_V2
ResNet50 BYOL
AlexNetGN IPCL Primary Model
AlexNetGN: IPCL Supervised Comparison
ResNet50 L2-norm (ε=3)
ResNet50 L∞-norm (ε=4/255)
ResNet50 L∞-norm (ε=8/255)
ResNet50: random L2-norm (ε=3)
ResNet50: random L∞-norm (ε=8/255)
AlexNet L2-norm (ε=3)
AlexNet L∞-norm (ε=8/255)
AlexNet: random L2-norm (ε=3)
AlexNet: random L∞-norm (ε=8/255)
Generation Model
Recognition Models
Average of Recognition Models
ResNet50 Stylized ImageNet
AlexNet Stylized ImageNet
AlexNet (Early Checkpoint)
LowpassAlexNet
VOneAlexNet

***Supplementary Figure 11.*** Image model recognition of metamers generated from other models. Metamers were generated from the model indicated in the plot title and recognition was measured by presenting the metamers to other models (each grey line on the plot corresponds to one recognition model). Error bars on individual model curves are bootstrapped SEM from the model predictions. Error bars on the average recognition curve is the SEM across recognition models. Model metamers from deep stages tend to be unrecognizable to other models, but models trained with adversarial perturbations or with architecturally fixed lowpass filtering operations have metamers that are more recognizable by other models.

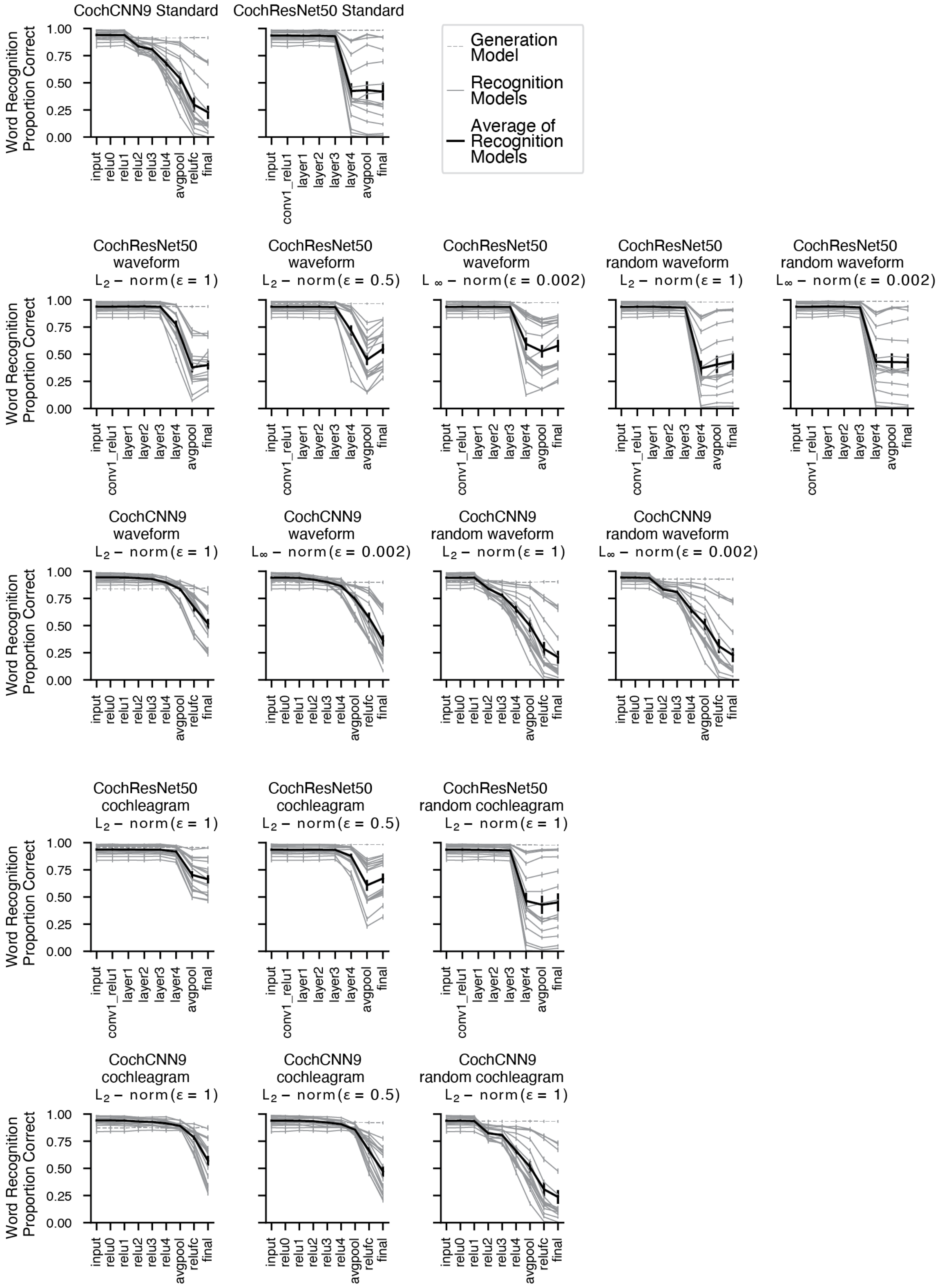

CochCNN9 Standard
CochResNet50 Standard
Generation Model
Recognition Models
Average of Recognition Models
Word Recognition Proportion Correct
CochResNet50 waveform $L_2$ − norm(ε = 1)
CochResNet50 waveform $L_2$ − norm(ε = 0.5)
CochResNet50 waveform $L_\infty$ − norm(ε = 0.002)
CochResNet50 random waveform $L_2$ − norm(ε = 1)
CochResNet50 random waveform $L_\infty$ − norm(ε = 0.002)
CochCNN9 waveform $L_2$ − norm(ε = 1)
CochCNN9 waveform $L_\infty$ − norm(ε = 0.002)
CochCNN9 random waveform $L_2$ − norm(ε = 1)
CochCNN9 random waveform $L_\infty$ − norm(ε = 0.002)
CochResNet50 cochleagram $L_2$ − norm(ε = 1)
CochResNet50 cochleagram $L_2$ − norm(ε = 0.5)
CochResNet50 random cochleagram $L_2$ − norm(ε = 1)
CochCNN9 cochleagram $L_2$ − norm(ε = 1)
CochCNN9 cochleagram $L_2$ − norm(ε = 0.5)
CochCNN9 random cochleagram $L_2$ − norm(ε = 1)
0.00 0.25 0.50 0.75 1.00
input relu0 relu1 relu2 relu3 relu4 avgpool relufc final
input conv1_relu1 layer1 layer2 layer3 layer4 avgpool final

***Supplementary Figure 12.*** Auditory model recognition of metamers generated from other models. Metamers were generated from the model indicated in the plot title and recognition was measured by presenting the metamers to other models (each grey line on the plot corresponds to one recognition model). Error bars on individual model curves are bootstrapped SEM from the model predictions. Error bars on the average recognition curve is the SEM across recognition models. As with the image models, model metamers from deep stages tend to be unrecognizable to other models, but models trained with adversarial perturbations have metamers that are more recognizable by other models.

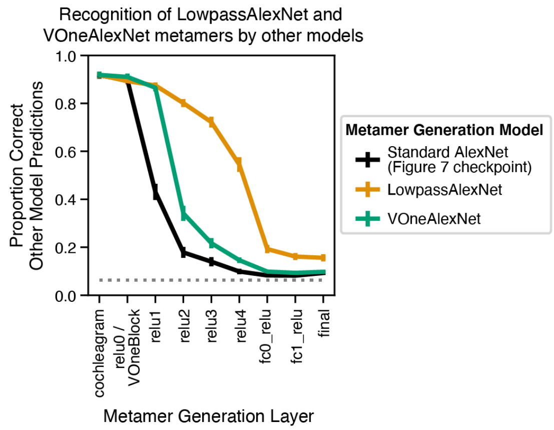


***Supplementary Figure 13.*** Other-model recognition of LowpassAlexNet and VOneAlex model metamers (averaged across all visual recognition models as in Figure 9b, but with Figure 7 models as the metamer generation models). Error bars are SEM over the recognition models. Model metamers for the LowpassAlexNet are much more recognizable to other models when generated from intermediate model stages, suggesting that the model architecture eliminates some model-specific invariances that are present in the Standard AlexNet or the VOneAlexNet.
