## Supplemental Architecture Description for "Model metamers illuminate divergences between biological and artificial neural networks"

### Supplemental Information: Model Architecture Descriptions

The general structure of the neural network architectures used in the paper are documented in this file, including names for the model stages that were used for metamer generation and included on figures. Model stage names and number of features are only given for the model stages used in metamer experiments (bolded in the tables below). All output shapes are specified without a batch dimension.

#### *CORnet-S*

The CORnet-S architecture was proposed in (73) and contains recurrent and skip connections motivated by brain and behavioral data.

| Model Stage Name | PyTorch Operation | Output Shape | Number of Features |
|---|---|---|---|
| **natural_image** | **input** | **(3,224,224)** | **150528** |
| | Conv2d(3, 64, kernel_size=7, stride=2, padding=3, bias=False) | (64, 112, 112) | |
| | BatchNorm2d(64) | (64, 112, 112) | |
| | ReLU | (64, 112, 112) | |
| | MaxPool2d(kernel_size=3, stride=2, padding=1) | (64, 56, 56) | |
| | Conv2d(64, 64, kernel_size=3, stride=1, padding=1, bias=False) | (64, 56, 56) | |
| | BatchNorm2d(64) | (64, 56, 56) | |
| **V1** | **ReLU** | **(64, 56, 56)** | **200704** |
| | Conv2d(64, 128, kernel_size=1, stride=1, bias=False) | (128, 56, 56) | |
| **V2** | **CORblock_S(128, scale=4, time=2)** | **(128, 28, 28)** | **100352** |
| | Conv2d(128, 256, kernel_size=1, stride=1, bias=False) | (256, 28, 28) | |
| **V4** | **CORblock_S(256, scale=4, time=4)** | **(256, 14, 14)** | **50176** |
| | Conv2d(256, 512, kernel_size=1, stride=1, bias=False) | (512, 14, 14) | |
| **IT** | **CORblock_S(512, scale=4, time=2)** | **(512, 7, 7)** | **25088** |

| | | | |
|---|---|---|---|
| | AdaptiveAvgPool2d(1) | (512, 1, 1) | |
| **final** | **Linear(512, 1000)** | **(1000)** | **1000** |

The CORblock_S(channels, scale, t) components of the architecture have the following structure:

| |
|---|
| 1. Input (x) |
| 2. Conv2d(channels, channels * scale, kernel_size=1, bias=False) |
| 3. BatchNorm2d(channels * scale) |
| 4. ReLU |
| 5. t=0: Conv2d(channels * scale, channels * scale, kernel_size=3, stride=2, padding=1, bias=False)<br>t != 0: Conv2d(channels * scale, channels * scale, kernel_size=3, stride=1, padding=1, bias=False) |
| 6. BatchNorm2d(channels * scale) |
| 7. ReLU |
| 8. Conv2d(channels * scale, channels, kernel_size=1, bias=False) |
| 9. BatchNorm2d(channels) |
| 10. Process skip connection (on input, x):<br>t = 0: x processed with<br>a. 1x1 Conv2d(channels, channels, kernel_size=1, stride=2, bias=False)<br>b. BatchNorm2d(channels)<br><br>t != 0: x processed with Identity() |
| 11. Add output from (9) to output from (10) |
| 12. (Output) ReLU |

To implement recurrent connections, the input passes through this CORblock_S block \`t\` times, where the convolutional layers share weights for each timestep \`t\` but the batch normalization layers have unique learnable weights for each \`t\`. The first pass \`t=0\` through the block contains additional downsampling of the residual connection and in the second convolution.

***VGG19***

The VGG19 architecture was proposed in (39) and has 16 convolutional layers, 5 max pooling layers, and 2 fully connected layers (plus one classification layer).

| Model Stage Name | PyTorch Operation | Output Shape | Number of Features |
|---|---|---|---|
| **natural_image** | **input** | **(3,224,224)** | **150528** |
| | Conv2d(3, 64, kernel_size=3, padding=1) | (64, 224, 224) | |

| | | | |
|---|---|---|---|
| | ReLU | (64, 224, 224) | |
| | Conv2d(64, 64, kernel_size=3, padding=1) | (64, 224, 224) | |
| **conv_relu_0_1** | **ReLU** | **(64, 224, 224)** | **3211264** |
| | MaxPool2d(kernel_size=2, stride=2) | (64, 112, 112) | |
| | Conv2d(64, 128, kernel_size=3, padding=1) | (128, 112, 112) | |
| | ReLU | (128, 112, 112) | |
| | Conv2d(128, 128, kernel_size=3, padding=1) | (128, 112, 112) | |
| **conv_relu_1_1** | **ReLU** | **(128, 112, 112)** | **1605632** |
| | MaxPool2d(kernel_size=2, stride=2) | (128, 56, 56) | |
| | Conv2d(128, 256, kernel_size=3, padding=1) | (256, 56, 56) | |
| | ReLU | (256, 56, 56) | |
| | Conv2d(256, 256, kernel_size=3, padding=1) | (256, 56, 56) | |
| | ReLU | (256, 56, 56) | |
| | Conv2d(256, 256, kernel_size=3, padding=1) | (256, 56, 56) | |
| | ReLU | (256, 56, 56) | |
| | Conv2d(256, 256, kernel_size=3, padding=1) | (256, 56, 56) | |
| **conv_relu_2_3** | **ReLU** | **(256, 56, 56)** | **802816** |
| | MaxPool2d(kernel_size=2, stride=2) | (256, 28, 28) | |
| | Conv2d(256, 512, kernel_size=3, padding=1) | (512, 28, 28) | |
| | ReLU | (512, 28, 28) | |
| | Conv2d(512, 512, kernel_size=3, padding=1) | (512, 28, 28) | |
| | ReLU | (512, 28, 28) | |

| | | | |
|---|---|---|---|
| | Conv2d(512, 512, kernel_size=3, padding=1) | (512, 28, 28) | |
| | ReLU | (512, 28, 28) | |
| | Conv2d(512, 512, kernel_size=3, padding=1) | (512, 28, 28) | |
| **conv_relu_3_3** | **ReLU** | **(512, 28, 28)** | **401408** |
| | MaxPool2d(kernel_size=2, stride=2) | (512, 14, 14) | |
| | Conv2d(512, 512, kernel_size=3, padding=1) | (512, 14, 14) | |
| | ReLU | (512, 14, 14) | |
| | Conv2d(512, 512, kernel_size=3, padding=1) | (512, 14, 14) | |
| | ReLU | (512, 14, 14) | |
| | Conv2d(512, 512, kernel_size=3, padding=1) | (512, 14, 14) | |
| | ReLU | (512, 14, 14) | |
| | Conv2d(512, 512, kernel_size=3, padding=1) | (512, 14, 14) | |
| **conv_relu_4_3** | **ReLU** | **(512, 14, 14)** | **100352** |
| | MaxPool2d(kernel_size=2, stride=2) | (512, 7, 7) | |
| **avgpool** | **AdaptiveAvgPool2d((7, 7)) (note: this is equal to the output of the above MaxPool2d)** | **(512, 7, 7)** | **25088** |
| | Linear(512 * 7 * 7, 4096) | (4096) | |
| **fc_relu_0** | **ReLU** | **(4096)** | **4096** |
| | Dropout(p=0.5) | (4096) | |
| | Linear(4096, 4096) | (4096) | |
| **fc_relu_1** | **ReLU** | **(4096)** | **4096** |
| | Dropout(p=0.5) | (4096) | |
| **final** | **Linear(4096, 1000)** | **(1000)** | **1000** |

***ResNet50***

The ResNet50 architecture was proposed in (40) and has 48 convolutional layers with 1 max pooling layer and 1 average pooling layer (plus one classification layer). It has 4 residual blocks (ResNetBlocks below) which have skip (or shortcut) connections. This architecture is used for all models labled as “ResNet50” with the exception of ResNet50: CLIP, which has modifications described below.

Layer names and number of features are only given for the layers used in metamer experiments.

| Model Stage Name | PyTorch Operation | Output Shape | Number of Features |
|---|---|---|---|
| **natural_image** | **input** | **(3,224,224)** | **150528** |
| | Conv2d(3, 64, kernel_size=7, stride=2, padding=3, bias=False) | (64, 112, 112) | |
| | BatchNorm2d(64) | (64, 112, 112) | |
| **conv1_relu1** | **ReLU** | **(64, 112, 112)** | **802816** |
| | MaxPool2d(kernel_size=3, stride=2, padding=1) | (64, 56, 56) | |
| **layer1** | **ResNetBlock(inplanes=64, planes=64, num_blocks=3, stride=1)** | **(256, 56, 56)** | **802816** |
| **layer2** | **ResNetBlock(inplanes=256, planes=128, num_blocks=4, stride=2)** | **(512, 28, 28)** | **401408** |
| **layer3** | **ResNetBlock(inplanes=512, planes=256, num_blocks=6, stride=2)** | **(1024, 14, 14)** | **200704** |
| **layer4** | **ResNetBlock(inplanes=1024, planes=512, num_blocks=3, stride=2)** | **(2048, 7, 7)** | **100352** |
| **avgpool** | **AdaptiveAvgPool2d(1,1)** | **(2048, 1, 1)** | **2048** |
| **final** | **Linear(2048, 1000)** | **(1000)** | **1000** |

The ResNetBlock components of the architecture have the following structure:

| |
|---|
| 1. input (x) |
| 2. 1x1 Conv2d(inplanes, planes, stride=1) |
| 3. BatchNorm2d(planes) |
| 4. ReLU |
| 5. 3x3 Conv2d (planes, planes, stride=1) |
| 6. BatchNorm2d(planes) |

| |
|---|
| 7. ReLU |
| 8. 1x1 Conv2d (planes, planes * expansion, stride=1) |
| 9. BatchNorm2d(planes) |
| 10. Residual connection on x (if inplanes !=planes * expansion): 1x1 Conv2D (inplanes, planes * expansion, stride) |
| 11. Residual connection on x (if inplanes !=planes * expansion): BatchNorm2d(planes * expansion) |
| 12. Add output from (9) to output from (11) |
| 13. (Output) ReLU |

Multiple of these residual blocks (num_blocks) are stacked together to form a single ResNetBlock. The expansion factor was set to four for all layers (expansion=4).

*ResNet50: CLIP modifications*

The CLIP model with a ResNet50 visual encoder obtained from https://github.com/openai/CLIP had three "stem" convolutions as opposed to one, with an average pool instead of a max pool. It also prepends an avgpool to convolutions with stride > 1 within the residual blocks, and uses a final attention pooling layer rather than average pooling. Full architecture details below.

Layer names and number of features are only given for the layers used in metamer experiments.

| Model Stage Name | PyTorch Operation | Output Shape | Number of Features |
|---|---|---|---|
| **natural_image** | **input** | **(3,224,224)** | **150528** |
| | Conv2d(3, 32, kernel_size=3, stride=2, padding=1, bias=False) | (32, 112, 112) | |
| | BatchNorm2d(32) | (32, 112, 112) | |
| | ReLU | (32, 112, 112) | |
| | Conv2d(32, 32, kernel_size=3, stride=1, padding=1, bias=False) | (32, 112, 112) | |
| | BatchNorm2d(32) | (32, 112, 112) | |
| | ReLU | (32, 112, 112) | |
| | Conv2d(32, 64, kernel_size=3, stride=1, padding=1, bias=False) | (64, 112, 112) | |
| | BatchNorm2d(64) | (64, 112, 112) | |
| | ReLU | (64, 112, 112) | |

| | | | |
|---|---|---|---|
| **stem** | **AvgPool2d(kernel_size=2, stride=2, padding=0)** | **(64, 56, 56)** | **200704** |
| **layer1** | **ResNetBlock(inplanes=64, planes=64, num_blocks=3, stride=1)** | **(256, 56, 56)** | **802816** |
| **layer2** | **ResNetBlock(inplanes=256, planes=128, num_blocks=4, stride=2)** | **(512, 28, 28)** | **401408** |
| **layer3** | **ResNetBlock(inplanes=512, planes=256, num_blocks=6, stride=2)** | **(1024, 14, 14)** | **200704** |
| **layer4** | **ResNetBlock(inplanes=1024, planes=512, num_blocks=3, stride=2)** | **(2048, 7, 7)** | **100352** |
| **attnpool** | **AttentionPool2D(spatial_dim=7, embed_dim=2048, num_heads=32, output_dim=1024)** | **(1024)** | **1024** |

The ResNetBlock components of the architecture have the following structure:

| |
|---|
| 1. input (x) |
| 2. 1x1 Conv2d(inplanes, planes, stride=1, bias=False) |
| 3. BatchNorm2d(planes) |
| 4. ReLU |
| 5. 3x3 Conv2d (planes, planes, stride=1, bias=False) |
| 6. BatchNorm2d(planes) |
| 7. ReLU |
| 8. Downsampling after second convolution (if stride > 1): AvgPool2d(kernel_size=stride, stride=stride, padding=0) |
| 9. 1x1 Conv2d (planes, planes * expansion, stride=1, bias=False) |
| 10. BatchNorm2d(planes) |
| 11. Residual connection on x (if inplanes !=planes * expansion):<br>a. (if stride>1): AvgPool2d(kernel_size=stride, stride=stride, padding=0)<br>b. 1x1 Conv2D (inplanes, planes * expansion, stride=1, bias=False)<br>c. BatchNorm2d(planes * expansion) |
| 12. Add output from (10) to output from (11) |
| 13. (Output) ReLU |

Multiple of these residual blocks (num_blocks) are stacked together to form a single ResNetBlock. The expansion factor was set to four for all layers (expansion=4).

***ResNet101***

The ResNet101 architecture was proposed in (40) and has 99 convolutional layers with 1 max pooling layer and 1 average pooling layer (plus one classification layer).It has 4 residual blocks (ResNetBlocks below) which have skip (or shortcut) connections. The main difference compared to the ResNet50 model is the increased depth of the third ResNetBlock (layer3) in the model.

| Model Stage Name | PyTorch Operation | Output Shape | Number of Features |
|---|---|---|---|
| **natural_image** | **input** | **(3,224,224)** | **150528** |
| | Conv2d(3, 64, kernel_size=7, stride=2, padding=3, bias=False) | (64, 112, 112) | |
| | BatchNorm2d(64) | (64, 112, 112) | |
| **conv1_relu1** | **ReLU** | **(64, 112, 112)** | **802816** |
| | MaxPool2d(kernel_size=3, stride=2, padding=1) | (64, 56, 56) | |
| **layer1** | **ResNetBlock(inplanes=64, planes=64, num_blocks=3, stride=1)** | **(256, 56, 56)** | **802816** |
| **layer2** | **ResNetBlock(inplanes=256, planes=128, num_blocks=4, stride=2)** | **(512, 28, 28)** | **401408** |
| **layer3** | **ResNetBlock(inplanes=512, planes=256, num_blocks=23, stride=2)** | **(1024, 14, 14)** | **200704** |
| **layer4** | **ResNetBlock(inplanes=1024, planes=512, num_blocks=3, stride=2)** | **(2048, 7, 7)** | **100352** |
| **avgpool** | **AdaptiveAvgPool2d(1,1)** | **(2048, 1, 1)** | **2048** |
| **final** | **Linear(2048, 1000)** | **(1000)** | **1000** |

The ResNetBlock components of the architecture have the following structure (same as in ResNet50 model):

| |
|---|
| 1. input (x) |
| 2. 1x1 Conv2d(inplanes, planes, stride=1) |
| 3. BatchNorm2d(planes) |
| 4. ReLU |
| 5. 3x3 Conv2d (planes, planes, stride=1) |
| 6. BatchNorm2d(planes) |

| |
|---|
| 7. ReLU |
| 8. 1x1 Conv2d (planes, planes * expansion, stride=1) |
| 9. BatchNorm2d(planes) |
| 10. Residual connection on x (if inplanes !=planes * expansion): 1x1 Conv2D (inplanes, planes * expansion, stride) |
| 11. Residual connection on x (if inplanes !=planes * expansion): BatchNorm2d(planes * expansion) |
| 12. Add output from (9) to output from (11) |
| 13. (Output) ReLU |

Multiple of these residual blocks (num_blocks) are stacked together to form a single ResNetBlock. The expansion factor was set to four for all layers (expansion=4).

***AlexNet***

AlexNet was proposed in (41) and consists of 5 convolutional layers, 3 max-pooling layers, and 2 fully connected layers (plus one classification layer).

Model stage names and number of features are only given for the stages used in metamer experiments.

| Model Stage Name | PyTorch Operation | Output Shape | Number of Features |
|---|---|---|---|
| **natural_image** | **input** | **(3,224,224)** | **150528** |
| | Conv2d(3, 64, kernel_size=11, stride=4, padding=2) | (64, 55, 55) | |
| **relu0** | **ReLU** | **(64, 55, 55)** | **193600** |
| | MaxPool2d(kernel_size=3, stride=2) | (64, 27, 27) | |
| | Conv2d(64, 192, kernel_size=5, padding=2) | (192, 27, 27) | |
| **relu1** | **ReLU** | **(192, 27, 27)** | **139968** |
| | MaxPool2d(kernel_size=3, stride=2) | (192, 13, 13) | |
| | Conv2d(192, 384, kernel_size=3, padding=1) | (384, 13, 13) | |
| **relu2** | **ReLU** | **(384, 13, 13)** | **64896** |
| | Conv2d(384, 256, kernel_size=3, padding=1) | (256, 13, 13) | |
| **relu3** | **ReLU** | **(256, 13, 13)** | **43264** |

| | | | |
|---|---|---|---|
| | Conv2d(256, 256, kernel_size=3, padding=1) | (256, 13, 13) | |
| **relu4** | **ReLU** | **(256, 13, 13)** | **43264** |
| | MaxPool2d(kernel_size=3, stride=2) | (256, 6, 6) | |
| | Dropout(p=0.5) | (9216) | |
| | Linear(256 * 6 * 6, 4096) | (4096) | |
| **fc0_relu** | **ReLU** | **(4096)** | **4096** |
| | Dropout(p=0.5) | (4096) | |
| | Linear(4096, 4096) | (4096) | |
| **fc1_relu** | **ReLU** | **(4096)** | **4096** |
| **final** | **Linear(4096, 1000)** | **(1000)** | **1000** |

***ViT-B_32: CLIP***

ViT-B_32 is a vision transformer architecture based on that proposed in (130). The visual encoder from CLIP with this architecture was used for metamer generation.

Model stage names and number of features are only given for the stages used in metamer experiments.

| Model Stage Name | PyTorch Operation | Output Shape | Number of Features |
|---|---|---|---|
| **natural_image** | **Input** | **(3,224,224)** | **150528** |
| **conv1** | **Conv2d(3, 768, kernel_size=(32,32), stride=(32,32), padding=0, bias=False)** | **(768, 7, 7)** | **37632** |
| | Reshape(768,49) | (768,49) | |
| | Permute(1,0) | (49,768) | |
| **class_embedding** | **Concatenate(class_embedding, x)** | **(50, 768)** | **38400** |
| | Add(x, positional_embedding) | (50,768) | |
| **ln_pre** | **LayerNorm(768, eps=1e-05, elementwise_affine=True)** | **(50,768)** | **38400** |
| **block_0** | **ResidualAttentionBlock(d_model=768, n_head=12, attn_mask=None)** | **(50,768)** | **38400** |

| | | | |
|---|---|---|---|
| | ResidualAttentionBlock(d_model=768, n_head=12, attn_mask=None) | (50,768) | |
| | ResidualAttentionBlock(d_model=768, n_head=12, attn_mask=None) | (50,768) | |
| **block_3** | **ResidualAttentionBlock(d_model=768, n_head=12, attn_mask=None)** | **(50,768)** | **38400** |
| | ResidualAttentionBlock(d_model=768, n_head=12, attn_mask=None) | (50,768) | |
| | ResidualAttentionBlock(d_model=768, n_head=12, attn_mask=None) | (50,768) | |
| **block_6** | **ResidualAttentionBlock(d_model=768, n_head=12, attn_mask=None)** | **(50,768)** | **38400** |
| | ResidualAttentionBlock(d_model=768, n_head=12, attn_mask=None) | (50,768) | |
| | ResidualAttentionBlock(d_model=768, n_head=12, attn_mask=None) | (50,768) | |
| **block_9** | **ResidualAttentionBlock(d_model=768, n_head=12, attn_mask=None)** | **(50,768)** | **38400** |
| | ResidualAttentionBlock(d_model=768, n_head=12, attn_mask=None) | (50,768) | |
| **blocks_end** | **ResidualAttentionBlock(d_model=768, n_head=12, attn_mask=None)** | **(50,768)** | **38400** |
| | slice(0,:) | (768) | |
| | LayerNorm(768, eps=1e-05, elementwise_affine=True) | (768) | |
| **linear_end** | **Linear(768, 512, bias=False)** | **(512)** | **512** |

The transformer ResidualAttentionBlock components of the architecture have the following structure:

| |
|---|
| 1. input |
| 2. LayerNorm(d_model) |
| 3. MultiheadAttention(d_model, n_head, attn_mask) |
| 4. Residual: Add (1) to output of (3) |
| 5. LayerNorm(d_model) |

| |
|---|
| 6. MLP Layer:<br>a. Linear (d_model, d_model * 4)<br>b. QuickGELU<br>c. Linear (d_model * 4, d_model) |
| 7. Residual: Add output of (4) to output of (6) |

Note that an attention mask of “None” as used for the visual encoder and corresponds to full attention between the tokens.

***SWSL-ResNext101-32x8d***

The ResxNet101-32x8d architecture was proposed in (131) and has 99 convolutional layers with 1 max pooling layer and 1 average pooling layer (plus one classification layer). It has 4 residual blocks (ResNextBlocks below) which have skip (or shortcut) connections. The main difference compared to the ResNet101 model is the presence of grouped 2D convolutions for the 3x3 convolution of the residual block.

| Model Stage Name | PyTorch Operation | Output Shape | Number of Features |
|---|---|---|---|
| **natural_image** | **input** | **(3,224,224)** | **150528** |
| | Conv2d(3, 64, kernel_size=7, stride=2, padding=3, bias=False) | (64, 112, 112) | |
| | BatchNorm2d(64) | (64, 112, 112) | |
| **conv1_relu1** | **ReLU** | **(64, 112, 112)** | **802816** |
| | MaxPool2d(kernel_size=3, stride=2, padding=1) | (64, 56, 56) | |
| **layer1** | **ResNextBlock(inplanes=64, planes=64, num_blocks=3, stride=1, cardinality=32, base_width=8)** | **(256, 56, 56)** | **802816** |
| **layer2** | **ResNextBlock(inplanes=256, planes=128, num_blocks=4, stride=2, cardinality=32, base_width=8)** | **(512, 28, 28)** | **401408** |
| **layer3** | **ResNextBlock(inplanes=512, planes=256, num_blocks=23, stride=2, cardinality=32, base_width=8)** | **(1024, 14, 14)** | **200704** |
| **layer4** | **ResNextBlock(inplanes=1024, planes=512, num_blocks=3, stride=2, cardinality=32, base_width=8)** | **(2048, 7, 7)** | **100352** |
| **avgpool** | **AdaptiveAvgPool2d(1,1)** | **(2048, 1, 1)** | **2048** |
| **final** | **Linear(2048, 1000)** | **(1000)** | **1000** |

The ResNextBlock components of the architecture have the following structure:

| |
|---|
| 1. input (x) |
| 2. 1x1 Conv2d(inplanes, width, stride=1, bias=False) |
| 3. BatchNorm2d(width) |
| 4. ReLU |
| 5. 3x3 Conv2d (width, width, stride=1, groups=cardinality, bias=False) |
| 6. BatchNorm2d(width) |
| 7. ReLU |
| 8. 1x1 Conv2d (width, planes * expansion, stride=1, bias=False) |
| 9. BatchNorm2d(planes) |
| 10. Residual connection on x (if inplanes !=planes * expansion): 1x1 Conv2D (inplanes, planes * expansion, stride) |
| 11. Residual connection on x (if inplanes !=planes * expansion): BatchNorm2d(planes * expansion) |
| 12. Add output from (9) to output from (11) |
| 13. (Output) ReLU |

Where width = (floor(planes * base_width / 64 ) * cardinality) and multiple of these blocks (num_blocks) are stacked together to form a single ResNextBlock. The expansion factor was set to four for all layers (expansion=4).

***ViT_large_patch-16_224***

ViT-large_path-16_224 is a vision transformer architecture based on that proposed in (130).

Model stage names and number of features are only given for the stages used in metamer experiments.

| Model Stage Name | PyTorch Operation | Output Shape | Number of Features |
|---|---|---|---|
| **natural_image** | **Input** | **(3,224,224)** | **150528** |
| | Conv2d(3, 1024, kernel_size=(16,16), stride=(16,16), padding=0, bias=False) | (1024, 14, 14) | |
| | Reshape(768,49) | (1024,196) | |
| **patch_embed** | **Permute(1,0)** | **(196,1024)** | |
| | Concatenate(class_embedding, x) | (197,1024) | |

| | | |
|---|---|---|
| **pos_embedding** | **Add(x, positional_embedding)** | **(197, 1024)** |
| **block_0** | **TransformerBlock(dim=1024, n_head=16, qkv_bias=True)** | **(197, 1024)** |
| block_1 | TransformerBlock(dim=1024, n_head=16, qkv_bias=True) | (197,1024) |
| block_2 | TransformerBlock(dim=1024, n_head=16, qkv_bias=True) | (197, 1024) |
| block_3 | TransformerBlock(dim=1024, n_head=16, qkv_bias=True) | (197, 1024) |
| block_4 | TransformerBlock(dim=1024, n_head=16, qkv_bias=True) | (197, 1024) |
| block_5 | TransformerBlock(dim=1024, n_head=16, qkv_bias=True) | (197, 1024) |
| **block_6** | **TransformerBlock(dim=1024, n_head=16, qkv_bias=True)** | **(197,1024)** |
| block_7 | TransformerBlock(dim=1024, n_head=16, qkv_bias=True) | (197, 1024) |
| block_8 | TransformerBlock(dim=1024, n_head=16, qkv_bias=True) | (197, 1024) |
| block_9 | TransformerBlock(dim=1024, n_head=16, qkv_bias=True) | (197, 1024) |
| block_10 | TransformerBlock(dim=1024, n_head=16, qkv_bias=True) | (197, 1024) |
| block_11 | TransformerBlock(dim=1024, n_head=16, qkv_bias=True) | (197, 1024) |
| **block_12** | **TransformerBlock(dim=1024, n_head=16, qkv_bias=True)** | **(197,1024)** |
| block_13 | TransformerBlock(dim=1024, n_head=16, qkv_bias=True) | (197, 1024) |
| block_14 | TransformerBlock(dim=1024, n_head=16, qkv_bias=True) | (197, 1024) |
| block_15 | TransformerBlock(dim=1024, n_head=16, qkv_bias=True) | (197, 1024) |
| block_16 | TransformerBlock(dim=1024, n_head=16, qkv_bias=True) | (197, 1024) |

| | | | |
|---|---|---|---|
| block_17 | TransformerBlock(dim=1024, n_head=16, qkv_bias=True) | (197, 1024) | |
| **block_18** | **TransformerBlock(dim=1024, n_head=16, qkv_bias=True)** | **(197,1024)** | |
| block_19 | TransformerBlock(dim=1024, n_head=16, qkv_bias=True) | (197, 1024) | |
| block_20 | TransformerBlock(dim=1024, n_head=16, qkv_bias=True) | (197, 1024) | |
| block_21 | TransformerBlock(dim=1024, n_head=16, qkv_bias=True) | (197, 1024) | |
| block_22 | TransformerBlock(dim=1024, n_head=16, qkv_bias=True) | (197, 1024) | |
| **blocks_end** | **TransformerBlock(dim=1024, n_head=16, qkv_bias=True)** | **(197,1024)** | |
| | LayerNorm(768, eps=1e-05, elementwise_affine=True) | (197,1024) | |
| | slice(0,:) | (1024) | |
| **final** | **Linear(1024, 1000, bias=True)** | **(1000)** | **1000** |

The TransformerBlock components of the architecture have the following structure:

| |
|---|
| 1. input |
| 2. LayerNorm(dim) |
| 3. Attention(dim, num_heads, qkv_bias) |
| 4. Residual: Add (1) to output of (3) |
| 5. LayerNorm(d_model) |
| 6. MLP Layer:<br>a. Linear (d_model, d_model * 4)<br>b. GELU<br>c. Linear (d_model * 4, d_model) |
| 7. Residual: Add output of (4) to output of (6) |

***LowpassAlexNet***

Modifications were made to the AlexNet architecture to reduce aliasing. The model consists of 5 convolutional layers, 5 weighted-average-pooling (HannPooling) layers, and 2 fully connected layers (plus one classification layer).

Model stage names and number of features are only given for the stages used in metamer experiments.

| Model Stage Name | PyTorch Operation | Output Shape | Number of Features |
|---|---|---|---|
| **natural_image** | **input** | **(3,224,224)** | **150528** |
| | Conv2d(3, 64, kernel_size=11, stride=1, padding=5) | (64, 224, 224) | |
| | ReLU | (64, 224, 224) | |
| | HannPooling2d(pool_size=17, stride=4, padding=5) | (64, 55, 55) | |
| **relu0** | **ReLU** | **(64, 55, 55)** | **193600** |
| | HannPooling2d(pool_size=9, stride=2, padding=2) | (64, 27, 27) | |
| | Conv2d(64, 192, kernel_size=5, padding=2) | (192, 27, 27) | |
| **relu1** | **ReLU** | **(192, 27, 27)** | **139968** |
| | HannPooling2d(pool_size=9, stride=2, padding=2) | (192, 13, 13) | |
| | Conv2d(192, 384, kernel_size=3, padding=1) | (384, 13, 13) | |
| **relu2** | **ReLU** | **(384, 13, 13)** | **64896** |
| | Conv2d(384, 256, kernel_size=3, padding=1) | (256, 13, 13) | |
| **relu3** | **ReLU** | **(256, 13, 13)** | **43264** |
| | Conv2d(256, 256, kernel_size=3, padding=1) | (256, 13, 13) | |
| **relu4** | **ReLU** | **(256, 13, 13)** | **43264** |
| | HannPooling2d (pool_size=9, stride=2, padding=2) | (256, 6, 6) | |
| | Dropout(p=0.5) | (9216) | |

| | | | |
|---|---|---|---|
| | Linear(256 * 6 * 6, 4096) | (4096) | |
| **fc0_relu** | **ReLU** | **(4096)** | **4096** |
| | Dropout(p=0.5) | (4096) | |
| | Linear(4096, 4096) | (4096) | |
| **fc1_relu** | **ReLU** | **(4096)** | **4096** |
| **final** | **Linear(4096, 1000)** | **(1000)** | **1000** |

***VOneAlexNet***

VOneAlexNet was proposed in (67)  and consists of 5 convolutional layers, 3 max-pooling layers, and 2 fully connected layers (plus one classification layer). Gaussian noise rather than Poisson-like noise was used during training, as proposed in (68).

Model stage names and number of features are only given for the stages used in metamer experiments.

| Model Stage Name | PyTorch Operation | Output Shape | Number of Features |
|---|---|---|---|
| **natural_image** | **input** | **(3,224,224)** | **150528** |
| | gabors(simple_channels=256, complex_channels=256, kernel_size=25, stride=4) | (512, 56, 56) | |
| **v1_output** | **additive_gaussian_noise(std=4)** | **(512, 56, 56)** | **1605632*** |
| | Conv2d(512, 64, kernel_size=1, stride=1, bias=False) | (64, 56, 56) | |
| | Conv2d(64, 192, kernel_size=5, stride=2, padding=2) | (192, 28, 28) | |
| **relu1** | **ReLU** | **(192, 28, 28)** | **150528** |
| | MaxPool2d(kernel_size=3, stride=2, padding=1) | (192, 14, 14) | |
| | Conv2d(192, 384, kernel_size=3, padding=1) | (384, 14, 14) | |
| **relu2** | **ReLU** | **(384, 14, 14)** | **75264** |
| | Conv2d(384, 256, kernel_size=3, padding=1) | (256, 14, 14) | |
| **relu3** | **ReLU** | **(256, 14, 14)** | **50176** |

| | | | |
|---|---|---|---|
| | Conv2d(256, 256, kernel_size=3, padding=1) | (256, 14, 14) | |
| **relu4** | **ReLU** | **(256, 14, 14)** | **50176** |
| | MaxPool2d(kernel_size=3, stride=2, padding=1) | (256, 7, 7) | |
| | Dropout(p=0.5) | (12544) | |
| | Linear(256 * 6 * 6, 4096) | (4096) | |
| **fc0_relu** | **ReLU** | **(4096)** | **4096** |
| | Dropout(p=0.5) | (4096) | |
| | Linear(4096, 4096) | (4096) | |
| **fc1_relu** | **ReLU** | **(4096)** | **4096** |
| **final** | **Linear(4096, 1000)** | **(1000)** | **1000** |

*Metamers were generated from the output of the VOneNet block. We note that this model stage has more parameters than the comparison relu0 stage of AlexNet and LowpassAlexNet, but that we plot the relu0/VOneBlock on the same line in Figure 6d and Supplementary Figure 13. Subsequent stages (relu1 onwards) have similar dimensionality, although differ slightly due to the padding in the VOneAlexNet architecture.

### Auditory Models

#### *CochResNet50*

The CochResNet50 model is a ResNet50 backbone architecture applied to a cochleagram representation (such that 2D convolutions learned on the cochleagram).

| Model Stage Name | PyTorch Operation | Output Shape | Number of Features |
|---|---|---|---|
| **natural_audio** | **Input (waveform)** | **(40000)** | **40000** |
| **cochleagram** | **Cochleagram** | **(1,211,390)** | **82290** |
| | Conv2d(1, 64, kernel_size=7, stride=2, padding=3, bias=False) | (64, 106, 195) | |
| | BatchNorm2d(64) | (64, 106, 195) | |
| **conv1_relu1** | **ReLU** | **(64, 106, 195)** | **1322880** |
| | MaxPool2d(kernel_size=3, stride=2, padding=1) | (64, 53, 98) | |

| | | | |
|---|---|---|---|
| **layer1** | **ResNetBlock(inplanes=64, planes=64, num_blocks=3, stride=1)** | **(256, 53, 98)** | **1329664** |
| **layer2** | **ResNetBlock(inplanes=256, planes=128, num_blocks=4, stride=2)** | **(512, 27, 49)** | **677376** |
| **layer3** | **ResNetBlock(inplanes=512, planes=256, num_blocks=6, stride=2)** | **(1024, 14, 25)** | **358400** |
| **layer4** | **ResNetBlock(inplanes=1024, planes=512, num_blocks=3, stride=2)** | **(2048, 7, 13)** | **186368** |
| **avgpool** | **AdaptiveAvgPool2d(1,1)** | **(2048, 1, 1)** | **2048** |
| **final** | **Linear(2048, 794)** | **(794)** | **794** |

The ResNetBlock components of the architecture have the following structure (same as in ResNet50 visual model):

| |
|---|
| 1. input (x) |
| 2. 1x1 Conv2d(inplanes, planes, stride=1) |
| 3. BatchNorm2d(planes) |
| 4. ReLU |
| 5. 3x3 Conv2d (planes, planes, stride=1) |
| 6. BatchNorm2d(planes) |
| 7. ReLU |
| 8. 1x1 Conv2d (planes, planes * expansion, stride=1) |
| 9. BatchNorm2d(planes) |
| 10. Residual connection on x (if inplanes !=planes * expansion): 1x1 Conv2D (inplanes, planes * expansion, stride) |
| 11. Residual connection on x (if inplanes !=planes * expansion): BatchNorm2d(planes * expansion) |
| 12. Add output from (9) to output from (11) |
| 13. (Output) ReLU |

Multiple of these residual blocks (num_blocks) are stacked together to form a single ResNetBlock. The expansion factor was set to four for all layers (expansion=4).

***CochCNN9***

The CochCNN9 architecture is based on found in (5) through a neural network architecture search. The architecture differs in that the input to the first stage of the model is not reshaped to 256x256, rather it is maintained as the 211x390 size cochleagram. The convolutional layer filters and pooling regions are similarly reshaped to maintain the approximate receptive field size in frequency and time. The model is also trained with batch normalization rather than the local response normalization used in (5).

| Model Stage Name | PyTorch Operation | Output Shape | Number of Features |
|---|---|---:|---:|
| **natural_audio** | **Input (waveform)** | **(40000)** | **40000** |
| cochleagram | Cochleagram | (1,211,390) | **82290** |
| | BatchNorm2d(1) | (1,211,390) | |
| | Conv2d(1, 96, kernel_size = [7, 14], stride = [3, 3], padding = 'same') | (96, 71, 130) | |
| **relu0** | **ReLU** | **(96, 71, 130)** | **886080** |
| | MaxPool2d(kernel_size = [2,5] , stride = [2,2], padding = 'same') | (96, 36, 65) | |
| | BatchNorm2d(96) | (96, 36, 65) | |
| | Conv2d(96, 256, kernel_size = [4,8], stride = [2,2], padding = 'same') | (256, 18, 33) | |
| **relu1** | **ReLU** | **(256, 18, 33)** | **152064** |
| | MaxPool2d(kernel_size = [2,5] , stride = [2,2], padding = 'same') | (256, 9, 17) | |
| | BatchNorm2d(256) | (256, 9, 17) | |
| | Conv2d(256, 512, kernel_size = [2,5], stride = [1,1], padding = 'same') | (512, 9, 17) | |
| **relu2** | **ReLU** | **(512, 9, 17)** | **78336** |
| | Conv2d(512, 1024, kernel_size = [2,5], stride = [1,1], padding = 'same') | (1024, 9, 17) | |
| **relu3** | **ReLU** | **(1024, 9, 17)** | **156672** |
| | Conv2d(1024, 512, kernel_size = [2,5], stride = [1,1], padding = 'same') | (512, 9, 17) | |
| **relu4** | **ReLU** | **(512, 9, 17)** | **78336** |

| | | | |
|---|---|---|---|
| **avgpool** | **AvgPool(kernel_size = [2,5] , stride = [2,2], padding = 'same')** | **(512, 5, 9)** | **23040** |
| | Linear(512*9*5 , 4096) | (4096) | |
| **relufc** | **ReLU** | **(4096)** | **4096** |
| | Dropout(p=0.5) | (4096) | |
| **final** | **Linear(4096, 794)** | **(794)** | **794** |
